# *De novo* design of transmembrane β-barrels

**DOI:** 10.1101/2020.10.22.346965

**Authors:** Anastassia A. Vorobieva, Paul White, Binyong Liang, Jim E Horne, Asim K. Bera, Cameron M. Chow, Stacey Gerben, Sinduja Marx, Alex Kang, Alyssa Q. Stiving, Sophie R. Harvey, Dagan C. Marx, G. Nasir Khan, Karen G. Fleming, Vicki H. Wysocki, David J. Brockwell, Lukas K. Tamm, Sheena E. Radford, David Baker

**Affiliations:** Department of Biochemistry, University of Washington, Seattle, WA 98195, USA; Howard Hughes Medical Institute, University of Washington, Seattle, WA 98195, USA; Astbury Centre for Structural Molecular Biology, School of Molecular and Cellular Biology, Faculty of Biological Sciences, University of Leeds, Leeds LS2 9JT; Department of Molecular Physiology and Biological Physics and Center for Membrane and Cell Physiology, University of Virginia, Charlottesville, VA 22903, USA; Institute for Protein Design, University of Washington, Seattle, WA 98195, USA; Department of Molecular Engineering and Sciences, University of Washington, Seattle, WA 98195, USA; Department of Chemistry and Biochemistry, Resource for Native Mass Spectrometry Guided Structural Biology, The Ohio State University, Columbus, OH 43210; TC Jenkins Department of Biophysics Johns Hopkins University, Baltimore, MD 21218

## Abstract

The ability of naturally occurring transmembrane β-barrel proteins (TMBs) to spontaneously insert into lipid bilayers and form stable transmembrane pores is a remarkable feat of protein evolution and has been exploited in biotechnology for applications ranging from single molecule DNA and protein sequencing to biomimetic filtration membranes. Because it has not been possible to design TMBs from first principles, these efforts have relied on re-engineering of naturally occurring TMBs that generally have a biological function very different from that desired. Here we leverage the power of *de novo* computational design coupled with a “hypothesis, design and test” approach to determine principles underlying TMB structure and folding, and find that, unlike almost all other classes of protein, locally destabilizing sequences in both the β-turns and β-strands facilitate TMB expression and global folding by modulating the kinetics of folding and the competition between soluble misfolding and proper folding into the lipid bilayer. We use these principles to design new eight stranded TMBs with sequences unrelated to any known TMB and show that they insert and fold into detergent micelles and synthetic lipid membranes. The designed proteins fold more rapidly and reversibly in lipid membranes than the TMB domain of the model native protein OmpA, and high resolution NMR and X-ray crystal structures of one of the designs are very close to the computational model. The ability to design TMBs from first principles opens the door to custom design of TMBs for biotechnology and demonstrates the value of *de novo* design to investigate basic protein folding problems that are otherwise hidden by evolutionary history.

**One sentence summary:** Success in *de novo* design of transmembrane β-barrels reveals geometric and sequence constraints on the fold and paves the way to design of custom pores for sequencing and other single-molecule analytical applications.

## Main text

Advances in *de novo* protein design have yielded water soluble proteins of increasing complexity (*1–5*), and several examples of α-helical membrane proteins (*6, 7*). However, the *de novo* design of an integral transmembrane β-barrel (TMB) has not yet been achieved. TMBs can spontaneously fold into lipid bilayers from an unfolded chain, possibly through a mechanism involving concerted membrane insertion and folding of the β-hairpins (*8, 9*). How this folding in a non-aqueous environment is encoded in the sequences of TMBs is not well understood because of experimental challenges in characterizing the rugged folding pathway - including possible off-pathway, misfolded or “invisible” states (*10*) - and the often non-superimposable folding and unfolding equilibria (hysteresis) (*11*). To prevent misfolding and aggregation of the TMB polypeptide *in vivo*, an array of chaperones assist TMB biogenesis, maintaining the polypeptide chain in a folding-competent state and transporting it to the relevant biological membrane – the outer membranes of prokaryotes, mitochondria and chloroplasts (*12*). Because of this complexity, the lipid-folding/water-aggregation trade-off places poorly understood constraints on the global sequence properties of TMBs, and the relatively small amount of biochemical and structural data have slowed down the development of computational design methods. Yet, engineering of naturally occurring TMBs have provided proteins with new and useful functions, such as nanopores for single molecule DNA sequencing (*13*), small-molecule sensing (*14, 15*) or water filtering bioinspired membranes (*16*).

To shed light on the sequence determinants of folding and stability of TMBs, and to enable the custom design of TMBs for specific applications, we set out to design TMBs *de novo*. We started by studying the constraints membrane embedding puts on both the backbone geometry and sequence of β-barrels.

### Geometric constraints on transmembrane β-barrel backbones

TMBs are formed from a single β-sheet that twists and bends to close on itself, so that all membrane-embedded backbone polar groups are hydrogen-bonded and shielded from the lipid environment. Insertion of TMBs into the lipid membrane is oriented (*17*), with β-strands usually connected with long loops on the translocating (*trans*) side of the β-barrel (extracellular in bacteria) and short β-turns on the non translocating (*cis*) (Fig. S1A). The β-barrel architecture is characterized by two discrete parameters: the number of strands (n) and the shear number (S)--the shift in the number of residues (register shift) along a strand after tracing around the barrel through the backbone hydrogen bonds (*18*). The ideal β-barrel radius r (eq. 1) and angle of the strands with the main barrel axis θ (eq. 2) are functions of n, S, the average distance between two β-strands (D) and the average distance between two residues on a β-strand (d) (Table S1) (*19*).

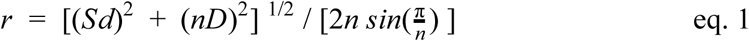

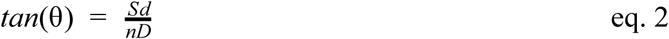

The shear number (S) and the number of strands (n) also define the packing arrangement of the stripes of Cβs packing along the interstrand hydrogen bonds (half of the Cβ-stripes point toward the β-barrel lumen and the other half toward the β-barrel exterior) (Fig. S1B).

We chose to focus on the simplest and smallest β-barrel architecture of 8 β-strands. We first considered a shear number of 8 (n==S). In such a configuration, the total register shift is distributed equally among the four β-hairpins (2 residues per hairpin) and the side-chains pointing toward the lumen of the barrel are arranged into 4-fold symmetric Cβ-stripes (Fig S1C). We found that such a symmetric packing arrangement combined with a small β-barrel radius does not allow tight jigsaw-puzzle like packing in the core as the C㬚-Cβ vectors at each rung of the barrel point at each other. We chose to break the symmetry in the core by designing β-barrels with a shear number of 10; in this case the C㬚-Cβ vectors are arranged into 5 intertwined Cβ-strips which spiral around the barrel axis so different side-chains are at different heights and more uniform packing can be achieved (Fig. S1D). To do so, we increased the register shift between two β-strands from 2 to 4 residues, which increases the barrel radius (eq. 1) and the angle of the β-strands with the barrel axis (eq. 2).

The uneven distribution of register shifts between hairpins complicates interactions with the lipid membrane. The bilayer can be approximated as two planes that must be parallel to ensure constant membrane thickness. In natural TMBs the *cis* (periplasmic) β-turns are close to the periplasmic lipid/water boundary (Fig. S2A-D). While the β-turn residues closely match the sequence preferences observed in water-soluble β-barrels (mostly polar residues), the surface-exposed residues flanking these β-turns are predominantly hydrophobic (Fig. S2H-K) (*20*). We postulated that the hydrophobic residues upstream of the β-turns define the *cis* boundary of the transmembrane region because of their lowest position on the staggered hairpins (“membrane anchor residues”, Fig. S2A-D). The geometric challenge is that the the plane representing the *cis* membrane boundary must be aligned with the position of these four anchor residues in 3D space. Whereas the symmetric n=S=8 barrel has flat rungs which can be readily aligned with the membrane planes, the n=8 S=10 barrel does not. The total change of level (Z) between the lowest and the highest of the 4 anchor residues along the main β-barrel axis (eq. 5) is the sum of the difference in vertical offset along the β-strands (eq. 3, where *a* is the register shift) and along the hydrogen bonds (eq. 4, where *b*=2 is the number of strands between each anchor residue) (eq. 5, Fig. S3A).

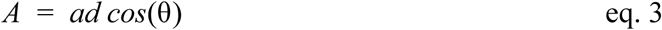

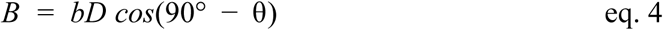

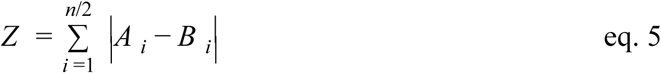

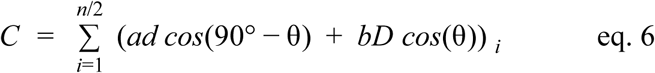

This vertical offset can in principle be accommodated by tilting the β-barrel by an angle *α* = arctan (Z/C) where the denominator is the length of the arc between anchor residues 1 to 4 projected onto the plane perpendicular to the main axis (eq. 6) (Fig. 1A, Fig. S3B). In the case of a β-barrel with symmetry (n=8,S=8), the vertical offset between each anchor residue is negligible (0.14 Å) and no tilt is required. When the shear number is increased to 10 by increasing the register shift between one pair of hairpins to 4 residues, the total vertical offset through the β-sheet is 3.9 Å over an arc length of 33.3 Å, and the barrel must be tilted by approximately 6.7° to the transmembrane axis (Fig. 1C, top). When the total 10 residue register shift is achieved instead by assigning a 6 residue register shift to one pair of hairpins, and zero shift to a second pair, eq. 5 and 6 predict a total vertical offset of 8.8 Å over 28.8 Å, and hence a more pronounced tilt angle *α* of approximately 16.9° (Fig. S4A). To validate this geometric model, we assembled sequence-agnostic protein backbones with the Rosetta fragment assembly protocol (*21*), designed the lipid-exposed surface, and predicted the lipid membrane position with the PPM server (*22*). The average predicted tilt angles of the barrel to the transmembrane axis are close to the predictions for each of the register shift distributions considered above (8.1° (Fig. 1C, top) and 16°, respectively (Fig. S4A,G)). We decided to continue our design efforts with the less tilted configuration, because it had a better match to the desired hydrophobic thickness of the membrane (24.3 Å+/−0.6) than the more tilted configuration (23.2 Å+/−1) (Fig. S4E) and had a more negative transfer energy from water to lipid (−38 kcal/mol vs -34 kcal/mol; predicted with the PPM server) (Fig. S4F)). Placing the four-residue register shift after any of the four *cis* hairpins resulted in structures with similar average hydrophobic thicknesses, tilt angle to the membrane axis and transfer energy from water to lipid and only differed on the direction of the tilt (Fig. S4B-G); we chose to focus on one of these placements in which the 4-residue register shift is in the middle of the β-sheet.

**Fig. 1.**
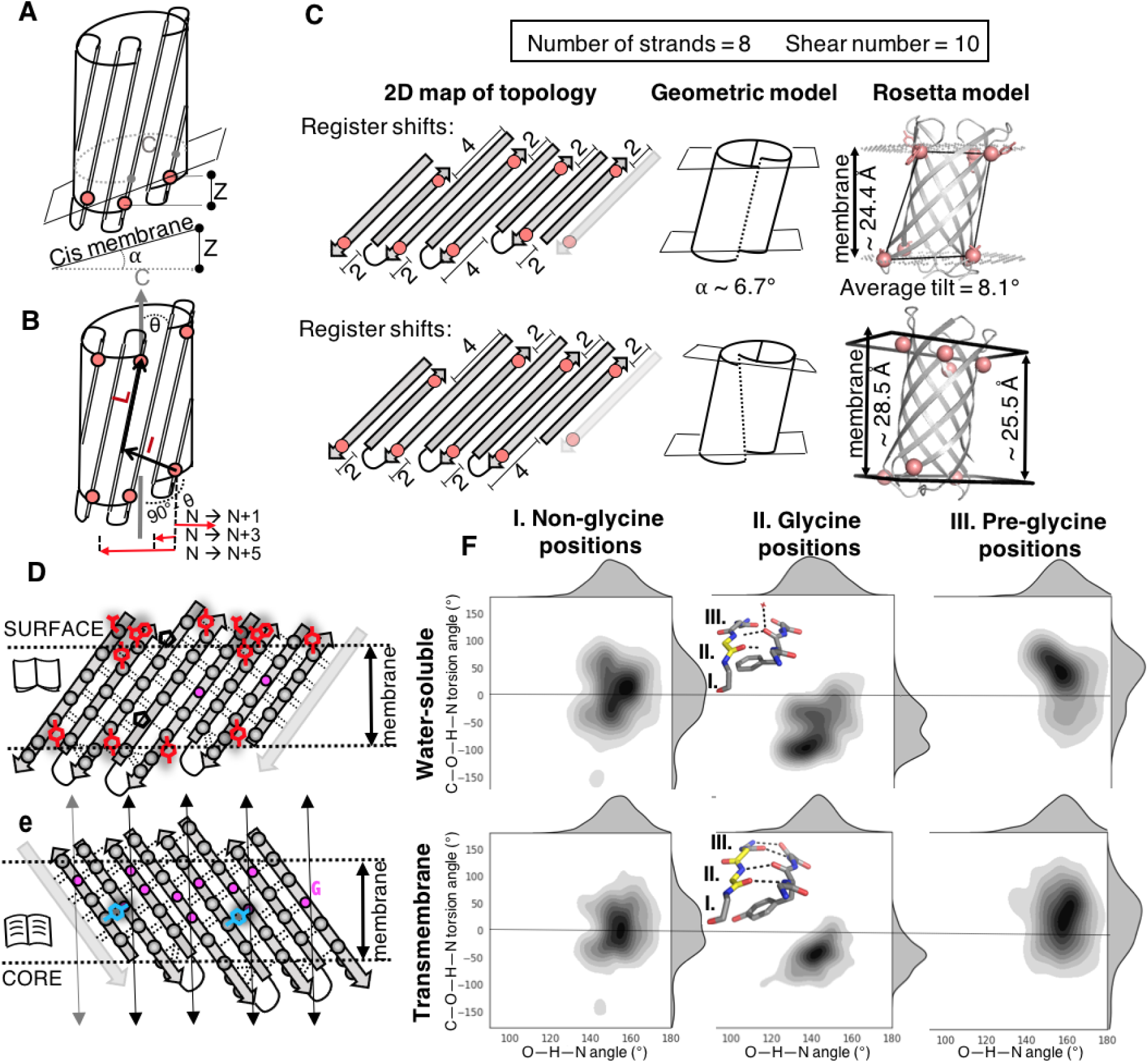
Principles for designing TMBs backbones. In panels A-D, the membrane anchoring residues are shown as salmon spheres. (A, B) Geometric model of membrane-association constraints on the β-barrel architecture. (A) Asymmetric register shifts between the β-hairpins can be accommodated by tilting the β-barrel to the transmembrane axis by an angle *α* = arctan(Z/C). (B) To place asymmetric register shifts on the *trans* and *cis* membrane boundaries, the distances between the *cis* anchor residue N and all anchor residues in *trans* were calculated and projected to the horizontal plane. θ is the angle of the β-strands to the main β-barrel axis. (C) The geometric model (center) and Rosetta modelling followed by hydrophobic thickness prediction with PPM (right) predict similar tilt angles (*α*) of the β-barrel to the membrane axis for a given β-strand arrangement: both models show inconsistent hydrophobic thickness for β-barrel architectures with double register shifts are located on strand 1 in cis (strand N) (Top) and on strand 6 in trans (strand N+5)(Bottom). (D and E) 2D schematic representation of the connectivity (hydrogen bonds as dashed lines) between β-strands in the TMB designs. Side-chains are shown as grey spheres and glycine residues as pink dots. (D) Aromatic girdle motifs on the surface of the β-barrel are shown as red side chains. Prolines are shown as black pentagons. (F) The tyrosines of the mortise/tenon motifs are shown as blue side-chains. Glycine kinks were arranged to bend the β-sheet into four corners (vertical arrows). (F) Comparison of the backbone hydrogen bond geometries in water soluble (top) and transmembrane (bottom) β-barrels with 8 β-strands; data for glycine kink residues (center, residue II.), residues preceding a glycine kink (right, residue III.) and other residues (left, residue I.) are plotted separately. An example of glycine kink (residue II.) with an aromatic rescue interaction is shown in the central panel, with glycine residues colored in yellow and water molecules shown as red dots.

We next investigated the structural consequences of the fact that the *cis* and *trans* planes representing the membrane boundary must be roughly parallel to each other to keep the thickness of the membrane constant. We reasoned that the two planes could only be kept parallel if the offset Z for any hairpin on the *cis* face is matched by a similar offset for the hairpin directly above it on the *trans* face (Fig. 1B). We determined the projection of the vector between a *cis* anchor residue and all four *trans* anchor residues for a β-barrel spanning a membrane of 24 Å (eq. 3) on the plane perpendicular to the main barrel axis; we consider the *cis* and *trans* anchor residue pairs with the smallest projected distance to stack on each other along the barrel axis. For barrel topologies of (n=8,S=10), we found that the stacking partner for an anchor residue on the *cis* side of strand N is the anchor residue on the *trans* side of strand N+3. Hence, to maintain constant thickness, the offset Z between strands N and N+1 on the *cis* side must be equal to the offset between strands N+3 and N+4 on the *trans* side. To confirm this prediction of our geometric model, we set the *cis* side register shift between strands N and N+1 to four residues, and ran Rosetta design simulations and transmembrane plane predictions on backbones with a matching 4-residue register shift on the *trans* side between (i) strands N+3 and N+4 and (ii) strands N+5 and N+6. We averaged planes representing the membrane boundary in *cis* and *trans* and found, consistent with the model, parallel planes and constant hydrophobic thickness for the four residue register shift in N+3, but a 3Å change in membrane thickness in the N+5 case (Fig. 1C, bottom).

### Sequence design on ideal β-barrel backbones

We used this constant hydrophobic thickness constraint to guide the distribution of the register shifts around the β-barrel. The *cis* hairpins were closed with short β-turns associated with an upstream β-bulge (abundant in water-soluble and transmembrane β-barrels (Fig. S2A)). Canonical β-turn sequences with strong β-hairpin nucleating properties (3:5 type I β-turns + G1 bulge with canonical SDG sequence (*23–25*)) were used to connect the strands on the *trans* side in place of the long loops found in native TMBs; such turns were previously used to design water-soluble β-barrels (Fig. S2E,G) (*5*). To relieve strain from high β-sheet curvature, we placed glycine kinks (*5*) - glycine residues with an extended β-sheet backbone conformation - into the TMB backbone description (the backbone “blueprint”) such that a) every Cβ-strip pointing to the core of the barrel contains a glycine and b) there are no more than 4 non-glycine residues in a row in the Cβ-strips (¼ of the average barrel circumference). Following these principles, we designed a β-barrel blueprint in which the glycine kinks in the core of the protein were stacked along four vertical lines together with β-bulges associated with the *cis* hairpins (Fig. 1D,E). Rosetta models built from the above blueprint have four regions of strong β-sheet bending surrounding a wide β-barrel lumen (Fig. S2F).

To delimit the upper and lower membrane boundaries, four tyrosine residues were placed two positions upstream of the anchor residues on the *cis* side, and alternating tyrosine and tyrosine/tryptophan motifs were placed at the *trans* boundary (Fig. 1D; such “aromatic girdles” are observed in native TMBs (*26*)). To design the remainder of the sequence, we first considered the possibility that the core residues could be largely non-polar (helical transmembrane proteins have been designed by keeping the core of soluble designs fixed and resurfacing the outside with hydrophobic residues). However, this approach was rapidly dismissed as the resulting sequences had very strong amyloid propensity (Fig. S5). Since native TMBs are usually less hydrophobic than α-helical membrane proteins (*27*), we next experimented with requiring the interior of the barrel to be polar to achieve the classic alternating hydrophobic-polar sequence pattern of canonical β-strands (inside/out model). We restricted the core positions to polar amino acids (excluding the glycine kink positions), increased the weight on the Rosetta electrostatic potential to favor sidechain-sidechain hydrogen bonds, and restricted the surface to hydrophobic amino acids. To help define the register between β-strands we placed tyrosine residues adopting the +60,90 rotamer angles to closely interact with the grove formed by a hydrogen-bonded glycine kink partner and donating a hydrogen bond to a negatively charged residue (*28*) (this is an extended definition of the mortise/tenon motif (*29, 30*)). Two positions were considered, the area of the sharp change of level between anchor residues (4-residues register shift) on the *cis* and *trans* faces, and the other side of the barrel between the first and last strands (Fig. 1E, Fig. S6). Finally, we designed full β-barrel sequences using Rosetta combinatorial sequence design guided by these principles and found, as expected, that the secondary structure was accurately recapitulated by secondary structure prediction programs (Fig. 2A).

**Fig. 2.**
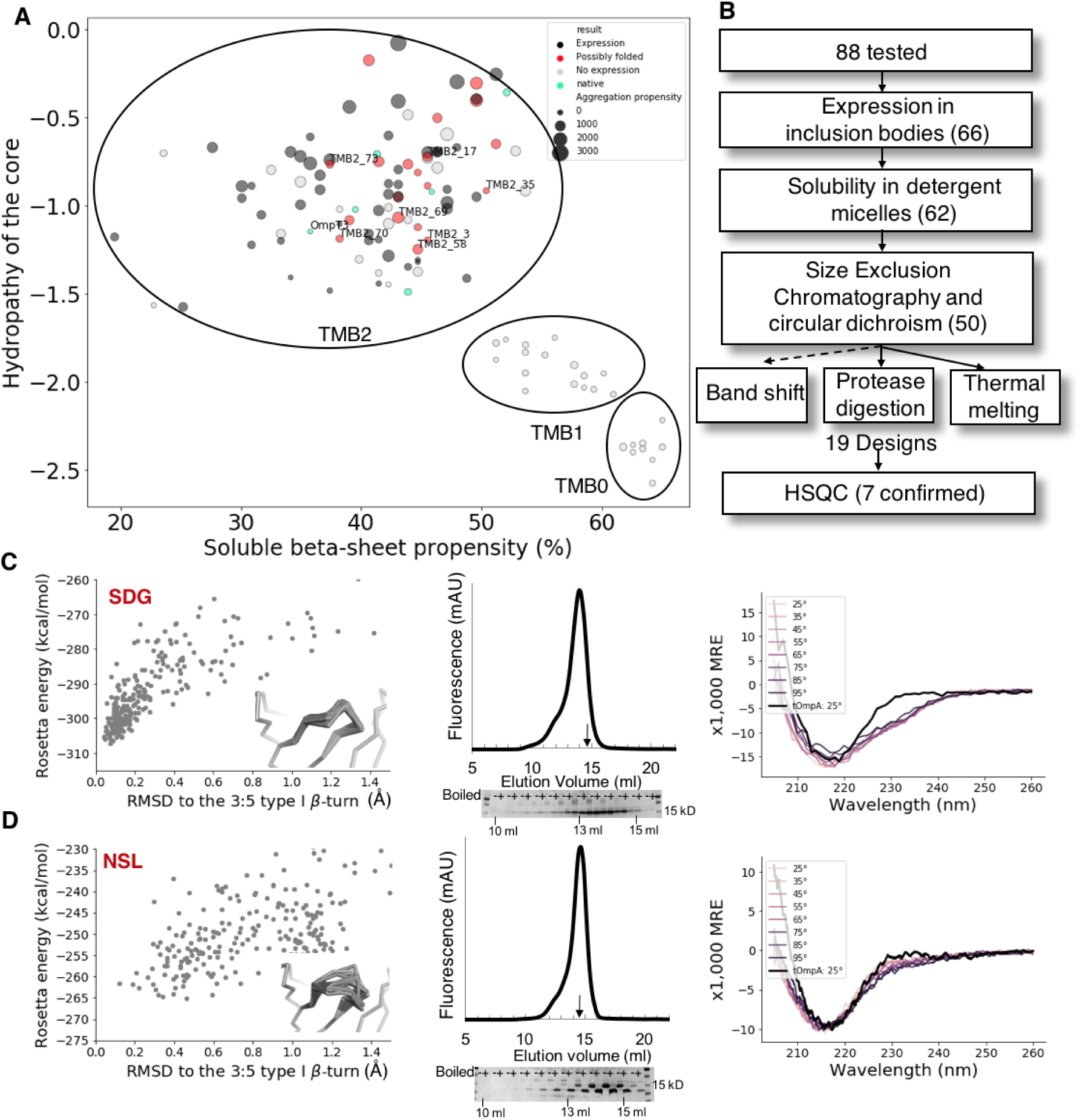
Negative design is critical for de novo TMB folding. (A) Successful design of TMBs requires reducing β-sheet propensity. X axis: β-sheet propensity (calculated with RaptroX (*62*), y axis: hydrophobicity of the core (GRAVY hydropathy index (*63*)). Grey spheres, Non-expressing TMB designs; black circles, expressing designs that do not fold; red, TMB designs that pass biochemical folding screening -- labels indicate folded species was validated by HSQC; Greenk, naturally occurring TMBs with 8 strands. Circle size, aggregation propensity of the sequence predicted with TANGO (*64*). (B) Experimental workflow. The number of unique designs (excluding loop doublons) satisfying each criteria is shown in brackets. (C and D) Proper folding of tOmpA requires negative design against strong β-turn nucleating sequences on the *trans* side. Left: Rosetta energy landscapes of designs with canonical low energy (C) or sub-optimal (D) sequences substituted in a 3:5 type I β-turn with a G1 β-bulge. Conformational perturbations were generated using Kinematic Loop Closure (*65*); the inset shows the backbone conformations of the twenty-five lowest-energy models. Center: After refolding in 2X CMC DDM detergent, OmpTrans3 elutes on SEC similarly to tOmpA (arrow, 14.62 ml for OmpTrans3 and 14.53 ml for tOmpA) and runs as a heat modifiable species on SDS-PAGE characteristic of folded tOmpA, while the OmpAAG peak elutes earlier (13.96 ml) and does not show a band shift. Right: The far-UV CD spectrum of OmpTrans3, but not OmpAAG, is similar to that of tOmpA.

Folding of TMBs is chaperone-mediated and catalyzed *in vivo* (by the β-barrel assembly machinery (BAM) complex in Gram-negative bacteria, the sorting and assembly machinery (SAM) complex in mitochondria, and the translocase of the outer chloroplast membrane (TOC) complex in chloroplasts) (*31*). Since it was unclear whether our TMB designs would be able to interact with the chaperone machinery to fold in the outer membrane of *E. coli*, we chose to express them in the cytoplasm, with the anticipation that the expressed sequences would form inclusion bodies that could then be solubilized in urea/guanidinium chloride (both natural and engineered TMBs have been produced in this way (*32*)). We obtained *E. coli* codon optimized synthetic genes for 9 designs (set TMB0, Fig. S7), but no protein of the correct molecular weight was produced upon the induction of protein expression (Data S2). Reasoning that the designed sequences may have had too much positive charge (*33*), in a second round of 16 designs, we reduced the number of charged residues in the core of the protein (set TMB1, Fig. S7), but again none expressed in *E coli* (Data S2).

These failures in expressing our TMB designs in *E. coli* were challenging because it was difficult to get feedback to improve the design methodology. To make progress, we took a step back and compared our designs to sequences of natural 8-strand TMBs. We noted two differences: first, the natural TMBs often have at least one of the *trans* loops disordered and greater than 20 residues in length (*20*), and second, the secondary structure propensity of the natural TMBs was lower than the designs we tried to express (Fig. 2A). We hypothesized that the strong secondary structure propensity of our designed sequences could result in folding of non-designed soluble β-sheet structures when expressed in the cytoplasm - possibly amyloid-like intermediates (*34*) - which could be toxic and hence cleared rapidly and/or hindering growth of expressing cells.

We first considered the possibility that the long disordered loops in *trans* might be necessary to slow down the non-native folding in the cytoplasm. To test this hypothesis, we obtained synthetic genes encoding 4 of the TMB0 designs with the extracellular loops replaced with either the extracellular loops of the native TMB domain of Outer Membrane Protein A of *E. coli* (tOmpA) or scrambled versions of these loops, as well as a redesigned version of tOmpA in which its *trans* loops were replaced with the 3-residues 3:5 type I β-turns used in our designs (with the canonical sequence SDG, Fig. S8A,B). The re-looped tOmpA construct (OmpSDG) expressed at high levels in *E. coli* (where it was found in inclusion bodies), however, only two of the *de novo* designs with long loops showed expression but at very low levels which were insufficient for further characterization (Fig. S9). These results suggest that the protein expression failure is largely determined by the transmembrane β-strands rather than by their β-hairpin connections. Further characterization showed that the OmpSDG protein, while highly expressed, was not correctly folded: its circular dichroism (CD) spectrum, particularly in the 230 nm region (Fig. 2C), was different from native tOmpA when refolded in n-dodecyl-β-D-maltopyranoside (DDM) detergent at 2 times the critical micelle concentration (CMC), and it did not show the heat-modifiable band shift on SDS-PAGE characteristic of folded tOmpA (*35*) in DDM detergent and when refolded in large unilamellar vesicles (LUVs) of 1,2-diundecanoyl-sn-glycero-3-phosphocholine (DUPC, *di*C_11:0_PC) (Fig. 2C, Fig. S10C,D).

To understand the failure of OmpSDG to fold, we searched the PDB for short β-turns at the *trans* membrane boundary of natural TMB PDB structures, which are rare. We found five 3-residue *trans* β-turns whose backbone conformation and hydrogen bonding pattern satisfied all the characteristics of the canonical 3:5 type I β-turn with a G1 β-bulge (Data S3). However, the sequences of these β-turns are suboptimal for their structure compared to the SDG canonical sequence (*36*), as shown by the structure/energy landscapes computed with Rosetta for each of these turns (Fig. 2C, Fig. S8D). Further evidence of different properties for *trans* and *cis* β-turns despite identical backbone conformations in crystal structures is that small protein fragments retreived from the PDB by matching sequences found in *cis* showed β-turn-like structural properties, while queries matching sequencies of *trans* β-turns did not show any structural convergence (Fig. S11). We tested whether this observation could be generalized by comparing sets of *trans* and *cis* β-turns of two to five residues and found worse predicted sequence-structure compatibility (Rosetta p_aa_pp score) in *trans* turns (Fig. S8C). We hypothesized that, much like the long loops of native tOmpA, short non-canonical sequences could slow down nucleation of the *trans* β-hairpins. Accordingly, we tested 4 variants of tOmpA (Omp*Trans*1-4) that each contain two 3:5 type I β-turns with suboptimal sequences (these designs are shorter than the shortest variant of tOmpA previously reported, which has *trans* connections of 5 to 18 residues (*37*)). The proteins were again expressed at high levels in inclusion bodies (Table S2), but this time all four of these sequences showed a heat-modifiable band in DDM detergent micelles and LUVs characteristic of a folded TMB (Fig. 2D, Fig. S10C,D). We selected one of the variants - Omp*Trans*3 - that appeared to be produced in the largest amounts for further characterization. Omp*Trans*3 refolded in detergent micelles had a similar retention time to native tOmpA on a Size Exclusion Chromatography (SEC) column (Fig. 2D, Fig. S12), a similar native mass spectrometry (nMS) profile (Fig. S13), well-dispersed resonance peaks by H^1^-N^15^-HSQC NMR in Fos-choline-12 (DPC) detergent (Fig. S10B) and a similar CD spectrum to tOmpA in DDM detergent (Fig. 2D) and in LUVs with the distinctive 231 nm peak (Fig. S10A) (*38*). These data support the idea that slowed folding due to the presence of long or short suboptimal β-hairpin connection sequences on the *trans* side are necessary for proper folding of TMBs *in vitro*.

Guided by these results, we used the suboptimal β-turns we had inserted into the Omp*Trans*3 design in all of subsequent TMB *de novo* designs. To address the expression problem, we hypothesized that the culprit was the relatively high secondary structure propensity of the β-strands, and sought to address this by (i) increasing the hydrophobicity of the β-barrel lumen and thereby disrupting the strict alternation of polar and hydrophobic residues along the β-strands and (ii) introducing glycines in specific positions on the lipid-exposed surface. We experimented with extending the tyrosine-glycine motifs to include a negative charged Asp or Glu hydrogen bond acceptor to the tyrosine, using the Rosetta HBNet protocol (*39*) to exhaustively search through all the possible positions. We kept such YGD/E networks fixed, and used Rosetta combinatorial sequence optimization to design the remainder of the sequence. We allowed all 18 amino acids other than Cys and Pro in positions facing the core of the barrel, and hydrophobic amino acids only on the lipid-exposed surface. The models were selected based on protein backbone quality (backbone torsion angles and hydrogen bonds) and the quality of the networks around each YGD/E motif (hydrogen bond potential, size, connectivity and robustness of the networks).

We compared the designed surface residue composition to that of native transmembrane barrels, and found that glycine (which destabilizes β-strands (*40*)), while very rare in the corresponding region of water soluble β-barrels (we found only four such examples - three were buried in the midst of dimerization interfaces (Supplementary text)) and disallowed in the above designs, represents 6.3% of all amino acids on the lipid exposed surface of natural 8-strands TMBs (Fig. S7D) (*41*). These surface glycine residues of TMBs often precede glycine kinks hydrogen-bonded with core polar network hot spots (such as the tyrosines in the mortise/tenon motifs) or are located between two glycine kinks. We inspected crystal structures of water-soluble and transmembrane β-barrels and found that, while most β-strand residues have canonical in-plane backbone hydrogen bonds (O--H--N angle ∼ 160°; C--O--H--N dihedral ∼ 0°) (*42*) and canonical ɸ and Ѱ torsion angles (Fig. 1F, left, Fig. S12C,D), glycine kinks have more extended backbone conformation (positive ɸ and/or negative Ѱ torsion angles (Fig. S14A,B)). In water-soluble β-barrels, glycine kinks also have out-of-plane hydrogen bonds geometries characteristic of a left-hand twist (O--H--N angle ∼ 130°; C--O--H--N dihedral ∼ -100°, Fig. 1F), while the surface residues preceding the glycine kink have more pronounced right-hand twist (C--O--H--N dihedral > 0°, Fig. 1F, Fig. S14G). Many backbone carbonyls in these out-of-plane hydrogen bond geometries were found to interact with a secondary hydrogen bond donor in the crystal structures (such as a water molecule or a surface residue side chain, Fig. S14E,F), but such hydrogen bonds would likely be disfavored in TMBs, in the absence of water available to interact with the exposed carbonyls. Indeed, TMBs have a smaller population of glycine kinks and pre-glycine hydrogen bonds significantly deviating from in-plane geometry (Fig. 1F). We hypothesized that glycines in positions preceding glycine kinks could allow more canonical hydrogen bonds by relieving backbone strain. We carried out a further round of design of the surface-exposed residues allowing glycine and increasing the weight on the Rosetta long range hydrogen bond energy (which favors in-plane geometries). All resulting designs had two to three surface glycines (in average 5.6% of the surface amino acids). Two of the glycines were common to all designs (G26 associated with a mortise/tenon motif and G56 between glycine kinks G55 and G57).

After three iterations between core and surface design, the design calculations converged on roughly 30 distinct network architectures with overall amino acid composition similar to that of natural 8-strands TMBs (Fig. S7D). Codon optimized synthetic genes were obtained for several representatives of each core network architecture for a total of 90 designs (set TMB2). We also expressed 20 additional variants of these designs incorporating the extracellular loops of tOmpA to evaluate the effect of loop length on folding. One hundred and eight of these designs were tested, 80 were well expressed and were found exclusively in inclusion bodies (or 66 unique designs, excluding *trans* loop doublons). Notably, the same designs expressed poorly or did not express at all either with short ideal β-turns or long loops (Data S2). The relatively high success rate in obtaining protein expression is in striking contrast to the earlier iterations described above in which no expression was observed.

### Characterization of folding, stability and structure

To test the ability of the designs to stably fold to TMB structures *in vitro*, we followed procedures used to fold tOmpA and other natural TMBs (Fig. 2B) (*43, 44*). Briefly, the inclusion bodies were dissolved in 8M urea and rapidly diluted into DDM, DPC or n-octyl-β-D-glucopyranoside (OG) detergents at 2X CMC (Data S4). Out of the sixty-six expressed unique designs, sixty-two formed soluble species in such conditions. We purified the protein/detergent complexes by SEC and characterized the fifty designs which had a SEC retention volume expected for a monomeric TMB (similar to the 8-stranded tOmpA monomer and Omp*Trans*3) and a far-UV CD spectrum characteristic of a β-sheet protein. Surprisingly, the well established band-shift assay on SDS-PAGE (*45*) was un-informative for the identification of folded *de novo* designed TMBs. Instead, we found a good agreement between the resistance of a design to protease digestion and a β-sheet characteristic far-UV CD spectrum even at high temperature (up to 95°C). Ten such designs were analyzed with ^1^H-^15^N HSQC NMR in DPC detergent micelles, and seven had well dispersed chemical shifts profiles characteristic of a folded protein in this detergent (Fig. S15, Fig. S16 - validated designs, Fig. S17 - examples of rejected designs, Fig. S18 - designs appearing as not folded by NMR). In total, 19 designs satisfied the biochemical screening criteria, suggesting that they fold into a TMB structure.

We selected two *de novo* designs (TMB2.17 (BLAST E-value to the non-redundant protein database: 0.10) and TMB2.3 (BLAST E-value: 0.035) and the Omp*Trans*3 construct for detailed biophysical characterization in a lipid bilayer to determine whether the proteins exhibit properties for a membrane spanning β-barrel (using tOmpA as a control for all our experiments). After refolding these four proteins into 100 nm DUPC LUVs, all proteins gave rise to far-UV CD spectra characteristic of a β-sheet both in 0.24 M and 2 M urea, and distinct from the spectra of the fully unfolded proteins in 8 M urea and from the proteins refolded in the absence of lipid (Fig. S10A, Fig. S20). We next determined the stability of the folded proteins by monitoring their ability to fold into/unfold out of LUVs at increasing urea concentrations, monitored by the change of fluorescence intensity between water-exposed and lipid embedded surface tryptophans (*46*). The results showed that the designed TMB proteins are more thermodynamically stable (midpoint urea concentration for folding (Cm^F^) 5.7 M and 7.2 M for TMB2.3 and TMB2.17, respectively, Fig. 3A) than tOmpA (Cm^F^ = 4.7 M), while Omp*Trans*3 is the most stable protein as it appears folded in 9 M urea (Fig. S19), in agreement with the far-UV CD data. It has been previously shown that many natural TMBs folding/unfolding transitions exhibit hysteresis due to the high kinetic barrier to unfolding and extraction from the membrane environment (*11, 47, 48*). Interestingly, in the conditions tested here, this behavior was observed for tOmpA but not for the designs TMB2.3 and TMB2.17 which showed superimposable and reversible unfolding/folding transitions, suggesting reduced kinetic stability relative to tOmpA. These observations likely explain the lack of a band-shift observed by SDS-PAGE, presumably since the lower kinetic stability causes the *de novo* designs to unfold during electrophoresis (*49*) (Fig. S21).

**Fig. 3.**
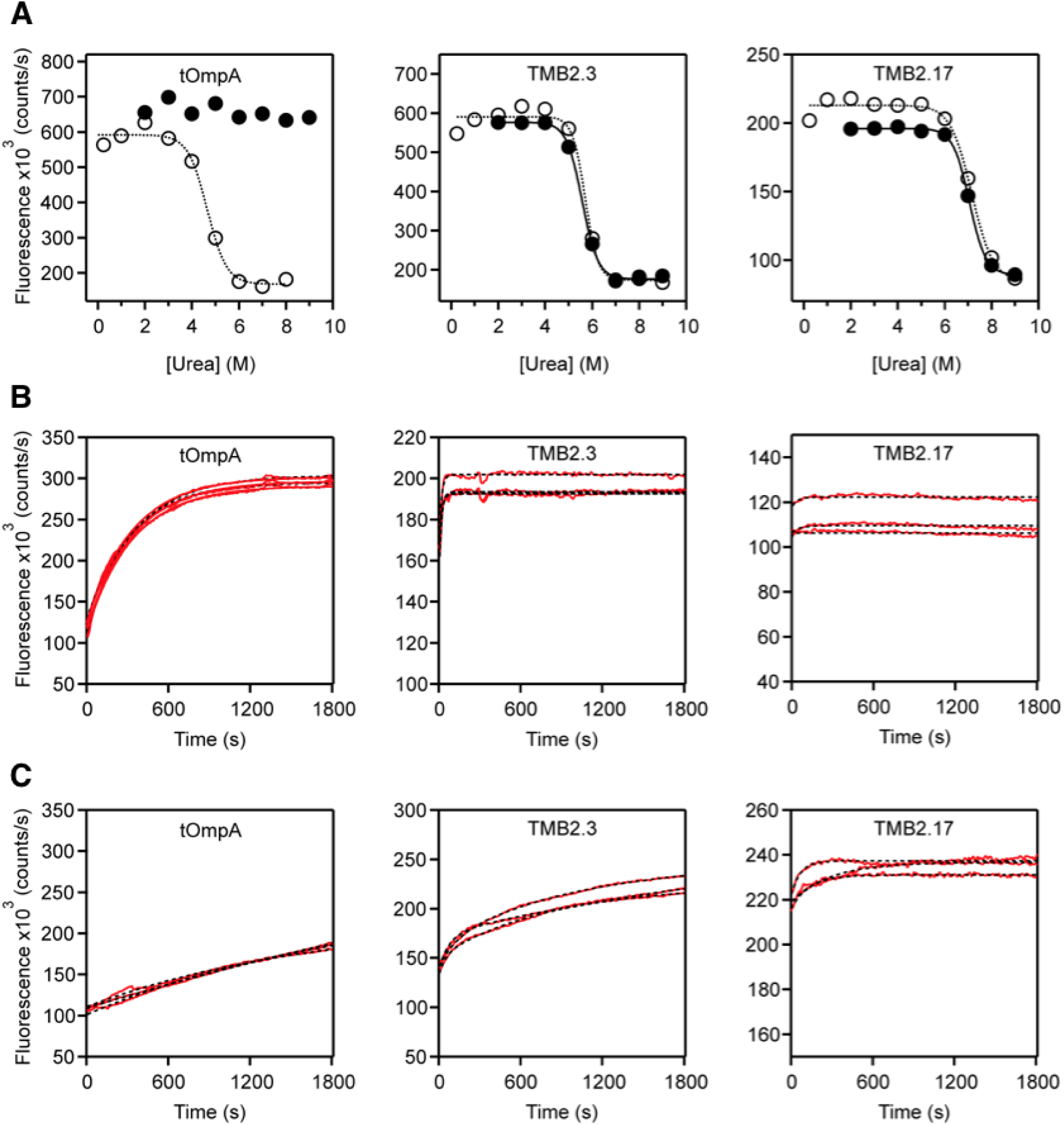
Biophysical characterisation of *de novo* designed TMB2.3 and TMB2.17 vs tOmpA in synthetic lipid membranes. (A) Urea dependence of folding and unfolding in DUPC LUVs. The fluorescence intensity at 335 nm was plotted against urea concentration to determine the midpoint urea concentration for folding (C_m_) (open circles, dashed line) and unfolding (C_m_) (filled circles, solid line). Kinetics of folding into (B) DUPC and (C) DMPC LUVs at an LPR of 3200:1 (mol/mol) in 50 mM glycine-NaOH pH 9.5, 2 M urea at 25 °C monitored by tryptophan fluorescence at 335 nm over 30 minutes (red line). Data were fitted with a single exponential function to determine folding rate constants (black dashed line). Three replicates are shown for each.

We next compared the kinetics of folding of the designed proteins to that of tOmpA (*50*) (Fig. 3B). These experiments showed that the designed TMBs fold over an order of magnitude more rapidly than tOmpA (folding rate constant of 3 x 10^-3^ s^-1^ for tOmpA) under identical conditions; with a rate too rapid to allow accurate measurement of the folding rate constant. Tryptophan fluorescence emission spectra of the end point of the folding reactions confirm the TMBs were indeed fully folded (Fig. S22). Finally, to confirm that the designs integrate into the lipid bilayer rather than folding on the lipid surface or in the absence of lipid, proteins dissolved in 8 M urea were diluted into 2 M urea without lipid or into LUVs composed of 1,2-dimyristoyl-sn-glycero-3-phosphocholine (DMPC, *di*C_14:0_PC). Consistent with previous results showing that the folding rates of natural TMBs are inversely correlated with lipid chain length (*9, 51*), the designed TMBs fold more slowly into lipids of longer acyl chain length (Fig. 3C), and do not fold in the absence of lipid (Fig. S23B), confirming that they indeed integrate into the lipid bilayer upon completion of their folding.

To characterize the structure of the designed TMBs in solution, we solved the structure of TMB2.3 folded into DPC detergent micelles using NMR spectroscopy (Table S3). Resonance peaks for 107 of the 117 non-proline residues of TMB2.3 were fully assigned; 6 more were partially assigned (Fig. 4A, Fig. S24A). Four out of six non-assigned residues were located in the *trans* β-turn regions - the remaining two were the N- and C-terminal residues of the protein. Analysis of the secondary structure content of TMB2.3, calculated using TALOS-N (*52*) is consistent with 8 β-strands that closely match the β-strand boundaries in the designed model (Fig. S24C). 9 out of 11 glycine residues pointing toward the core of the β-barrel (glycine kink residues) have the designed torsional irregularities based on the positive Cα chemical shifts [41] (Fig. S25A,B) and the more extended predicted backbone conformations (ɸ and Ѱ closer to 180°; Data S5). To validate the residue connectivity between the β-strands, we collected a total of 81 unique nuclear Overhauser Effects (NOEs) between amide protons; these suggest 72 inter-strand backbone hydrogen bonds that are in agreement with the β-strand connectivity of the design and the sheer number of 10 across the β-barrel (Fig. 4C, Fig. S24D). The NMR structure ensemble generated based on the chemical shifts and NOE information was in close agreement with the design model (average of 2.2 A RMSD, Fig. 4B). We observed low-intensity additional resonance peaks for a subset of residues, indicating the presence of a (minor) secondary conformation. The secondary signals strong enough for analysis were consistent with the secondary structure assignment and NOEs of the main conformation, indicating that the secondary conformation does not involve modification of the β-barrel architecture. Most of the residues producing double peaks cluster in the *cis* region of strands 1, 2 and 8 (Fig. S24B). Multiple resonance peaks might be explained by close proximity to the flexible N-terminus or by the transient dimeric interactions identified by native mass spectrometry in detergent micelles (Fig. S26, Fig. S27).

**Fig. 4.**
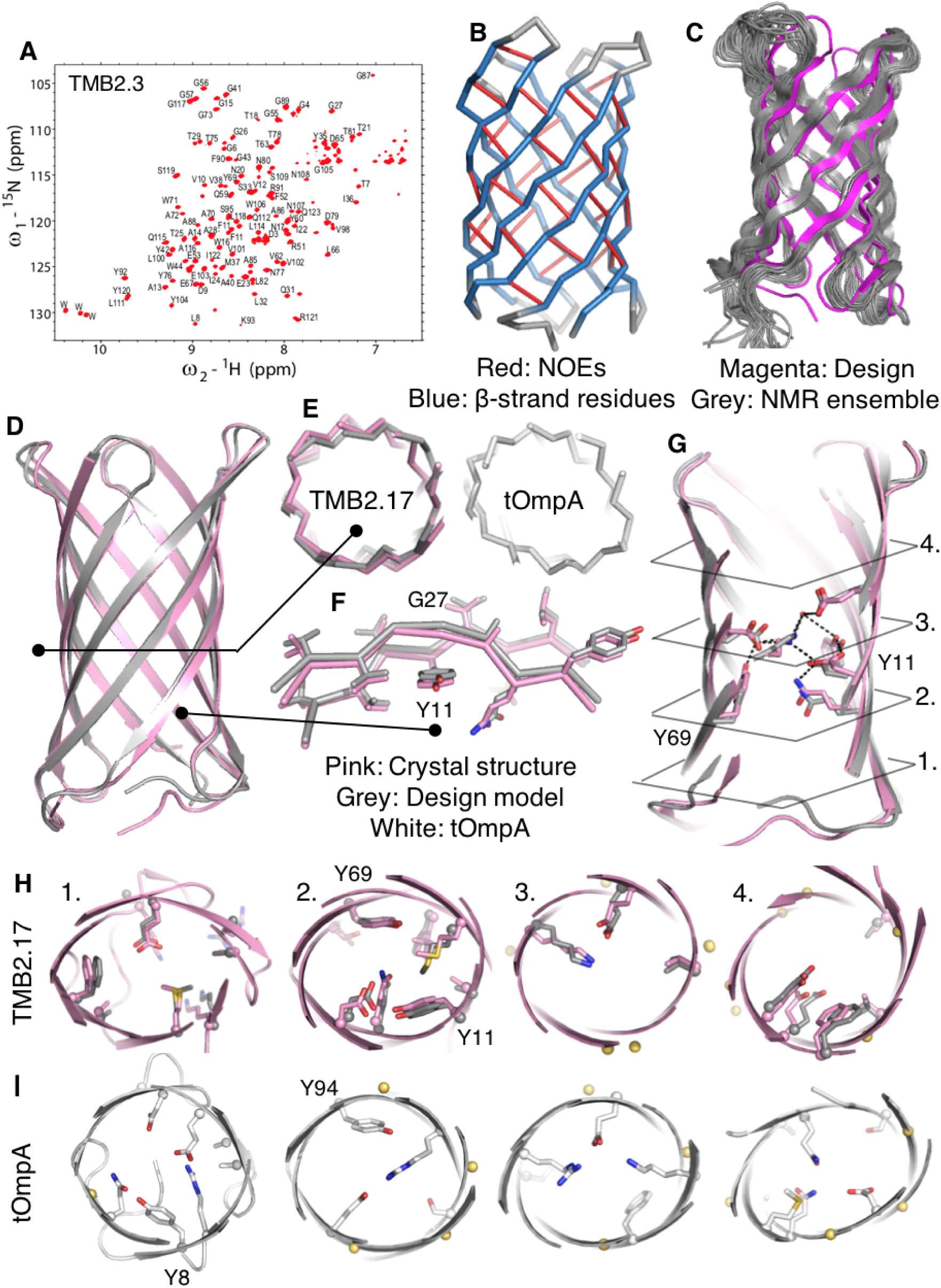
NMR structure of TMB2.3 in DPC detergent. (A) Assigned ^15^N-^1^H TROSY spectrum of TMB2.3. (B) NMR constraints mapped on the TMB2.3 design model. Residues predicted to have β-sheet secondary structure are colored in blue. Collected inter-residues NOEs are shown as red sticks. (C) TMB2.3 Rosetta design model (magenta) aligned to the 20 lowest energy models generated with NMR constraints (grey). (D-I) Superposition of the TMB2.17 crystal structure (pink) to the design model (grey) and comparison to the crystal structure of the naturally occurring tOmpA (white, PDB ID: 1QJP). (D) Full backbone superposition. (E) Comparison of the transverse β-barrel cross-section geometries. (F) Superposition of the β-strands around a mortise-tenon motif, showing the extended backbone conformation of the glycine kink (G27) and the rotamer of the tyrosine involved in the aromatic rescue interaction (Y11) which are nearly identical in crystal structure and design model. (G) Superposition of the side-chains involved in the core network of polar interactions around the two mortise-tenon motifs. The black lines indicate the locations of the four transverse slices for which core packing is shown in for the design model and crystal structure (H; the two are very similar) and compared to core packing in tOmpA (I) which is quite different. C*α* atoms are shown as spheres and glycine kink residues are colored in yellow; the positions of the tyrosines in the mortise/tenon folding motifs are labeled.

To determine the structure at the atomic level, we crystallized TMB2.17 and solved the structure at 2.05 Å resolution (Table S4). All but two residues located in one *trans* β-turn were resolved in the electron density map. The crystal structure of TMB2.17 closely matches the design model (1.1 Å backbone RMSD over all residues, Fig. 4d), and the β-barrel has a wide lumen delimited by glycines in an extended conformation that form kinks in the β-strands as designed (Fig. 4ef). The two YGD/E interactions (Y69, Y11, G27, G89, D39, E103) belonging to the extended mortise/tenon motifs are present in the crystal structure and the second shell of interactions, involving K71, E53 and Q29, is also properly recapitulated with additional interactions to water molecules (Fig. 4g); these extended side-chain hydrogen bond networks fill the lumen of the β-barrel. Overall, the buried amino acid side chain conformations and interactions in the design model are in very good agreement with the crystal structure (Fig. 4h; compare pink and gray). We compared the core of TMB2.17 to tOmpA, the most similar naturally occurring TMB whose structure has been determined (17% sequence identity, BLAST E-value of 1.6 against the non-redundant database, Fig. S28). The shape of the β-barrel lumen is quite different in the two proteins (Fig. 4e), as are the amino acid identities and packing arrangements of the core sidechains (compare the structure cross sections in Fig. 4H and 4I).

## Conclusions

The challenge of TMB *de novo* design is highlighted by the failure of the first three approaches we tried. The sequential approach previously used to build helical transmembrane proteins (*6*) - design and characterization of soluble proteins and subsequent hydrophobic residue re-surfacing to convert them to membrane proteins - yielded sequences strongly predicted to form amyloid. Designs with more polar cores which had high β-sheet propensity because of the perfect alternation of hydrophobic and polar residues systematically failed to express. Iterative improvement of the design protocol ultimately enabled the generation of a set of sequences with at least 8 % of sequences encoding proteins able to adopt a β-barrel fold (based on HSQC NMR). The NMR structure of one of these designs is very close to the design model. The power of our iterative “hypothesize, design, test” approach to explore the sequence landscape of membrane proteins is highlighted by the contrast between the failure in our first rounds of design, and the success in the final round in designing proteins that not only express and fold, but also have atomic structures nearly identical to the design model. The key to this success was introducing glycine kinks, β-bulges and register-defining sidechain interactions - also critical for the folding of water soluble β-barrels (*5, 53*) and hence important for defining β-barrel architecture irrespective of the solvent environment - and balancing hydrophobicity and β-sheet propensities of the sequences. The extent to which essentially all of the key design features are recapitulated with atomic level accuracy in the crystal structure of TMB2.17 suggests considerable control over TMB structure.

The overall β-sheet propensity and hydrophobicity of our successful designs are in the range of those of naturally-occurring TMBs sequences, suggesting that the natural TMBs might be under a similar negative selection pressure against formation of non-native β-sheet structures in aqueous environment (*54*). This is supported by our finding that replacing the tOmpA loops with short strong β-turn-nucleating sequences, but not by suboptimal turn sequences, blocks folding into a native β-barrel structure. Slowing down the folding and assembly of *trans* hairpins could allow more time for passage of the mostly hydrophilic amino acids in these β-strand connections across the lipid membrane, which likely has a large activation barrier. As well as encoding functional properties (*55–58*), the long loops commonly found on the *trans* side of the natural TMBs could play a role in slowing folding, although the energetic cost of translocation through the membrane would be much higher, consistent with the different kinetics of folding of tOmpA with long loops and short non-canonical turns. In Gram-negative bacteria, the BAM complex is responsible for accelerating the assembly of natural TMB substrates into the outer membrane by lowering the kinetic barrier to folding (*10*). Once folded, natural TMBs remain stably folded into the outer membrane, facilitated by both their kinetic stability (slow unfolding) and thermodynamic stability (ΔG°s range from ∼3-30 kcal/mol (*46, 59–61*)). Our design incorporates neither signals for BAM complex association nor evolution-conserved functional motifs and hence represent a “blank slate” for probing the tradeoffs between TMB folding, stability and function, as well as the underlying consequences and evolutionary constraints on OMP trafficking and biogenesis. Finally, the general design principles and methods we have described here - from the definition of the β-barrel architecture to the sequence properties - should be directly applicable to the design of larger pore containing β-barrels. The atomic level of accuracy in sidechain placement demonstrated by the crystal structure of TMB2.17 should enable custom design of transmembrane pores geometric and chemical properties tailored for specific applications.

## Acknowledgments

We thank I. Anishanka, Q. Cong, D. Kim, S. Berhanu Lemma, L. Stewart, L. Carter, X. Li, M. Dewitt and A. Saleem for their help for computational and experimental work; and L. Goldschmit and P. Vecchiato for IT support. We also thank many members of the Baker, IPD, Radford and Brockwell groups for discussions, the Advanced Photon Source beamline 24-ID-E for data collection.

## Author contributions

AAV and DB designed the research. AAV developed the design methods and expressed the designs with help of SRG, SM and CMC. AAV, SRG, SM and DCM cloned, expressed and characterized the tOmpA variants, supervised by KGF and DB. JEH and AAV conceived the biochemical screen and screened the designs with help of SM and CMC. CMC expressed isotopically labeled proteins and BL performed the NMR experiments and solved the NMR structure, supervised by LKT. PW designed, performed and analyzed the biophysical characterization in LUVs, with help from GNK and supervised by SER and DJB. AQS and SRH performed and analyzed native mass spectrometry experiment, supervised by VHW. ASK set up crystallization trays and AKB collected data and solved crystal structure. AAV and DB wrote the manuscript with input from all authors.

## Data and materials availability

The Rosetta software suite is available free of charge to academic users and can be downloaded from http://www.rosettacommons.org. The scripts, datasets, design models and MSAs used for this study have been deposited in Zenodo (DOI: 10.5281/zenodo.4068108). The NMR structure of the design TMB2.3 and the crystal structure of TMB2.17 have been deposited into PDB (6X1K, 6X9Z). Plasmids of the constructs are available upon request to the corresponding author.

## Supplementary materials

Materials and Methods Supplementary text Figures S1-S29

Tables S1-S4

External Databases S1-S5 References (66-98)

