## Supplementary materials for "*De novo* design of transmembrane β-barrels"

**This PDF file includes:**

Materials and Methods
Supplementary Text
Figs. S1 to S29
Tables S1 to S4
Captions for Data S1 to S5
References

**Other Supplementary Materials for this manuscript include the following:**

Data S1 to S5: sequences_designs.xlsx, designs_TMB_expression.xlsx, TMB_native_beta_turns.xlsx, designs_TMB_folding_screen.xlsx, talos_pred.xlsx

### Material and methods

Computational *de novo* design of a new protein with the Rosetta molecular modelling suite has two steps: first, a protein backbone is built, which is then used to guide the search for low energy sequence/structure pairs. For each of these steps, example scripts and shell commands are available from the Zenodo release DOI: 10.5281/zenodo.4068108.
Instructions to run each computational method are provided below.

<u>Backbone generation</u>

The same backbone generation approach ("backbone_generation.xml") was applied throughout this study and was described elsewhere (*5*, *66*). The desired protein backbone was described in a blueprint format ("TMB_blueprint"), where every residue in the protein was assigned a secondary structure type and a Ramachandran plot bin using Rosetta ABEGO type (*67*). The backbone-to-backbone hydrogen bond interactions for the protein were specified with constraints ("hbond_constraints"). To achieve control over the type of β-turns and torsional irregularities incorporated into the designed backbones, specific Ramachandran bins and hydrogen bonding patterns were assigned to β-turn, β-bulge and glycine kink residues. To design type I β-turns (3:5) on the *trans* side of the β-barrel, the `ABEGO` sequence "AAG" was used while type I β-turns on the *cis* side were designed with the `ABEGO` sequence "AA". A β-bulge was defined as a single residue in the alpha region of the ramachandran plot ("A" `ABEGO` type) with β-strand secondary structure. A glycine kink was defined as a single residue with a positive φ backbone angle ("E" `ABEGO` type) and a β-strand secondary structure.The rationale to design blueprints and specify constraints specific to β-barrels is provided in the supplementary text. The blueprint and constraints are used as input to the `BluePrintBDR` application (*21*) in Rosetta ("backbone_generation.xml"), which uses the information in the blueprint to pick fragments (9-mers and 3-mers) from crystal structures in the PDB and uses these fragments to search the structure space for low-energy structures using a Monte Carlo algorithm. Achieving enough conformational sampling to build all the hydrogen bonds in the β-barrel is computationally challenging, so the models produced by the `BluePrintBDR` are further minimized in the presence of the constraints and Rosetta's hydrogen bond potential (`hbond_lr_bb`) to drive the pairing between the β-strands. Every hydrogen bond is described with a distance constraint (between the N and O backbone atoms) and an angle constraint (the N--H--O angle). Such detailed description of the geometry of the interactions is necessary to compensate for Rosetta's inability to detect and score the hydrogen bonds that are located more than 5 Å apart in the input model and that are therefore excluded from the calculations of the interaction graph. The minimization step is done using a generalized rama potential ("Rama_XPG_3level.txt") and a coarse-grained energy function (Rosetta's centroid energy function), that was specifically optimized to balance long-range hydrogen bonding requirements with the local torsion angle requirements ("fldsgn_cen_omega02.wts"). The output of this design protocol is a set of three-dimensional protein backbone models with valine residues as placeholders at every position, except at the predefined glycine kink positions. High quality backbones to use in the sequence design step were selected based on the `vdw`, `rama` and `omega` scoring terms ("backbones_analysis.ipynb"). In this study, 10,000 backbone generation trajectories were necessary to obtain 200 backbones satisfying the quality criteria.

The Rosetta XML script was run using the following command (Rosetta version dadbb40b38ec2d4232fe9aa72241646019c4a758, 2017-05-11), where the input.pdb is a placeholder PDB file containing a single Ala residue.

```
< path to binary > /rosetta_scripts.default.linuxgccrelease
-database < path to database > -s input.pdb -parser:protocol
backbone_generation.xml -nstruct 10,000 -rama_map < path to
database > /scoring/score_functions/rama/Rama_XPG_3level.txt
```

Combinatorial sequence redesign of surface residues of water soluble β-barrels

The PDB coordinates of the previously designed water soluble beta-barrels (*5*) were used as template to redesign ("design_surface.xml") polar surface-exposed positions to hydrophobic amino acids (VILAF, resfile in "all.resfile"), with additional constraints ("girdle_cst") enforcing specific rotamers for aromatic girdle residues at the water/lipid interface. The ref2015 default Rosetta energy function (*68*) with modified reference energy for phenylalanine was used to limit the density of phenylalanines designed on the hydrophobic surface and match the distributions observed in naturally occurring TMBs (ref2015_F.wts). The lowest energy design was selected for each starting crystal structure out of five independent design trajectories.
The Rosetta XML script was run using the following command:

```
< path to binary >/rosetta_scripts -s input.pdb -parser:protocol
design_surface.xml -holes:dalphaball < path to Rosetta main
>/source/external/DAlpahBall/DAlphaBall.gcc -nstruct 10
```

Combinatorial sequence design

For all three generations of designs reported in this study (TMB0, TMB1, and TMB2), the search for a low-energy sequence was done over several rounds of iterative design following a genetic algorithm approach (~10% best scoring designs from one round of design were used as input for the next round of design). If necessary, changes were implemented to obtain designs to more closely match the hypothetical model that was tested.

Combinatorial sequence design of the design set TMB0

The set TMB0 was designed over four rounds of combinatorial sequence design ("design_gly.xml"). For all rounds of design, only polar amino acids were allowed in the core of the β-barrel, with the exception of the two tyrosines that occur in the mortise/tenon motifs; hydrophobic amino acids were allowed on the surface and aromatic amino acids at the lipid/water boundaries. All allowed amino acid combinations were specified in a resfile ("resfile"). After each round, the designs were selected based on the following criteria: 1) the correct rotameric state of the tyrosines 10 and 68, belonging to the mortise/tenon motifs, which is enforced with constraints during design ("mortise_tenon_cst"), and 2) the Rosetta `total_score` and four backbone quality metrics `omega, rama_prepro, p_aa_pp, and hbond_lr_bb`. The designs that scored better than the average for all four of the Rosetta metrics were selected for the next round of design ("analysis_21_02_16.ipynb"). These criteria typically eliminated approximately 90% of the initial designs with a correct mortise/tenon motif. For the last (fourth) round of design, a modified energy function with increased weights on the electrostatic interactions was used ("ref2015_fa_elec.wts") to favor more charged residues in the core. We hypothesized that a sharper contrast in hydrophobicity between the core and the surface of the β-barrel could improve the typical hydrophobic/polar alternation of residues characteristic of β-strands and hence improve β-strand secondary structure definition. Good definition of secondary structure elements on the sequence level is one of the key criteria for success of the design of new water-soluble protein folds (*21*).

The Rosetta XML script used for design rounds one to four was run with the following command (Rosetta version: f03bab6fdb3ea98290cd788afff2cb49878184cd, 2016-08-16):

```
< path to binary >/rosetta_scripts -database < path to database
> -s input.pdb -parser:protocol design_gly.xml -maxruntime 74000
-nstruct 10 -holes:dalphaball < path database >/DAlphaBall.icc
-beta
```

Combinatorial sequence design of the design set TMB1

To generate the set of designs TMB1, a small subset of designs generated after the third iteration in TMB0 (before the increase of the `fa_elec` weight to design more charged residues in the core) were selected. The surface was designed one more time with hydrophobic residues ("design.xml", "surface.resfile") to more closely match the amino acid probabilities on the surface of naturally occuring TMBs ("surface.comp").

Combinatorial sequence design of the design set TMB2

The first round of sequence design of the set TMB2 consisted of two stages. First, the centroid models from the backbone generation step were pre-designed in full-atom mode with Rosetta's default energy function ref2015 (*68*) ("design_1.xml") and by specifying allowed amino acids in the core and surface based on the inside-out model ("resfile_1"). The tyrosines in the mortise/tenon motifs were included at this stage and the specific rotamers characteristic of these interactions were enforced with constraints ("constraints_1"). The designs that scored better than average for Rosetta's `total_score`, `omega`, `rama_prepro` and `hbond_lr_bb` scores ("backbones_analysis.ipynb") were selected to serve as input models for the next design stage.

The Rosetta XML script used for full-atom pre-design of the full structure in round one was run with the following command (Rosetta version: dadbb40b38ec2d4232fe9aa72241646019c4a758, 2017-05-11):

```
< path to binary >/rosetta_scripts.default.linuxgccrelease
-database < path to database > -s input.pdb -parser:protocol
design_1.xml -nstruct 5 -rama_prepro_steep -beta_nov15
```

In the second stage, we searched for all the possible positions of aspartate or glutamate side-chains to act as a hydrogen bond acceptor to the tyrosines in the two mortise/tenon motifs. All the residues in the designs, except glycines, prolines and the two tyrosines (that belong to the mortise/tenon motifs), were mutated to alanine and the models were exhaustively searched for possible polar interactions stemming from the found D/E using the Rosetta HBNet protocol (*39*) ("hbnet.xml"). The parameters of the HBNet protocol, `hb_threshold` in particular, were adjusted to be able to consistently recover hydrogen bond interactions to the extent found in that of relaxed crystal structures of native TMBs. Each output model from the "hbnet.xml" run was relaxed with coordinate constraints ("fast_relax.xml"). The HBNet solutions found for each tyrosine of the mortise/tenon motif were recombined to generate all possible combinations of one or two designed mortise/tenon motifs or YGD/E motifs for every input backbone ("get_all_motifs.py").

The Rosetta XML script used to design full mortise/tenon motifs with HBNet (full-atom pre-design of the full structure) was run with the following command:

```
< path to binary >/rosetta_scripts -s input.pdb -beta_nov16
-parser:protocol hbnet.xml
```

The models generated with the "get_all_motifs.py" script (poly-alanine with the glycines, prolines and the designed YGD/E motifs) were used as input for the next round of sequence design. Three additional rounds of combinatorial sequence design were performed. The core and surface positions were designed independently in each round of design.
The Rosetta XML script used to design the core residues in round two ("design_core_all_round2.xml") was run with the following command (Rosetta version: 2018.52+HEAD.674a960 674a9609eaf25635511ef92f90f154dfb7a91256, 2018-12-24):

```
< path to binary >/rosetta_scripts -s input.pdb -parser:protocol
design_core_all_round2.xml -nstruct 5
```

For each input model, a constraints file and a resfile were generated. The resfile (Listing S1) defines the allowed amino acids in the β-turn regions and amino acid identities of the residues in the designed YGD/E motifs. A constraints file (Listing S2) was generated for each model to enforce the rotameric state of the tyrosine(s) in the motif(s) and to maintain the hydrogen bond interaction to the negatively charged amino acid. The resfile and constraints files were generated with the "get_all_motifs.py" script. The best designs were selected based on the energy of the hydrogen bond interactions between the tyrosine(s) and the negatively charged residue(s) and based on the total energy per residue of these negatively charged residue(s) evaluated with Rosetta ("select_best_motif_round2.ipynb").
In the surface design stage of round two ("design_surf_round2.xml"), the aromatic residues forming the aromatic girdle at the water/lipid boundaries were introduced ("surface_round2.resfile") and their rotameric state enforced with constraints ("constraints_surface_round2"). Since the core residues were allowed to repack during the surface residues design stage, the designs that retained low-energy YGD/E motifs were selected to move onto round three ("select_best_motif_round2.ipynb").
The Rosetta XML script used to design the surface residues in round two was run with the following command (Rosetta version: 2018.52+HEAD.674a960 674a9609eaf25635511ef92f90f154dfb7a91256, 2018-12-24):

```
< path to binary >/rosetta_scripts -s input.pdb -parser:protocol
design_surf_round2.xml -nstruct 10
```

Listing S1: Example of a resfile for an input model that had two mortise/tenon motifs with Y68 and Y10 which interact with D100 and D38, respectively. The columns left to right denote residue position, chain identifier, command to control sequence identity, and the list of allowed amino acids in their one-letter code. The variable section of the resfile, that specifies the unique mortise/tenon motifs for each input backbone, is highlighted in bold.

```
ALLAA # default command that applies to everything
#...... without a non-default setting; Allow all 20
#...... amino acids at all positions unless
#...... specified

start

2 A PIKAA NTEDPGRKQS # Allow only specified amino acids
#.................... at position 2
6 A PIKAA STDN

16 A PIKAA DENST
17 A PIKAA AES
18 A PIKAA D
19 A PIKAA G

32 A PIKAA STD
33 A PIKAA P
34 A PIKAA DEHTY

44 A PIKAA DENST
45 A PIKAA AES
46 A PIKAA D
47 A PIKAA G

62 A PIKAA STD
63 A PIKAA P
64 A PIKAA DEHTY

76 A PIKAA DENST
77 A PIKAA AES
78 A PIKAA D
79 A PIKAA G

81 A POLAR

94 A PIKAA STD
95 A PIKAA P
96 A PIKAA DEHTY

106 A PIKAA DENST
107 A PIKAA AES
108 A PIKAA D
109 A PIKAA G

68 A PIKAA Y # Allow only tyrosine at position 68
100 A PIKAA D
10 A PIKAA Y
38 A PIKAA D
```

Listing S2: Example of a constraints file for an input model that had two mortise/tenon motifs with Y68 and Y10 which interact with D100 and D38, respectively. Each line in the constraint file lists the constraint type (angle and distance, in this case) and constraint definition.

```
# Constrains dihedral angle of tyrosine atoms C ->
#.... CA -> CB -> CG at residue position 68. Dihedral
#.... angle is measured in radians on -pi -> pi. Function
#.... to be used on the angle constraint is
#.... "CIRCULARHARMONIC" with the specified definition.
Dihedral C 68 CA 68 CB 68 CG 68 CIRCULARHARMONIC -1.22 0.30
Dihedral C 10 CA 10 CB 10 CG 10 CIRCULARHARMONIC -1.22 0.30

# Constrains distance between atom pair OD1 and OH,
#...... at residue position 68 and 100 respectively.
#...... Function to be used on the atom pair constraint
#...... is "BOUNDED" with the specified definition.
AtomPair  OD1 38  OH  10 BOUNDED 2.6 3 0.2 hbond
AtomPair  OD1 100  OH  68 BOUNDED 2.39198 2.79198 0.2 hbond
```

The core of the β-barrel was designed again ("design_core_all_round3.xml"). The resfile (Listing S1) and constraints files (Listing S3) were adapted to the YGD/E motifs present in each design. Constraints to enforce the right rotameric states of the residues in the aromatic girdle were added to the previously used constraints files (Listing S2).

The Rosetta XML script used to design the core residues in round three ("design_core_all_round3.xml") was run with the following command (Rosetta version: 2019.01+HEAD.dbc838b6ae6 dbc838b6ae620b1293476b1bd4366ffc2facc5b5, 2019-01-03):

```
< path to binary >/rosetta_scripts -s input.pdb -parser:protocol ../design_core_all_round3.xml -nstruct 10
```

Listing S3: Example of a constraints file for an input model that had one mortise/tenon motif with Y68 which interacts with D102.

```
# Angle constraints for aromatic residues
Dihedral N 59 CA 59 CB 59 CG 59 CIRCULARHARMONIC -3 0.20
Dihedral N 15 CA 15 CB 15 CG 15 CIRCULARHARMONIC 3.0 0.20
Dihedral N 103 CA 103 CB 103 CG 103 CIRCULARHARMONIC 3.0 0.20
Dihedral N 29 CA 29 CB 29 CG 29 CIRCULARHARMONIC -3 0.20
Dihedral N 91 CA 91 CB 91 CG 91 CIRCULARHARMONIC -3.0 0.20
Dihedral N 119 CA 119 CB 119 CG 119 CIRCULARHARMONIC -3.0 0.20
Dihedral N 75 CA 75 CB 75 CG 75 CIRCULARHARMONIC 3.0 0.20
Dihedral N 41 CA 41 CB 41 CG 41 CIRCULARHARMONIC 3.0 0.20

# Distance and angle constraints between Y68 and D102
AtomPair  OE1 102  OH  68 BOUNDED 2.6 3 0.2 hbond
Dihedral C 68 CA 68 CB 68 CG 68 CIRCULARHARMONIC -1.22 0.30
```

All the designs from the core design stage of round three as well as the designs selected after round two were collected and the properties of the core polar interactions networks were analyzed in more detail. A custom Rosetta XML script ("filters.xml") was run to score the models based on packing of side chains around the glycine kinks, the packing of side-chains around the core polar network residues, and the number of unsatisfied hydrogen bonds in the core network of polar residues (Rosetta version: 2019.02+HEAD.81dca65976c 81dca65976c984579722d2b774ec357a16a8ad95, 2019-01-10):

```
< path to binary >/rosetta_scripts -s input.pdb -parser:protocol filters.xml -holes:dalphaball < path to main >/source/external/DAlpahBall/DAlphaBall.gcc
```

A Rosetta HBNet protocol was used to identify the existing hydrogen bond networks in the core of each design ("find_hbnet.xml") (Rosetta version: 2019.02+HEAD.81dca65976c 81dca65976c984579722d2b774ec357a16a8ad95, 2019-01-10). Only networks comprising the tyrosines belonging to the YGD/E motifs were analyzed.

```
< path to binary >/rosetta_scripts -s input.pdb -parser:protocol find_hbnet.xml
```

The outputs of the two scripts were used to compute the size, energy and saturation of the networks and the number of satisfied and unsatisfied hydrogen bonds. These metrics, Rosetta's side-chain hydrogen bond score (`hbond_sc`) and the metrics computed using the "filters.xml" script were used to select the designs with the most extensive and stable core networks for the next round of surface design ("filter_networks.ipynb").

For the surface design stage of round three ("design_surf_round3.xml"), glycines were allowed in lipid-exposed surface positions ("surface_gly_round3.resfile") and the weight on the long-range hydrogen bond potential (`hbond_lr_bb`) was increased to 2.0 to find strained positions on surface and design them into glycine. The rotameric state of the residues belonging to the aromatic girdles was enforced with constraints (Listing S4). The core networks were allowed to repack during the surface design stage, and the designs with the highest retention of these networks after repacking and lowest Rosetta `omega` score were selected ("analyse_round3_surf.ipynb"). Seven hundred and seventy-five designs were selected following this procedure. After manual inspection of the core network of hydrogen bonds, two hundred and four designs were excluded for presenting unsatisfied polar atoms potentially buried in hydrophobic pockets (which is difficult to detect automatically in a reliable way). The four hundred and eighty-eight designs with the lowest total side-chain to side-chain hydrogen bond energy (`hbond_sc`) were selected for the last stage of combinatorial sequence design ("cluster_round4.ipynb").

The Rosetta XML script used to design the surface residues in round three was run with the following command (Rosetta version: 2019.05+HEAD.8e9940c32d6 8e9940c32d62464b0d303e1706faa0cdf22218a9, 2019-01-30):

```
< path to binary >/rosetta_scripts -s input.pdb -parser:protocol design_surf_round3.xml -nstruct 10
```

Listing S4: Example of a constraints file used to constrain the rotameric state of aromatic residues in the aromatic girdles at the water/lipid interface during the surface design stage of round three.

```
Dihedral N 29 CA 29 CB 29 CG 29 CIRCULARHARMONIC -3 0.20
Dihedral N 59 CA 59 CB 59 CG 59 CIRCULARHARMONIC -3 0.20
Dihedral N 91 CA 91 CB 91 CG 91 CIRCULARHARMONIC -3.0 0.20
Dihedral N 119 CA 119 CB 119 CG 119 CIRCULARHARMONIC -3.0 0.20
Dihedral N 15 CA 15 CB 15 CG 15 CIRCULARHARMONIC 3.0 0.20
Dihedral N 41 CA 41 CB 41 CG 41 CIRCULARHARMONIC 3.0 0.20
```

```
Dihedral N 75 CA 10 CB 75 CG 75 CIRCULARHARMONIC 3.0 0.20
Dihedral N 103 CA 103 CB 103 CG 103 CIRCULARHARMONIC 3.0 0.20
```

The designs were manually clustered based on the similarity of their core hydrogen bond networks ("cluster_round4.ipynb"). The amino acids on the surface of these designs were designed one more time ("design_surf_round4.xml") to incorporate phenylalanines and therefore increase the hydrophobicity of the lipid-exposed surface of the β-barrel. Since it is an artefact of Rosetta's energy function to excessively favor phenylalanine amino acids, the reference weight for phenylalanines was modified in the default energy function ("ref2015_F4.wts") to incorporate phenylalanines at a rate similar to what is observed in naturally occuring TMBs. The rotameric state of the residues belonging to the aromatic girdle was enforced with constraints that were used for the previous rounds of surface design (Listing S4). A resfile was used to define allowed amino acids on the lipid exposed surface (VILAF) excluding the positions that have been previously designed as glycine or proline (Listing S5). For each input model, ten independent surface design trajectories were run and the lowest energy design (`total_score`) was selected ("analyse_clusters.ipynb").

The ninety ordered designs were selected to span each of these structural clusters as well as a broad range of hydrophobicity of the core and propensity for β-sheet and alpha-helix secondary structure (as predicted with RaptorX). The analysis and selection criteria can be found in the provided Jupyter Notebooks ("analyse_round4.ipynb" to select TMB2.1 to TMB2.20 that have unique core networks that do not belong to any existing cluster; "analyse_clusters.ipynb" to select designs TMB2.21 to TMB2.90 from the network clusters). The placeholder sequences of the *trans* β-turn used throughout the design process were replaced with the suboptimal sequences necessary for TMB folding identified in this study.

The Rosetta XML script used to design the surface residues in round three ("design_surf_round4.xml") was run with the following command (Rosetta version: 2019.26+HEAD.98bf209eda5 98bf209eda57a65099b83743e89bc0631d112bdc, 2019-06-25):

```
< path to binary >/rosetta_scripts -s input.pdb -parser:protocol design_surf_round4.xml -holes:dalphaball < path to main >/source/external/DAlpahBall/DAlphaBall.gcc -nstruct 10
```

Listing S5: Example of a resfile used during the surface design stage of round four for an input model that had two surface glycines at positions 55 and 25.

```
NATAA
start

7 A PIKAA VILF
9 A PIKAA VILAF
11 A PIKAA VILAF
13 A PIKAA VILAF

21 A PIKAA VILF
23 A PIKAA PVILAF
# 25 A PIKAA VILAF
27 A PIKAA VILAF
31 A PIKAA VILF
```

```
35 A PIKAA VILF
36 A PIKAA AST
37 A PIKAA VILAF
39 A PIKAA VILAF

51 A PIKAA VILAF
53 A PIKAA VILAF
# 55 A PIKAA VILAF
57 A PIKAA VILAF
58 A NOTAA CPGWYI
60 A NOTAA CPGFWYI
61 A PIKAA VILF

65 A PIKAA VILF
69 A PIKAA VILAF
71 A PIKAA VILAF
73 A PIKAA VILAF
75 A PIKAA WY

81 A PIKAA VILAF
85 A PIKAA VILAF
87 A PIKAA VILAF
89 A PIKAA VILAF
93 A PIKAA VILF

97 A PIKAA VILF
99 A PIKAA VILAF
101 A PIKAA VILAF

113 A PIKAA VILAF
115 A PIKAA VILAF
117 A PIKAA VILAF
121 A PIKAA VILF
```

Rosetta simulations with PPM predictions

The protein backbones for the tested topologies were generated based on blueprints and constraints files provided in the GitHub repository. A sequence was designed for each of the 20-25 best scoring backbones following the inside-out model and with aromatic residues at membrane anchoring positions to the β-turns to define the aromatic girdle. The 20-25 models were submitted to the PPM server to define its position in the lipid bilayer. The tilt angles, water-to-lipid partition energies and hydrophobic thicknesses were averaged per topology. For every tested topology, an average molecular model was generated by averaging the heavy atoms of the proteins as well as the planes defining the lipid membrane leaflets (“average_hydrophobic_thickness.ipynb”). Such an average model was used to verify the continuity of the hydrophobic thickness.

Computational simulation of the structure/energy landscape of β-turns

To compute structure/energy landscapes for the β-turn sequences, one low energy poly-valine TMB backbone was selected for the simulation and the *trans* β-turn positions and two additional β-strand flanking residues on both sides of the β-turn were mutated to the target sequence. The backbones conformations were readjusted to the new sequences by running the Rosetta FastRelax protocol. Two hundred fifty loop conformations were generated by independent KIC sampling and scored with Rosetta's default energy function (Listing S6, Listing S7). To do so the Rosetta loopmodel protocol was run with KIC backbone perturbation, using the following command line (Rosetta version: 2019.45+HEAD.fc8ab401178 fc8ab401178f6f099884fd5a39597b1a9cac8db2 (November 2019)):

```
< path to binary >/loopmodel @flags.kic -s input.pdb -nstruct 250 -out:file:silent loop.out
```

The RMSD of each generated loop conformation to the conformation in the starting model (canonical backbone for the 3:5 type I β-turn with a G1 β-bulge) was calculated.

Listing S6: Example of a flags.kic file used to run the loopmodel protocol.

```
-loops:remodel perturb_kic
-loops:refine refine_kic
-kic_rama2b
-loops:ramp_fa_rep
-loops:ramp_rama
-loops:refine_outer_cycles 1
-loops:max_inner_cycles 1
-kic_omega_sampling
-allow_omega_move true
-ex1
-ex2
-loops:kic_recover_last

-loops:loop_file loopdef.loop
-overwrite
```

Listing S7: Example of a loop definition file (perturbation of the first *trans* β-turn).

```
15 21 21 0 1
```

<u>Amino acid propensities in naturally occuring 8-strands TMBs</u>

The multiple sequence alignments (MSA) were generated by searching for homologs of 8-strands TMBs with crystal structures deposited in the PDB (1qjp, 2f1v, 1thq, 1qj8, 2k0l, 2mlh, 1p4t, 4fuv, 4rlc, 2n6l, 2lhf, 2erv, 3qra) using GREMLIN (*69*). The sequences in the MSA were merged and filtered for maximum 90% sequence similarity with CD-HIT (*70*). The MSA is provided in the GitHub repository.

To compute the amino acid compositions of the transmembrane β-strands and the β-turns, we assumed that the interaction with the lipid membrane constrains the evolution of the β-barrel architecture and results in constant position of the transmembrane regions in the sequence of the protein. This hypothesis was supported by the comparison between the amino acid compositions computed with our method and the statistic reported in a previous study based on crystal structures of TMBs with different strand lengths (*71*) (Fig. S3). The regions of the MSA corresponding to the transmembrane β-strands or the β-turns were identified based on the crystal structures of the query sequences, extracted from the MSA and used for the downstream analysis. The transmembrane β-strand regions were defined as the span from the membrane anchor position from one side of the membrane to the membrane anchor position to the other side of the membrane.

To investigate how the well the β-turn structure is defined by the sequence profiles derived from the MSA, we used Rosetta fragment_picker protocol (*72*) to pick fragments from crystal structures in the PDB . Only the sequence profiles from the MSA were considered for fragment picking (Listing S8). We compared the *cis* and *trans* β-turn sequence profiles for identical types of β-turn backbones on the same protein to avoid potential bias from MSA depth.

The fragment picking application was run with the following command:

```
< path to binary >/fragment_picker -in:file:vall id30_d150314_sprf_7mer_50frag_filt.vall frags:n_candidates 1000 -frags:n_frags 200 -frags:write_ca_coordinates -in:file:fasta msa.fasta -frags:scoring:config t000__scores9.cfg -frags:frag_sizes 9
```

Listing S8: Score weight used to pick fragments based on the MSA ("t000__scores9.cfg").

```
# score name         priority  wght   min_allowed  extras
ProfileScoreL1           700     1.0      -
```

Protein expression and purification

Codon-optimized genes encoding the TMB and tOmpA loop variants were synthesized and cloned into the pET-29 vector (Integrated DNA technologies). The natural tOmpA and full-length OmpA genes were cloned into the same vector from the *E. coli* K-12 strain. The OmpA, tOmpA and OmpAAG constructs were originally expressed with a C-terminal 6xHis-tag fusion, which did not influence the ability of the protein to fold into lipid membrane or detergent micelles. However, the Omp*Trans* and TMB designs were not fused to the 6xHis-tag because his-tagged proteins were found to produce less compact and more difficult to purify inclusion bodies. Plasmids were transformed into BL21*(DE3) *E. coli* strain (NEB). Protein expression was induced by overnight growth at 37°C in the Studier autoinduction medium and replicated at least twice for the designs from set TMB0, the designs TMB2.1 to TMB2.20 and the designs TMB2.21-TMB2.90 that failed to express. To isolate the proteins in inclusion bodies, the cells were lysed either by sonication (50 ml cultures for design screening) or with a MicroFluidizer (Microfluidics) in lysis buffer (50 mM Tris pH 8.0, 40 mM EDTA pH 8.0). The cell lysate was incubated for 60 min at 4°C with 0.1 % of Brij-35. The inclusion bodies were collected by centrifugation, re-suspended in the washing buffer (10 mM Tris pH 8.0, 1 mM EDTA pH 8.0) by sonication and pelleted again. The washing step was repeated three times. The pellets were stored at -20°C. The proteins prepared for the small scale screening assay were dissolved in 6 M urea and used immediately. The proteins prepared for biochemical and structural characterization were first dissolved in 8 M guanidinium chloride (GuCl) and further purified by Akta Pure fast protein liquid chromatography (GE Healthcare) using a Superdex 75 increase 10/300 GL column (GE Healthcare) in denaturing conditions.

Expression of $^{15}N$ and $^{1}H$-$^{15}N$-$^{13}C$ isotopically labelled proteins

*$^{15}N$ isotopically labelled proteins*

A LB media starter culture was prepared at equal volume to the desired expression volume and grown overnight at 37˚C, 200 rpm. Cells were harvested at 4,000 RPM, 4˚C for 10 minutes or until a solid pellet forms. Cell pellet was gently resuspended (do not vortex) with M9 minimal media (30mM $Na_2HPO_4$, 20mM $KH_2PO_4$, 10mM NaCl, 10mM $NH_4Cl$, 0.2% glucose, 1mM $MgSO_4$, 0.1mM $CaCl_2$, 0.01g/L biotin, 0.01g/L thiamin, 1x trace metals, appropriate antibiotic) with $^{15}N$-$NH_4Cl$ (Cambridge Isotopes). Cultures were grown at 37˚C, 200 rpm. $OD_{600}$ was measured after 2 hours after inoculation. Cultures were induced with 0.5mM IPTG at $OD_{600}$ 0.8-1.0 and grown overnight at 22˚C, 200 rpm. 500µL of pre-induced culture was retained for later analysis. Cells were harvested at 4,000 RPM, 4˚C for 10 minutes. Supernatant was discarded and the cell pellet was stored at -80˚C or used immediately for protein purification. Protein expression was assessed via SDS-PAGE with pre- and post-induction retain samples.

*$^{1}H$-$^{15}N$-$^{13}C$ isotopically labelled proteins*

Due to decreased cell growth and protein expression yields in the presence of $D_2O$, the gradual introduction of deuterated media is recommended. A 5mL starter culture in 100% $H_2O$ LB media was prepared and the percentage of $D_2O$ LB media was increased in a stepwise fashion (100% $H_2O$:0% $D_2O$, 75:25, 50:50, 25:75, 0:100). Cultures were grown at 37˚C, 200 rpm overnight prior to a 1:10 inoculation ratio for subsequent steps. 0.2% glucose was added to LB media to promote cell growth. A glycerol stock was prepared when the bacterial culture has adopted 100% deuterated media, the remaining overnight was used to start an expression culture. Protein was expressed and harvested using the previously described $^{15}N$ isotopically labelled proteins protocol using M9 media containing $^{15}N$-$NH_4Cl$ (Cambridge Isotopes) and 0.2% $^{13}C$-glucose (Cambridge Isotopes), in deuterium.

Screening assay in detergent micelles

The first twenty TMB2 designs (and their variants with tOmpA loop inserts) were tested in DDM detergent micelles. We later switched to DPC detergent for improved refolding efficiency (by comparing the refolding efficiency of a few designs in both detergents by HSQC NMR) and to simplify the interpretation of the results. For a few designs, the screening assay was repeated in OG detergent micelles. Before the folding experiment, the protein pellets were dissolved in urea and centrifuged 30 min at maximum speed. The concentration of protein in the supernatant was measured using a nanodrop and the stocks were diluted to 80 µM. 250 µM of the 80 µM stock solutions were diluted drop-by-drop into 5 ml of vortexed refolding buffer (20 mM Tris pH 8.0, 150 mM NaCl, 2X CMC detergent). DPC detergent was used at a concentration of 0.1%; DDM detergent was used at a concentration of 0.02%; OG detergent was used at a concentration of 1%. In parallel, 250 µM of the 80 µM stock solutions were diluted drop-by-drop into 5 ml of TBS buffer (20 mM Tris pH 8.0, 150 mM NaCl) to test the solubility of the design in the absence of detergent. The samples were incubated overnight at 4°C on a rocker. To assess protein solubility, 20 µl of each sample and the corresponding control without detergent were centrifuged 30 min at maximum speed and analyzed on SDS-PAGE. A non-centrifuged sample was analyzed alongside them to provide the total protein band. The samples prepared in detergent were concentrated to 1 ml in an Amicon Ultracentrifugation device with a cut-off of 10 kDa (Merck Millipore). After centrifugation for 30 min at maximum speed, the protein/detergent complexes were separated from larger aggregates using a Superdex 200 increase 10/300 GL SEC column (GE Healthcare) in the refolding buffer. If a major species with a retention volume compatible with a monomeric 8-strands TMB was detected by SEC, that species was further tested for the presence of a heat-modifiable species (SDS-PAGE band-shift assay), for resistance to proteases and for a β-sheet characteristic far-UV CD spectrum.

Far UV circular dichroism spectroscopy

For the TMB screening in detergent micelles, the protein/detergent complex collected out of SEC was directly analyzed by CD spectrometry in SEC buffer (20 mM Tris pH 8.0, 150 mM NaCl, 2X CMC detergent). CD spectra were obtained using a Jasco model J-1500 spectropolarimeter over a wavelength range of 260–190 nm. The temperature was controlled with a Peltier and spectra were recorded every 10°C, from 25°C to 95°C. One last spectrum was recorded after cooling the sample down back to 25°C. For detailed biophysical characterization of designs TMB2.3 and TMB2.17 in synthetic lipid membranes, the TMBs denatured in 50 mM glycine-NaOH pH 9.5, 8 M urea were diluted into DUPC LUVs in 50 mM glycine-NaOH pH 9.5 containing 0.24 M, 2M and 8 M urea, and folding was allowed to proceed overnight at 25°C. The final protein concentration was 6 μM the lipid/protein ratio (LPR) was 600:1 (mol/mol). Average CD spectra from four repeats were obtained using a Chirascan Plus (Applied Photophysics) spectropolarimeter equipped with Peltier temperature controller set at 25°C, over a wavelength range of 260–190 nm, a digital integration time of 2 seconds, and a 2 nm bandwidth.

Protease challenge

Trypsin-EDTA (0.25%) solution was purchased from Life Technologies and stored at stock concentration (2.5 mg/mL) at -20°C. α-Chymotrypsin from bovine pancreas was purchased from Sigma-Aldrich as lyophilized powder and stored at 1 mg/mL in TBS +100 mM $CaCl_2$ at -20°C. A sample of the protein/detergent complex collected out of SEC was directly subject to a test for protease resistance. 19 μl of the protein/detergent sample were mixed with 1 μl of Trypsin-EDTA and another 19 μl sample was treated with 1 μl of α-Chymotrypsin. The samples were incubated 15 min at Room Temperature. The reaction was quenched with 2X Laemmli Sample Buffer (BioRad). The samples were heated at 95°C for 10 min and analyzed on SDS-PAGE gel (Any kD™ Mini-PROTEAN® TGX™ Precast Protein Gels, BioRad) alongside an undigested sample.

SDS-PAGE band-shift assay

In the context of TMB screening in detergent micelles, 2X 20 μl of each sample collected from SEC were mixed with 2X Laemmli Sample Buffer (BioRad). For each tested protein, one sample was heated at 95°C for 10 min while the other sample was kept at room temperature. The samples were analyzed on a SDS-PAGE gel (Any kD™ Mini-PROTEAN® TGX™ Precast Protein Gels, BioRad). For detailed biophysical characterization of designs TMB2.3 and TMB2.17, samples of the folding reaction used for far-UV CD were mixed with 4x SDS-PAGE loading buffer (200 mM Tris-HCl pH 6.8, 6% (w/v) SDS, 40% (v/v) glycerol, 0.004% (w/v) bromophenol blue, and folded/unfolded species were resolved on a 15 % (w/v) acrylamide/bis-acrylamide (37.5:1) Tris-Tricine SDS-PAGE gel at pH 8.45 operating at 60 mA for 90 minutes at room temperature. Boiled samples were heated to >95°C for 10 minutes. Gels were stained with InstantBlue™ (Expedeon) and imaged using an Alliance Q9 Advanced gel doc (UVITEC, Cambridge, UK).

Equilibrium folding/unfolding by tryptophan fluorescence

To determine the urea dependence of TMB folding, urea denatured TMBs in 50 mM glycine-NaOH pH 9.5, 8 M urea were diluted into DUPC LUVs at an LPR of 600:1 (mol/mol) in 50 mM glycine-NaOH pH 9.5 containing 0.24-9 M urea, and folding was allowed to proceed overnight at 25 °C. To measure the urea dependence of unfolding, TMBs were initially folded in DUPC LUVs at an LPR of 600:1 (mol/mol) in 50 mM glycine-NaOH pH 9.5, 2 M urea overnight at 25 °C. The folded TMB stock was then diluted 10-fold into 50 mM glycine-NaOH pH 9.5 containing 2-9 M urea and incubated overnight at 25 °C to initiate unfolding. The final protein concentration was 0.4 µM and the LPR was 600:1 (mol/mol). Tryptophan fluorescence emission spectra were obtained using a PTI QuantaMaster spectrofluorometer (Photon Technology International) in QS quartz cuvettes with excitation slits set to 1 nm and emission slits set to 5 nm. Fluorescence was excited at 280 nm and emission spectra were acquired between 300-400 nm using a step size of 1 nm and an integration time 1 second. The fluorescence intensity at 335 nm was plotted against the urea concentration and data were fitted with a sigmoid function to extract the urea concentration midpoint for folding ($Cm^{F}$) and unfolding ($Cm^{UF}$).

Folding kinetics measured using tryptophan fluorescence

Kinetics of TMB folding into DUPC and DMPC LUVs were measured at a final OMP concentration of 0.4 µM and an LPR of 3200:1 (mol/mol). The TMB unfolded proteins were diluted 20-fold from 8 M urea into LUVs created from DUPC or DMPC in 50 mM glycine-NaOH pH 9.5 containing 2 M or 9 M urea. The choice of using 2 M urea to monitor TMB folding was made based on the results of the band-shift assay on SDS-PAGE (Fig. S21), that showed partial aggregation of tOmpA at lower concentrations of urea. TMBs were also diluted from 8 M urea in 2 M urea without lipids to determine the lipid dependence of folding. Upon addition of denatured TMBs to LUVs in the folding buffer pre-equilibrated at 25 °C, the reaction was mixed rapidly and fluorescence emission was monitored at 335 nm following excitation at 280 nm over 30 minutes. Excitation slits were set to 0.5 nm, emission slits were 5 nm, the bandwidth was 1 nm and integration time was 2 seconds. Kinetics were measured in triplicate and, where possible, were globally fitted to a single exponential function to extract folding rate constants.

NMR Spectroscopy and Structural Calculations

All NMR spectra were collected on a Bruker Avance 800 MHz spectrometer equipped with a cold-probe. For initial sample optimization and screening, 2D TROSY-HSQC spectra were collected for $^{15}$N-labeled samples. For backbone assignments of the TMB2.3, TROSY-versions of 3D experiments [HNCA, HN(CA)CB, HNCO, HN(CA)CO] were collect on a $^{2}$H, $^{13}$C, $^{15}$N-labeled sample with a non-uniformed sampling (NUS) technique. Two 3D NOE experiments, $^{15}$N-$^{15}$N-$^{1}$H HSQC-NOESY-HSQC and $^{15}$N-$^{1}$H-$^{1}$H NOESY-TROSY, were performed with mixing times of 120 ms, also in the NUS mode. In addition, a TROSY-based 2D $^{1}$H-$^{15}$N heteronuclear NOE experiment was collected with a saturation recovery delay of 5 s with an interleaved approach. All spectra were processed and analyzed with NMRPipe (*73*) and Sparky (*74*), and in particular, NUS scheduling and reconstruction were carried out with hmsIST (*75*).

The presence of a well-ordered TMB2.3 structure was supported by NMR dynamics measurements. We measured $^{1}H$-$^{15}N$ heteronuclear NOE values for non-overlapped 101 residues, the high average of 0.83 ± 0.11 indicated restricted motions for the whole protein. Dihedral angle restraints were predicted from the TALOS-N program (*76*) on the basis of the experimental Ca, Cb, CO, N, and HN chemical shifts. Good predictions from TALOS-N were converted to input values for structural calculations with tolerances at either twice the standard deviations or 20°, whichever the larger value. All assigned NOE peaks were converted to NOE distances using their peak height values and calibrated based on the fact that the average HN-HN distance between anti-parallel beta-strands is 3.3 Å. For structure calculations, the distances were categorized as having strong, medium, and weak NOEs with upper limits of 3.5, 5.0, 6.0 Å, respectively. The presences of hydrogen bonds were determined by strong NOEs in both NOE spectra, as well as their beta-sheet secondary chemical shifts. Each hydrogen bond was constrained with two upper limits of 2.5 and 3.5 Å for HN…O and N…O, respectively. Structural calculations were performed using Xplor-NIH v2.39 (*77*) from an extended structure with the default anneal.py script. A total of 200 structures were calculated, and the final 20 structures were selected based on the lowest total violation energies.

Native mass spectrometry in detergent micelles

tOmpA, OmpAAG, Omp*Trans*2, and Omp*Trans*3 proteins were analyzed by native mass spectrometry (MS) using a Thermo Q ExactiveTM Ultrahigh Mass Range (UHMR) Orbitrap instrument (Thermo Fisher Scientific, Bremen, Germany). Prior to MS analysis, protein samples received in 20mM Tris, 150mM NaCl, 0.02% n-Dodecyl-β-D-Maltopyranoside (DDM), pH 8.0 were buffer exchanged into 200 mM ammonium acetate, 2X CMC DDM, pH 8.0 using Micro Bio-Spin P6 columns with a 6 kDa cutoff (Bio-Rad, Hercules, CA, USA). Proteins were analyzed at concentrations of 3-4 µM monomer. Ions were generated via nano-electrospray ionization using borosilicate capillaries pulled in-house using a micropipette tip puller (Sutter Instruments model P-97, Novato, CA). The protein solution was inserted into the capillary and a platinum wire was inserted into the solution. A spray voltage of 0.5-1.0 kV was used for all experiments. Following ionization, in-source trapping (typically 250-275 V) was used to remove the detergent micelles in the gas phase. Voltages were applied throughout the instrument to optimize ion transmission while minimizing unnecessary ion activation. Mass spectra were collected at a resolution (@ m/z 400) of 12,000 to determine relative ratios of proteins present and at a resolution of 100,000 for confirmation of proteins by accurate mass. Mass spectra were deconvoluted using UniDec version 4.0.0 Beta (*78*).

Crystallization

TMB2.17 purified in denaturing conditions was refolded by rapid dilution from 80 µM to 4 µM into a buffer containing 2X CMC of DPC detergent. The solution was incubated at room temperature overnight to allow the proteins to fold and the sample was concentrated to 1 ml using an Amicon Ultra 10 kDa centrifugation device (20 - 25 mg/ml protein). The protein/detergent complex was further purified by SEC on a Superdex 200 increase 10/300 GL column (GE Healthcare) and dialysed against 20 mM Tris 150 mM NaCl pH 8.0, 2X CMC of DPC detergent. Both LCP and classical sitting drops were set up in DPC using Mosquito LCP by STP Labtech. Diffraction quality crystals appeared in D10 (0.1 M Tris at pH 8.5 and 10 % PEG8000) of MemStart+MemSys HT by Molecular Dimensions. Crystals were subsequently harvested in a cryo-loop and flash frozen directly in liquid nitrogen for synchrotron data collection.

Data collection

Data collection from crystal of TMB2-17 was performed with synchrotron radiation at the Advanced Photon Source (APS), 24ID-E. Crystals belonged to space group *R 3 :H* with cell dimensions $a = b = 51.08$ Å, and $c = 116.71$ Å, $\alpha = \beta = 90°$ and $\gamma = 120°$. X-ray intensities and data reduction were evaluated and integrated using XDS (*79*) and merged/scaled using Pointless/Aimless in the CCP4 program suite (*80*).

Structure determination and refinement

Starting phases were obtained by molecular replacement using Phaser (*81*) using the designed model. Following molecular replacement, the models were improved using phenix.autobuild (*82*); efforts were made to reduce model bias by setting rebuild-in-place to false, and using simulated annealing and prime-and-switch phasing. Structures were refined in Phenix. Model building was performed using COOT (*83*). The final model was evaluated using MolProbity (*84*). Structure deposited to PDB (PDB id 6X9Z). Data collection and refinement statistics are recorded in Table S4.

**Supplementary text:**

General consideration about the β-barrel architecture

We compared the architecture of the previously designed idealized water-soluble β-barrels with naturally occuring TMBs. We found that these two β-barrel architectures of type (n=8, S=10) share structural similarity that can be associated with the canonical constraints on the β-barrel fold, although they fold into very different environments. Both β-barrel architectures have a common orientation that is defined by the unique structural properties of the β-hairpins on either side of the β-barrel. Because of the chirality of the β-turns, we previously found that the β-strand residues flanking the turns on the bottom side of the water-soluble β-barrels (defined as the side with the N- and C-termini) point towards the surface of the barrel while the β-strand residues flanking the turns on the top of the β-barrel point into the core. Additionally, the β-turns on the two sides of the β-barrels are subject to different constraints on their local twist; the register shifts between each β-hairpin at the bottom of the barrel occur between each β-hairpin and the previous one while at the top they occur between each β-hairpin and the following one. Following these principles that are mostly dictated by the chirality of natural amino acids, the orientation of the TMBs can be easily matched to the orientation of the water-soluble β-barrels.
The bottom side of water-soluble β-barrels structurally match the periplasmic side (*cis* side) of TMBs; therefore the extracellular (*trans*) side of TMBs corresponds to the top side of water-soluble β-barrels. We also found similarities in the function of each side of the barrel in both architectures. The bottom side contributes to stability and/or folding. In water-soluble β-barrels, it is often packed with hydrophobic side-chains and features a capping motif with a tryptophan corner critical to folding the protein (*5*, *85*, *86*). The bottom (*cis*) side of the TMBs feature mostly short β-turns with strongly defined β-turn sequences which might be critical for folding since these interactions form early on in the folding pathway (*87*). However, TMBs lack a tryptophan corner folding motif between the first and the last strand by contrast to the water-soluble β-barrel. This difference is discussed later in the supplementary text.

The top side of many water-soluble β-barrels have evolved to support a ligand-binding or catalytic function. To support that function, the core of the β-barrel on the top side is often carved to accommodate the active site and the top β-hairpins are connected with longer loops contributing to the function. TMBs also often feature long and disordered loops on the top (*trans*) side that support many of the functions attributed to the TMBs. These similarities suggest that structural constraints intrinsic to the β-barrel fold could shape the folding and the stability/function trade-offs in both water-soluble β-barrels and TMBs.

<u>Rationale for designing blueprints and hydrogen bonding constraints for β-barrels</u>

The relationship between the number of strands (n) and the shear number (S) of a β-barrel is explained in the main text and illustrated in Table S1. This supplementary material aims to describe a logic to apply to automatically generate blueprints and constraint files for idealized up-and-down β-barrel backbones connected with short β-turns on the *cis* and *trans* sides. We previously showed that the β-sheet of β-barrels with the architecture (n=8, S=10) is strained due to the structural constraints of the hydrogen bonds and the tight packing of core residues (*5*). We described simple rules to design strain-free backbones by introducing glycine kinks at strategic positions in the Cβ-strips and associating each bottom β-turn (or *cis* β-turn in TMB) with a classic β-bulge at position -2 on the first β-strands; and each top β-turn (or *trans* β-turn in TMBs) with a G1 β-bulge. As a simple rule-of-thumb to relieve the clashes within the β-strip in the core of the β-barrel, we do not allow more than four side chains in a row in each Cβ-strip. The row of side chains is interrupted by placing a glycine kink (which lacks a side chain) or a register shift (interruption of the hydrogen bond pattern). The four side chain rule originates from two observations: (i) exceptions to this rule are rare in naturally occuring β-barrels of 8 strands and a shear number of 10; (ii) in the β-barrel architecture (n=8, S=10), the vector spanning four residues in the direction of the hydrogen bonds (along the Cβ-strip) and projected to on the plane perpendicular to the main β-barrel axis has a norm of approximately 12.5 Å (equation 4); which represent a quarter of the ideal β-barrel circumference (calculated based on the ideal radius obtained from equation 1). To understand the effect of the number of side chain in a row along a Cβ-strip, it is useful to think about the β-barrel cross-section along the main axis as a 2D geometric shape - where glycine kinks form geometric corners connected by straight lines (which are the rows of side chains in the core Cβ-strips, assuming that the clashes between those side chains favor straight β-sheets). Every additional side chain in a row along a Cβ-strip will increase the length of one side by approximately 3 Å. We reasoned that the a β-barrel cross-section with one long side might be unfavorable because (i) the additional length would have to be accommodated with acute angles which might result in more strain on the glycine kink corners (ii) the increase of the length of one side above 12 Å would result in a decrease of the volume in the core of the β-barrel, which could lead to difficult core pack and to more side chain clashes. It is, however, important to note that the principles above do not apply to other β-barrel architectures. The strain in the β-sheet of larger β-barrel architectures and the effect of glycine kinks require more investigation that were out of the scope of this work.

We further defined ideal β-barrel topologies in the context of membrane-associated architectural constraints. A basic assumption of the provided guidelines is that the entire β-barrel is embedded in the membrane. Hence, the transmembrane span of a β-strand is defined as the number of residues between the *cis* and *trans* anchor residues (z). The distance between these two surface residues (z x d; where d is the average distance between two Calphas along a β-strand of 3.3 Å) is projected on the main axis of the β-barrel to calculate the transmembrane span Z (equation 7, where theta is the angle of the strands to the main axis).

$$Z = zd\cos(\theta) \qquad \text{eq. 7}$$

For a β-barrel of architecture (n=8, S=10), a β-strand of 11 residues (z=10) will have a transmembrane span Z of approximately 24.1 Å, which is similar to the transmembrane span of TMBs in the outer membrane of *E. coli*.

Once the length of the transmembrane region of the β-strands has been calculated to match the desired transmembrane span, the total length of each β-strand has to be adjusted to satisfy structural constraints related to the β-barrel architecture. For an ideal β-barrel with as constant as possible distribution of the register shifts, there are several considerations:

(i) The previously described principles of ideal β-strand connections (*21*) state that, for strands connected by short β-turns, the residues flanking the β-turns must form a hydrogen bonded pair. In the context of the β-barrel, this rule implies that the edge residues on *cis* hairpins point to the surface of the β-barrel (they are the *cis* anchor residues) while the edge residues on *trans* hairpins face the core of the β-barrel. Since the transmembrane span of the β-strands is calculated from the *cis* and *trans* anchor residues, which are both surface-exposed, the length of each β-strand in the β-barrel is increased by one residue on the *trans* side.

(ii) To accommodate the β-bulges at the *cis* side of the β-barrel, the lengths of the β-strands with an odd number must be increased by one residue.

(iii) Because of the up-and-down sequence of β-hairpins and of the tilt of the strands to the β-barrel axis, the odd-numbered strands are shorter than than the even-numbered strands by two residues.

(iv) In the case of a β-barrel architecture (n=8, S=10), the β-strands length has to account for two additional register shifts between *cis* and *trans* hairpins as described in the main text. Assuming that the additional register shifts in cis happens after the β-strand N (which must be an odd number), the lenght of the β-strands N+1, N+2 and N+3 must be increased by two residues.

To summarize, the β-strand lengths of an ideal β-barrel architecture (n=8, S=10) with a β-bulge residue associated to every *cis* β-turn, a transmembrane beta-strand span z and two additional register shifts after the beta-strand N can be calculated as followed:

- Length of odd beta-strands: z
- Length of even beta-strands: z+2
- Length of the odd beta-strand N+2 : z+2
- Length of even beta-strands N+1 to N+3: z+4

For example, a β-barrel with a transmembrane span of 24 Å (z=10) and two additional register shifts after strand 1 (N=1) would be described with the following topology:

$$\mathbf{E_1}10\text{-}\mathbf{L}3\text{-}\mathbf{E_2}14\text{-}\mathbf{L}2\text{-}\mathbf{E_3}12\text{-}\mathbf{L}3\text{-}\mathbf{E_4}14\text{-}\mathbf{L}2\text{-}\mathbf{E_5}10\text{-}\mathbf{L}3\text{-}\mathbf{E_6}12\text{-}\mathbf{L}2\text{-}\mathbf{E_7}10\text{-}\mathbf{L}3\text{-}\mathbf{E_8}12$$

The constraints describing each backbone hydrogen bond were defined starting from the β-turns. In the absence of a β-turn to guide the strand pairing between the first and the last strand in the β-barrel, the register between these two strands was manually defined to match the desired shear number S. In an ideal β-hairpin connected with a short β-turn (less than six residues long (*21*)), the last residue on the first β-strand and the first residue on the second β-strand form a hydrogen-bonded pair. One hydrogen bond constraint was designed between the backbone amide of the last residue on the first strand and the backbone carbonyl of the first residue on the second strand (the β-turn flanking residues). For two-residues β-turns (*cis* side of the β-barrel), a second hydrogen bond was designed between those two residues. For three-residues β-turns (*trans* side of the β-barrel), the second hydrogen bond was designed between the backbone carbonyl of the last residue on the first strand and the third residue in the β-turn, consistently with the hydrogen bond pattern characteristic of the 3:5 type I β-turn with a G1 β-bulge. Since antiparallel β-strands are characterized by alternating pairs of residues sharing two hydrogen bonds and pairs of residues without hydrogen bonds, two hydrogen bond constraints were designed between every second pair of residues while moving away from the β-turn flanking residues until the end of one of the β-strand. To introduce a β-bulge, an additional hydrogen bond constraint was designed between the backbone amide of the β-bulge residue and the backbone carbonyl of the residue on the neighbour strand forming two regular hydrogen bonds to the residue that follows the β-bulge. The next closest residues forming a hydrogen bonded pair are two positions upstream of the β-bulge residue and two positions downstream of the residue that follows the β-bulge.

<u>Aromatic girdle motifs placement</u>

The presence of motifs that delimit the *cis* and *trans* boundaries of the lipid membrane leaflets has been previously demonstrated (*88*, *89*). We derived a pattern for the *cis* and *trans* aromatic girdles, based on observations of naturally occuring TMBs and the analysis of the constructed MSA for homologous β-barrels of 8 β-strands.

On the *cis* membrane boundary, we found a strong signal for tyrosine at the third position from the end of the strands with even numbers (β-strands in the *cis* hairpins). The frequency of the tyrosine amino acid is as high as 50% at these positions in the MSA. The second most abundant amino acid is phenylalanine, with only 10% frequency (Fig. S6E). Inspection of crystal structures of naturally occuring TMBs confirmed this trend and showed that these tyrosines specifically adopt a *t* rotamer so that the phenolic hydroxyl on the tyrosine side-chain points towards the *cis* water/lipid membrane boundary. To compensate for the four-residue register shift in the β-barrel architecture (n=8,S=10), we placed an additional tyrosine at the second position of the β-turn preceding the large change of register.

Tyrosine was also the most abundant amino acid at the *trans* membrane anchor positions (last position of the first β-strand in the *trans* hairpins), although the preference was not as clearly marked (25% tyrosine frequency, Fig. S6F). The tyrosine side-chain again adopts the specific *t* rotamer in crystal structure to point toward the *trans* water/lipid membrane boundary. In the crystal structures, the tyrosine often interacts with an asparagine residue located two positions up the neighbor strand. We designed two types of motifs on the *trans* side of the TMBs alternating between the β-hairpins: i) a tyrosine at the last position of the first β-strand interacting with an asparagine at the third position of the β-turn (G1 bulge position); ii) a tyrosine at the third position from the end of the first β-strand interacting with an asparagine at the first position of the second β-strand; as well as a tryptophan residue at the last position of the first β-strand involved in an aromatic stacking interaction with the tyrosine. The tryptophans were introduced to facilitate biophysical characterization of the designs based on intrinsic fluorescence.

<u>Glycine and proline residues placement</u>

Previous computational design work on the lipid-exposed surface of tOmpA revealed the key role of surface glycine and prolines in TMBs (*41*). However, the exact positions and mechanism by which such residues, which are generally destabilizing to β-strands, can enable TMB folding is unknown. In the main text, we describe the hypothesis made to place surface glycine and proline residues in the designs. The rationale is described in more details in the text below.

*Glycines in the β-sheet*

The glycines in positions facing the core of the barrel - the glycine kinks - were placed in a strategic way to relieve the strain in the β-sheet and shape the β-barrel lumen as described in a previous paragraph. It is worthwhile to note that the rationale proposed here implies that the number and positions of glycine kinks depend on the strain in the β-sheet and will therefore be different for different β-barrel architectures. The exact relationship between the number and position of glycine kinks, the number of strands in the β-barrel and the shear number requires more investigation.

The high frequency of glycine residues on the surface of TMBs is in striking contrast to water-soluble β-barrels, where solvent-exposed glycines on protein surface are rare. We found a conserved glycine residue on the surface of streptavidin (G74 on PDB structure 1STR), but that position is not solvent-exposed but rather buried amidst a dimerization interface. More examples of surface glycines located at dimerization interfaces are provided by the PDBs 2OVS and 5EE2. Excluding glycines involved in non-canonical β-turns or β-bulges, we found only one solvent-exposed glycine on the surface of the PDB 4REV (G175). These very limited data, together with the high contribution of tight aromatic-to-glycine packing interactions in the core to protein stability ("aromatic rescue" (*28*)), suggest that water-exposed glycines in β-sheet are energetically unfavorable but can be stabilized by hydrophobic interactions. We therefore hypothesized that surface glycines in the β-sheet might be less unfavorable in the hydrophobic environment of the lipid membrane and that the extended torsional space accessible to the glycine amino acid might be able to compensate for the out-of-plane hydrogen bond geometry of glycine kink residues.

*Prolines in the β-sheet*

Two proline residues were introduced into the TMB designs for different purposes. Pro83 has a similar role to the prolines that were placed in our previous water-soluble β-barrel designs. It was designed in the middle of the longest edge-strand resulting from the 4-residue register shift at the *cis* side of the β-barrel and aimed to protect the edge strand from non-desired strand-strand associations and re-enforce the designed shear number and topology.

Pro67 was associated to the mortise/tenon motif located in the β-sheet region between the 4-residue *cis* and *trans* register shift. We previously observed that, in naturally occuring TMBs, several tyrosines in mortise/tenon motifs are preceded by a proline creating a disruption of the hydrogen bonding pattern in the middle of the β-sheet. We hypothesized that the proline could have a similar role to the surface glycine, relieving the frustration associated with out-of-plane hydrogen bond geometry of the glycine-tyrosine pair and the hydrophobic environment of the lipid membrane. We relaxed TMB design models with and without a proline at position 67 associated with the Tyr68 that forms a mortise/tenon motif with Gly88. We found that in the presence of Pro67, Gly88 adopts a more extended conformation characterized by more negative psi torsion angles and out-of-plane hydrogen bonds (Fig. S29D,E). To check whether the more extended glycine kink conformation stabilizes the mortise/tenon motif, we analyzed the Rosetta energy of Tyr68 and Gly88 and found that both residues have in average lower `total_score` in the models relaxed in the presence of Pro67. Tyr68 had an average lower `fa_dun` score, indicating that the rotameric state in the motif was stabilized (Fig. S29A, B).

Mortise/tenon folding motifs

We previously found that the key to the design of water-soluble β-barrels was the strategic placement of specific folding motifs to ensure correct association between β-strands that have ambiguous register definition (such as the interaction between the first and the last β-strands in an up-and-down β-barrel). The tryptophan corner motif was found to tie together the first and last strands of the β-barrel, the longest-range set of interactions and which register is not defined by β-turns. Mutations of the residues belonging to the tryptophan corner into alanine resulted in the failure of the protein to fold into a monomer (*5*). The tryptophan corner motif is absent from TMBs. The putative folding motifs is the mortise/tenon (*29*), which was described as a core tyrosine adopting a +60,90 rotamer to closely interact with the grove formed by the glycine kink in an aromatic rescue type of interaction (*28*) and can be used to predict strand registry (*89*).

In this work, we used the mortise/tenon in the TMBs designs and made two additional hypotheses regarding the structure and position of the motifs in the protein.

First, we propose to extend the definition of the mortise/tenon motif. The analysis of the generated MSA of homologous sequences to tOmpA showed that the negatively charged residue (aspartate or glutamate) forming a hydrogen bond to the tyrosine is as critical or conserved? as the tyrosine and glycine positions, while the rest of positions involved in the second layer of the polar interaction network are less conserved (Fig. S6C). In naturally occuring TMBs, aromatic residues involved in aromatic rescue interactions appear to be also often stabilized by a cation/pi stacking interaction. However, the cation/pi stacking is an interaction that is poorly captured by Rosetta's energy function and we choose to focus exclusively on the canonical YGD/E motif.

Second, it is unknown which of the ambiguously defined registers in TMBs require a mortise/tenon motif. The topology maps of some naturally occuring TMBs and the positions of the mortise/tenon and comparable motifs are shown in Fig. S6D,E. We tested two different positions for the mortise/tenon motif in our TMB designs. We propose that the first area of the β-sheet to require a mortise/tenon motif is formed of three β-strands located between the *cis* and *trans* four-residue register shifts. In other words, we propose to define ambiguous β-strand registers in TMBs based on uneven distribution of register shifts between hairpins rather than the positions of N- and C-termini as in the water-soluble β-barrels. Indeed, many of the mortise/tenon or comparable motifs were observed in that area in naturally occuring TMBs and the N- and C-terminal interactions do not appear to be critical in TMBs since they can be split and circularly permuted (*90*, *91*)., we tested a second mortise/tenon position associated with a glycine kink located closer to the N- and C-termini in our TMB blueprint and on the opposite side of the β-barrel to the first position. The *de novo* designed TMB sequences have either both or only one of these mortise/tenon motifs.

*Cis* and *trans* β-turns

The design of β-turn sequences is discussed in the main text. Here, we justify the choice of the type of short β-turns (the β-turn backbone conformation and length) used to assemble TMB backbones. These principles are valid for the water-soluble and transmembrane β-barrels, which share similar backbone properties.

We previously showed that β-bulges associated with β-hairpins were necessary to relieve the strain associated with the high curvature of the β-sheet in the β-barrel architecture (n=8, S=10) (*5*). Since the structural environments of the four β-turns on each side of the β-barrel are similar, the same β-turns and β-bulge positions were used throughout the *cis* side as well as the *trans* side. Because of the prefered chirality of ββ connections (*21*) and the hydrogen bond patterns characteristic to β-bulges (*92*), the ideal placement of β-bulges is at position -2 from the *cis* β-turns (preceding the paired β-strand residue at position -1) and position +1 from the *trans* β-turns (preceding and replacing the β-strand residue at position +1, which now shifts to position +2). We previously found that the type I β-turn (with the ABEGO type sequence AA) is prefered when a β-bulge is located in position -2 (*5*) and used that type of β-turn to connect *cis* β-hairpins. The *trans* β-hairpins were connected with 3:5 type I β-turns (with ABEGO type AAG) which feature an intrinsic G1 β-bulge at third position (*25*), which modifies the hydrogen bonding pattern of the first residue in the second β-strand. This is equivalent to placing a β-bulge at position +1 from the β-turn, and the 3:5 type I β-turn has been both described as a 3-residue turn and a 2-residue turn followed by a β-bulge (*92*, *93*).

Combinatorial sequence design of set TMB2

The goal of the last set of designs reported here is to increase the hydrophobicity of the core of the TMB designs which will disrupt the alternation of polar and hydrophobic residues along the β-strand and reduce the β-sheet propensity. In short, we started from the mortise/tenon motifs and grew second shell polar interactions to stabilize the tyrosine rotamers. Hydrophobic residues were packed in patches between the resulting polar networks.

To achieve this result, we introduced the tyrosines early in the design process at the first stage of full-atom backbone refinement. Based on our extended definition of the folding motif (YGD/E), we used Rosetta HBNet (*39*) to exhaustively search all the positions that can accommodate a negatively charged aspartate or glutamate residue acting as hydrogen bond acceptor to the tyrosines. The YGD/E motifs identified on each backbone were recombined to generate all the possible combinations of one or two motifs per design. We further ran three additional iterations of combinatorial sequence design that aimed to grow second-shell polar networks around the YGD/E motifs. For each iteration, the surface and core of the TMBs were designed independently to limit the time necessary to achieve each step and to be able to quickly re-adjust subsequent design trajectories. All amino acids except cysteine, proline and glycine were allowed for the design of the core with backbone movement enabled (the glycine kinks were introduced at the backbone-building stage). Only hydrophobic amino acids and the aromatic girdle residues were allowed for the surface design stage, with backbone movement and core side-chain repacking enabled. After each core or surface design step, the best designs were selected based on metrics describing the quality of the core networks of polar interactions in terms of their size, energy and robustness.

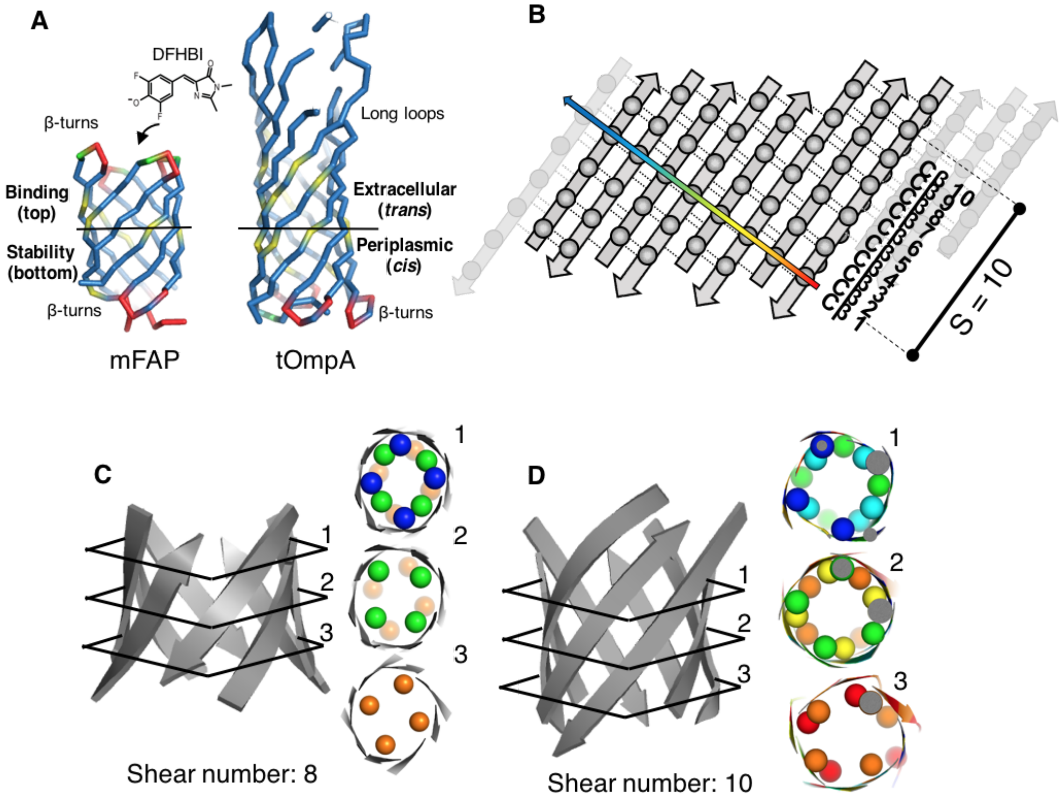


**Fig. S1. Structural constraints on the β-barrel architecture**.

(**A**) Comparison between the overall architecture of the previously reported de novo designed water-soluble β-barrels (mFAPs) and the native tOmpA. Both the water-soluble and membrane protein can be oriented in the same way based on the chirality of the β-strand connections and the location of the N- and C-termini, with the "bottom" of the mFAPs corresponding to the *cis* side of tOmpA; and the "top" to the transmembrane "trans" side. (**B**) The β-barrel architecture is defined by the number of β-strands (N) and the shear number S. S is the number of register shifts along a given β-strand after circling the whole β-barrel in the direction of the hydrogen bonds. S equals the number of Cβ strips in the barrel; half of which point to the β-barrel lumen. (**C**, **D**) The combination of the shear number and of the number of strands define the packing arrangement of side-chains in the core of the β-barrel. For a given number of β-strands (N=8), the core of the β-barrel of type (S=N) is packed with a 4-fold symmetric arrangement of side-chains (**C**, side-chains are represented as spheres and colored according to their position in their respective Cβ-strip). (**D**) The packing symmetry is broken when the shear number is increased by two register shifts so that (S=N+2). The asymmetric arrangement increases the degree of contact between the intertwined side-chains.

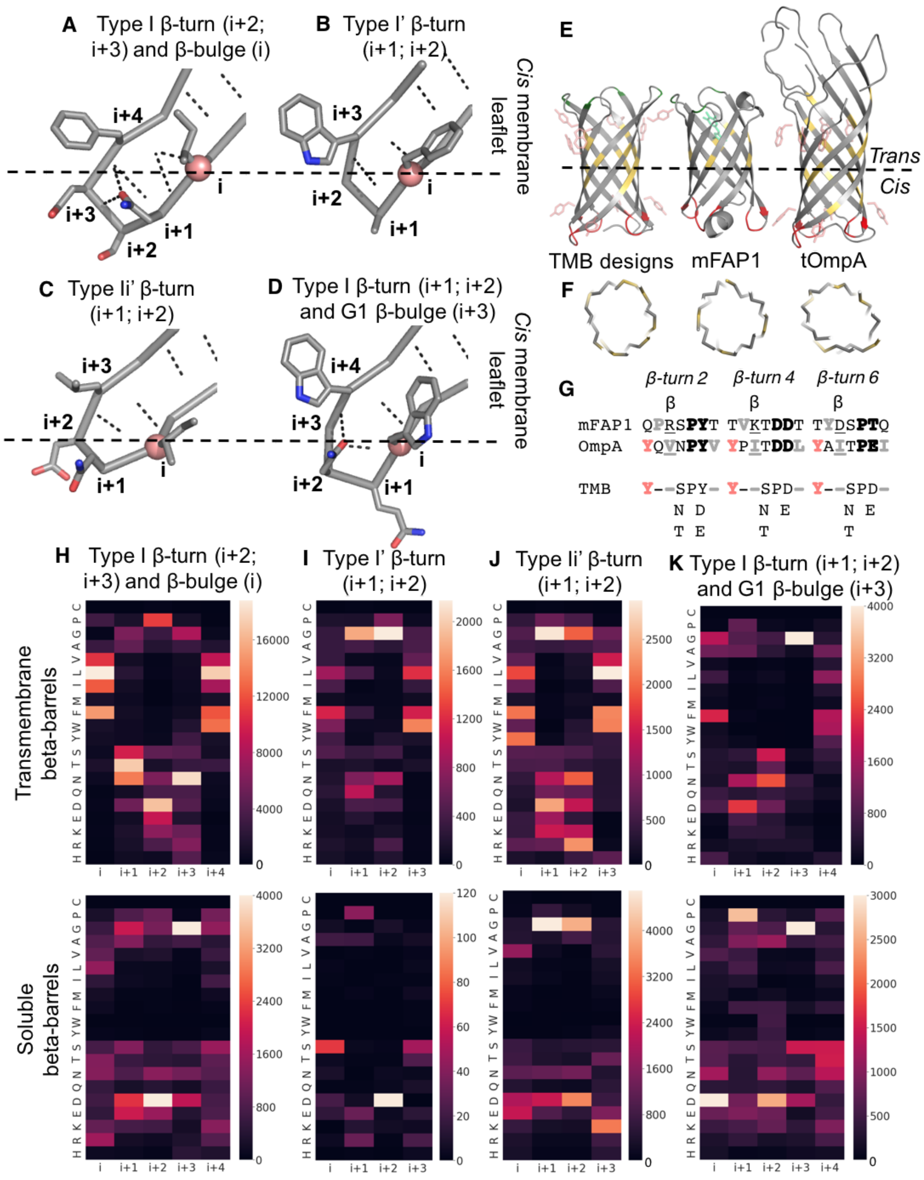


**Fig. S2. Constraints on the structure and sequence of the *cis* β-turns.**

(**A-D**) Representative structures of common canonical type I (**A**), type I' (**B**), type II' (**C**) and type I with G1 β-bulge (**D**) and their membrane context in our model (supported by predictions with the PPM server). The membrane anchoring residue (i) is highlighted with a pink sphere. Hydrogen bond interactions are shown as black dashes. (**E, F**) β-barrel architecture (**E**) and cross-section (**F**) comparisons between the TMB de novo designs, the previously reported soluble *de novo* designed mini Fluorescence Activating Protein 1 (mFAP1) and the transmembrane domain of the native Outer Membrane Protein A from E. coli (tOmpA). (**G**) Comparison of the *cis* β-turn sequences in tOmpA, in mFAPs and concensus used for TMB design. The β-turn residues are shown in bold, the β-bulge residue is underlined, the tyrosine of the aromatic girdle is red and hydrophobic residues are shown in grey. (**H-K**) Heatmaps showing the amino acid preference per position for *cis* β-turns with canonical backbone conformations in natural transmembrane and water-soluble β-barrels.

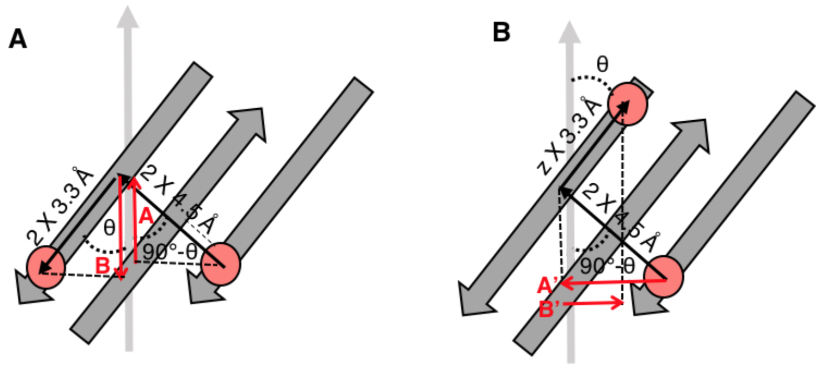


$A = 2 \times 4.5 \times \cos(90 - \theta)$
$B = 2 \times 3.3 \times \cos(\theta)$
Vertical offset = |A - B|

$A' = 2 \times 4.5 \times \sin(90 - \theta)$
$B' = z \times 3.3 \times \sin(\theta)$
Horizontal offset = |A' – B'|

**Fig. S3. Mathematical formula to calculate the vertical and horizontal offset between two residues in the β-barrel as a function of the angle $\theta$ of the β-strands to the main barrel axis**.
(**A**) The vertical offset between two anchor residues on the same side of the β-barrel was obtained by calculating the difference between the vertical offset when moving from strand to strand along the hydrogen bonds (A) and the vertical offset when moving along one β-strand (B). (**B**) The horizontal offset between two anchor residues located on the opposite sides of the β-barrel was obtained by calculating the difference between the horizontal offset when moving from strand to strand along the hydrogen bonds (A') and the horizontal offset when moving along one β-strand (B'). The number of residues z is a function of the desired hydrophobic thickness (Supplementary text). The tilt angle $\theta$ of the β-strands to the main axis of the β-barrel is a function of the parameters n and S (main text).

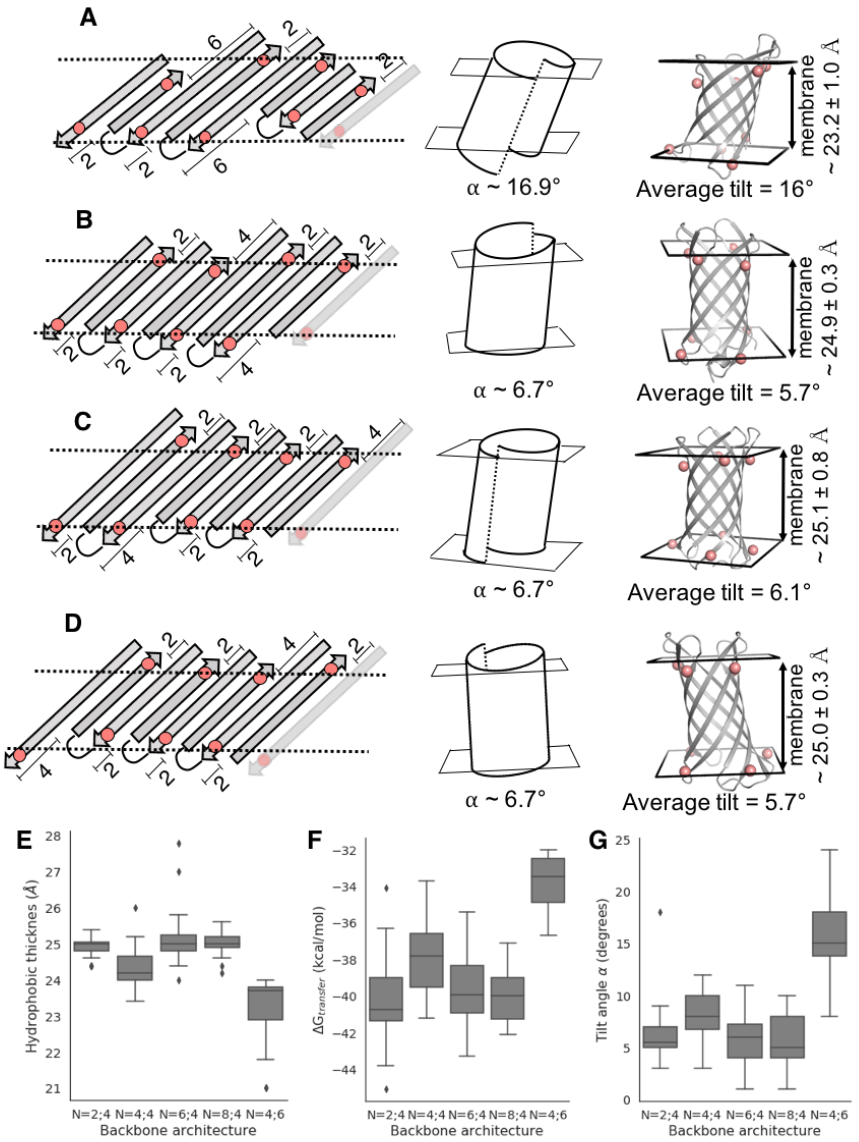


**Fig. S4. Membrane-association constraints on the β-barrel architecture (part 2)**.

(**A**-**D**) Relationship between the topology (left), the geometric model (center) and the Rosetta molecular model coupled with PPM lipid bilayer prediction (right) of four β-barrels with 8 strands and a shear number of 10 and different register shift distributions. From the N- to the C-terminus: (**A**) register shifts 0+6+2+2 (topology N=4;6); (**B**) register shifts 4+2+2+2 (topology N=2;4); (**C**) register shifts 2+2+4+2 (topology N=6;4); (**D**) register shifts 2+2+2+4 (topology N=8;4). (**E**-**G**) Average hydrophobic thickness (**E**), energy of transfer from water to lipid (**F**), and tilt angle to the membrane axis (**G**) predicted by the PPM server on 20-25 Rosetta models. The more uneven distribution of register shifts (**A**) results in a more tilted β-barrel. The topologies (**B**), (**C**) and (**D**) differ only by the positions of the four-residue register shift in the β-sheet. These three topologies and the one presented in the main text (register shifts 2+4+2+2; N=4;4) result in very similar predicted interaction with the lipid bilayer and differ only in the direction of the tilting to the membrane axis.

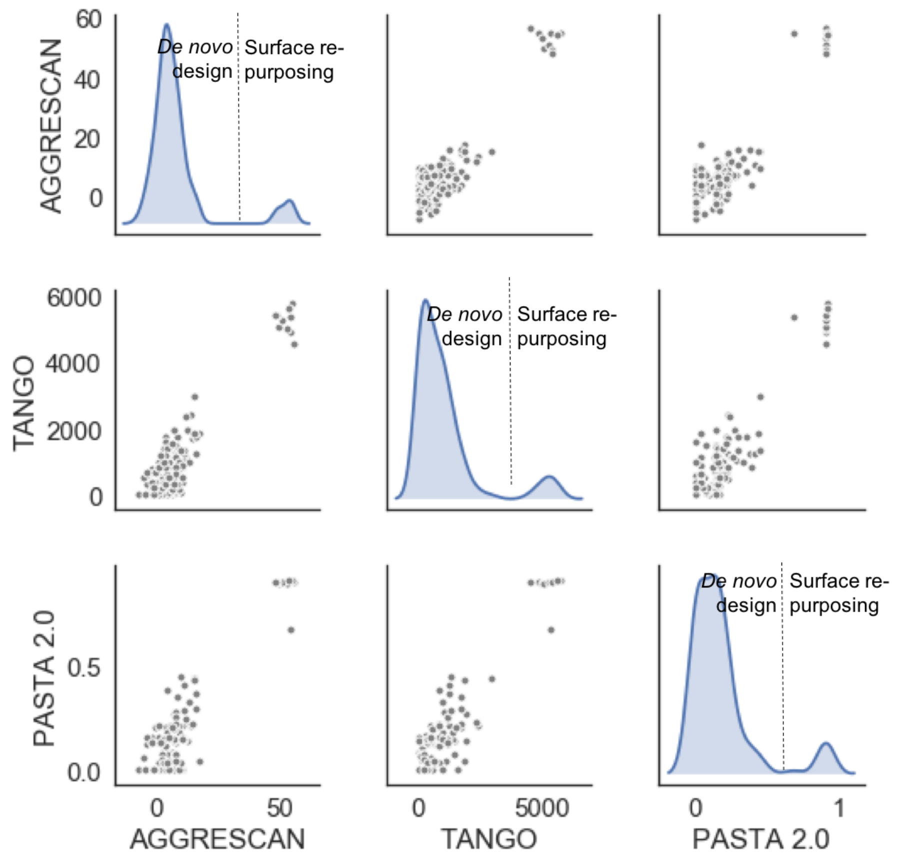


**Fig. S5. The resurface water-soluble β-barrel designs have high aggregation propensity.**

The aggregation propensity of sequences obtained by redesigning the surface of water-soluble β-barrels with hydrophobic residues (surface re-purposing) or designed completely from scratch (*de novo* design) was predicted using PASTA2.0 (*94*), TANGO (*64*) and AGGRESCAN (*95*) prediction servers. All three servers predicted higher aggregation propensity for the "surface re-purposed" designs.

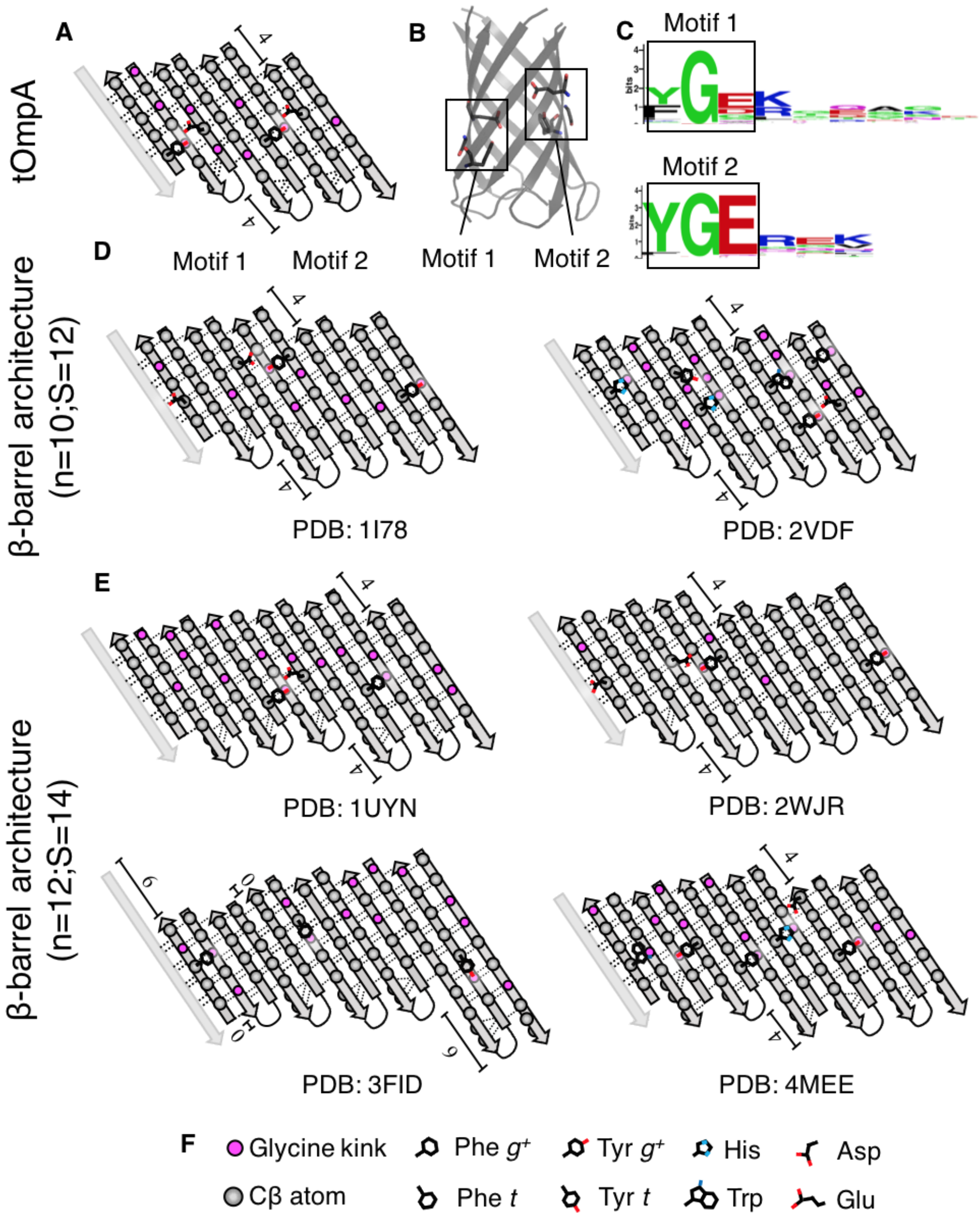


**Fig. S6. Positions of mortise/tenon motifs in some naturally occurring TMBs.**

(**A-C**) Two extended-definition mortise/tenon motifs (YGD/E) found in the native tOmpA TMB mapped on tOmpA topology (**A**) and structure (**B**). (**C**) Weblogo (*96*) representation of the amino acid diversity in the MSA of tOmpA homologs for residues of the YGD/E motifs (black box) and residues from the second shell of polar interactions. (**D**) Putative mortise/tenon motifs identified in two native TMBs with β-barrel architecture (n=10,S=12) and mapped on the 2D representation of the β-barrel topology. (**E**) Putative mortise/tenon motifs identified in four native TMBs with β-barrel architecture (n=10,S=12) and mapped on the 2D representation of the β-barrel topology. (**F**) Legend of the pictograms used in panels (**A**), (**D**) and (**E**).

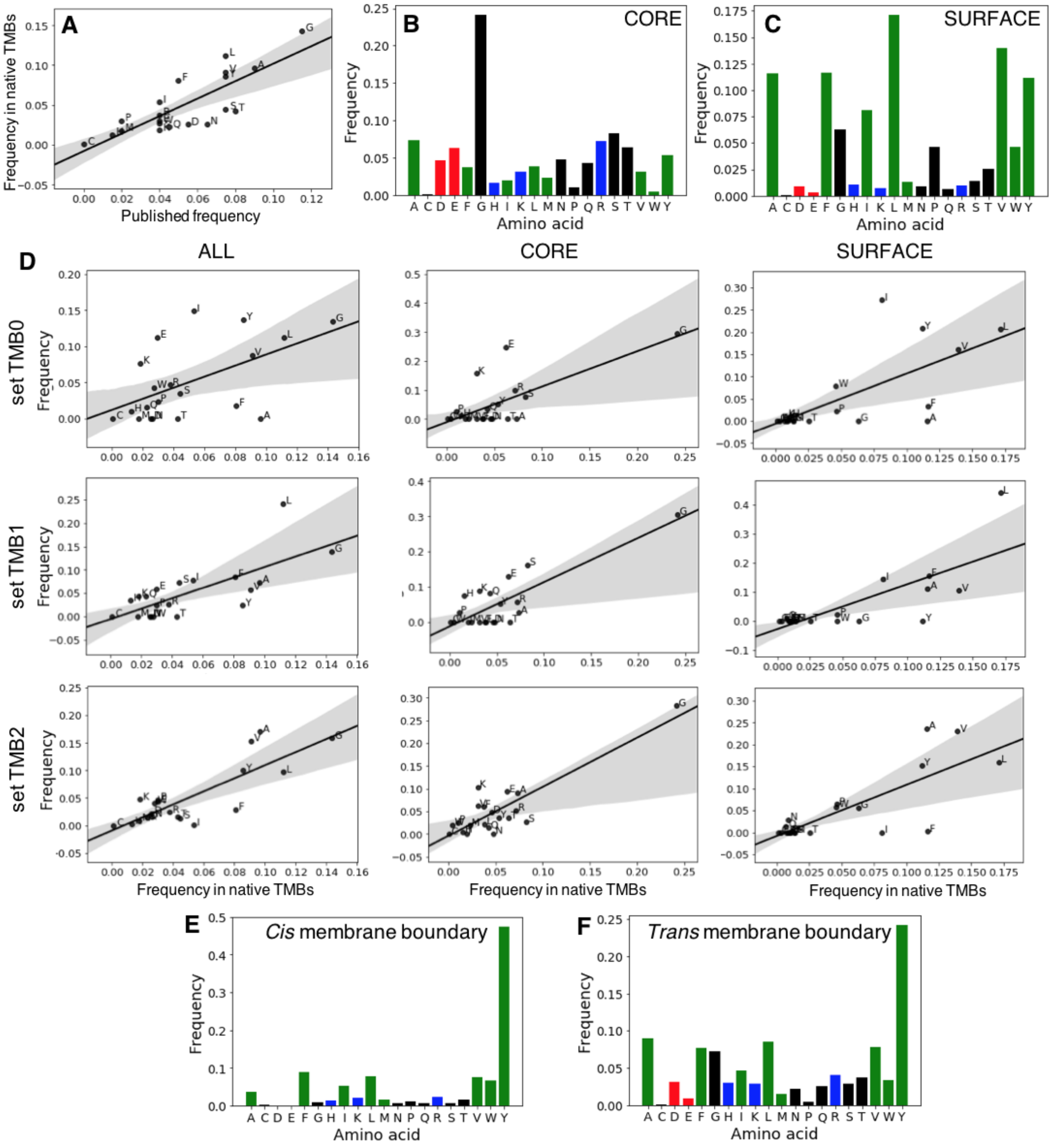


**Fig. S7. Frequency of amino acids in *de novo* MB designs and natural TMBs.**

(**A**) The amino acid frequencies in native 8-strands TMBs derived from the MSAs were validated against previously published frequencies obtained from crystal structures of natural TMBs of different numbers of strands (*8*). (**B** and **C**) Amino acid distributions in the core and on the surface of the reference TMB set. The negatively charged, positively charged, polar and hydrophobic amino acids are shown in red, blue, green and black, respectively. (**D**) Frequency of amino acids in sequences generated in the sets of designs TMB0, TMB1 and TMB2. The distributions are broken down into core and surface positions and compared to the reference set obtained from the MSA in (**A**). (**E** and **F**) Frequency of each amino acid on the aromatic girdle position on the *cis* hairpins (**E**, three positions away from the *cis* β-turn on strand 1) and on the *trans* hairpins (**F**, four positions away from the *trans* β-turn on strand 1).

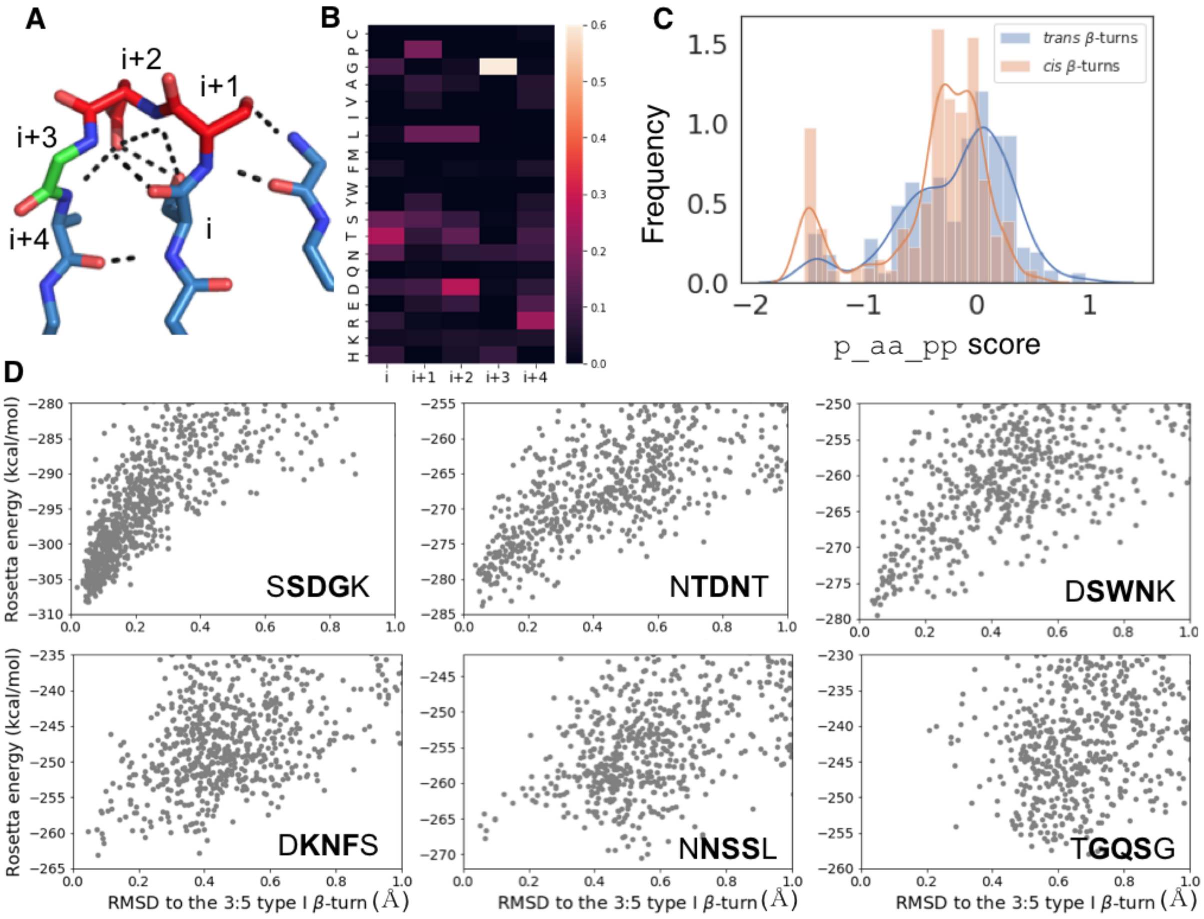


**Fig. S8. Naturally occurring β-turns on the *trans* side of TMBs have sub-optimal sequences for the backbone conformations observed in crystal structures (part 1).**

(**A**) Backbone conformation characteristic of the 3:5 type I β-turn with a G1 bulge. The hydrogen bonds are shown as black dashed lines. The residues in the turn are colored according to which Ramachandran graph binning the backbone phi and psi angles belong to. Blue, green and red correspond to the β-sheet, G-helix (positive phi) and α-helix space, respectively. The residues are numbered from residue i (last residue of the first β-strand) to i+4 (first residue on the second β-strand). Part of the neighbour β-strand is shown on the right. (**B**) Heatmap showing per position amino acid preference in 3:5 type I β-turns fragments extracted from the PDB (and biased toward water-soluble protein statistics). The sequence SDG results in a tight intra-turn hydrogen bond network (**A**). (**C**) Rosetta's `p_aa_pp` scores computed on 100 *trans* and 119 *cis* β-turn residues (two to five-residue β-turns) extracted from 13 crystal structures (Data S3). (**D**) Structure to energy landscape computed with Rosetta loopmodel protocol with KIC (*65*) for the canonical (SSDGK) and sub-optimal β-turn sequences found in natural TMBs. The x axis shows the RMSD of the simulated conformation to the canonical backbone of the 3:5 type I β-turn.

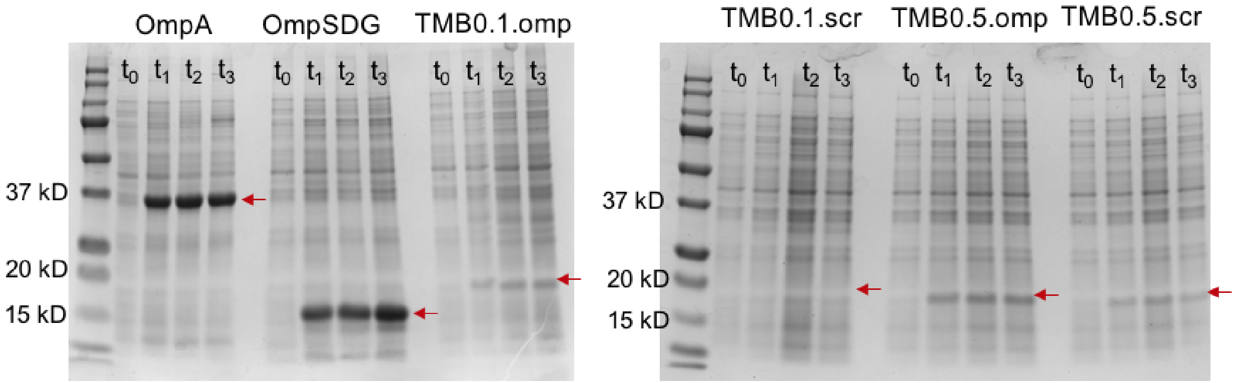


**Fig. S9. Expression gels of designs from set0 with long loops in *trans*.**

SDS-PAGE gels showing whole cells expressing native (full length OmpA and OmpSDG) and designed (TMB0.1 and TMB0.5 which have inserted native loop sequences (.omp) or scrambled loop sequences (.scr) from tOmpA) constructs at t0 (induction), t1, t2 and t3 (one, two and three hours after induction of protein expression). The red arrow shows the expected molecular weight (Mw) for each construct.

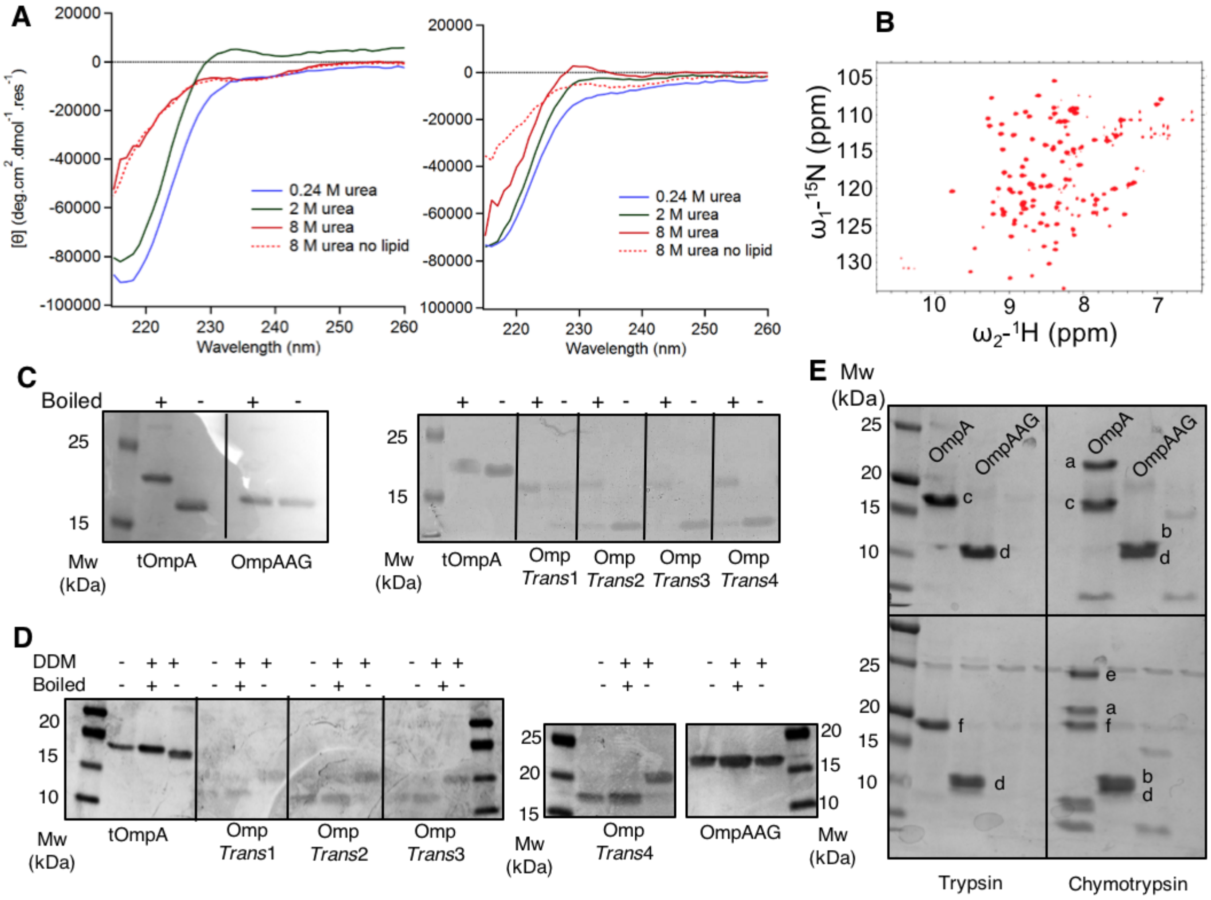


**Fig. S10. Sub-optimal sequences of *trans* β-turns support folding of the native tOmpA TMB while the canonical sequence does not**.

(**A**) Far-UV CD spectra of the native tOmpA and the Omp*Trans*3 construct refolded into DUPC LUVs and in the presence of different concentrations of urea. (**B**) $^{15}N$-$^{1}H$ HSQC NMR spectrum of the Omp*Trans*3 construct refolded into DPC detergent micelles shows well-dispersed resonance peaks characteristic of a folded TMB. (**C**) Band-shift assay on SDS-PAGE gels after refolding native tOmpA and *trans* β-turn variants in synthetic lipid membranes. (**D**) Band-shift assay on SDS-PAGE gel after refolding native tOmpA and *trans* β-turn variants in DDM detergent micelles. (**E**) OmpAAG is protected from protease digestion by the lipid bilayer and therefore forms a species associated with the lipid membrane. The SDS-PAGE gels show the results of a protease challenge (trypsin and chymotrypsin) of full-length OmpA and β-turn construct OmpAAG after refolding into synthetic membranes. Top: gel loaded after protease treatment. Bottom: gel loaded after protease treatment and boiling. The position of band a. matches the expected position of the full-length OmpA protein in its folded state. The position of band c. matches the position expected for the transmembrane domain tOmpA in its folded state.

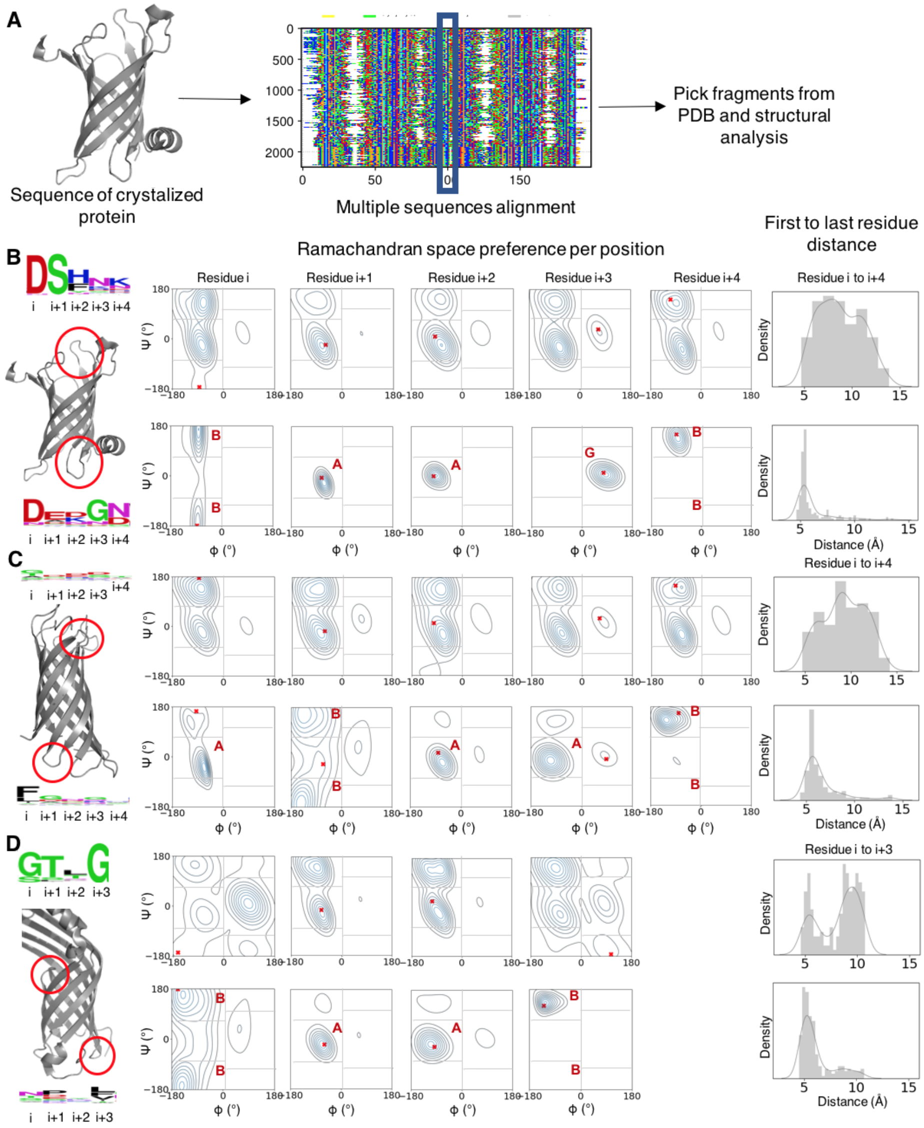


**Fig. S11. Naturally occuring β-turns on the *trans* side of TMBs have sub-optimal sequences for the backbone conformation observed in crystal structures (part 2).**

(**A**) Schematic representation of the computational experiment. Homologous sequences to crystallized TMBs were collected into multiple sequence alignments (MSAs). The resulting β-turn sequence profiles were used to pick fragments from the PDB. We compared the structural properties of the fragments picked based *cis* and *trans* β-turn sequence profiles (**B-D**, Left, represented with Weblogo (*96*)). (**B-D**, Center): Distributions of the Ψ and Φ torsion angles per position (residue i to residue i+3 or residue i+4). The torsion angles observed in the crystal structure are shown as a red cross and the most enriched ABEGO bin is shown in red. (**B-D**, Right): Distributions of distances between the Cα atoms of the first and the last residues. The average distance between the Cα atoms in a β-turn is around 5 Å. We compared the compared the *trans* and *cis* 3:5 type I β-turns in PDB 1THQ (**B**); the 3:5 type I β-turns in PDB 2ERV (our results suggest that many homologs have a different 2-residues type I β-turn with a classic β-bulge in *cis*) (**C**) and the 2-residues type I β-turns in PDB 4FUV (**D**).

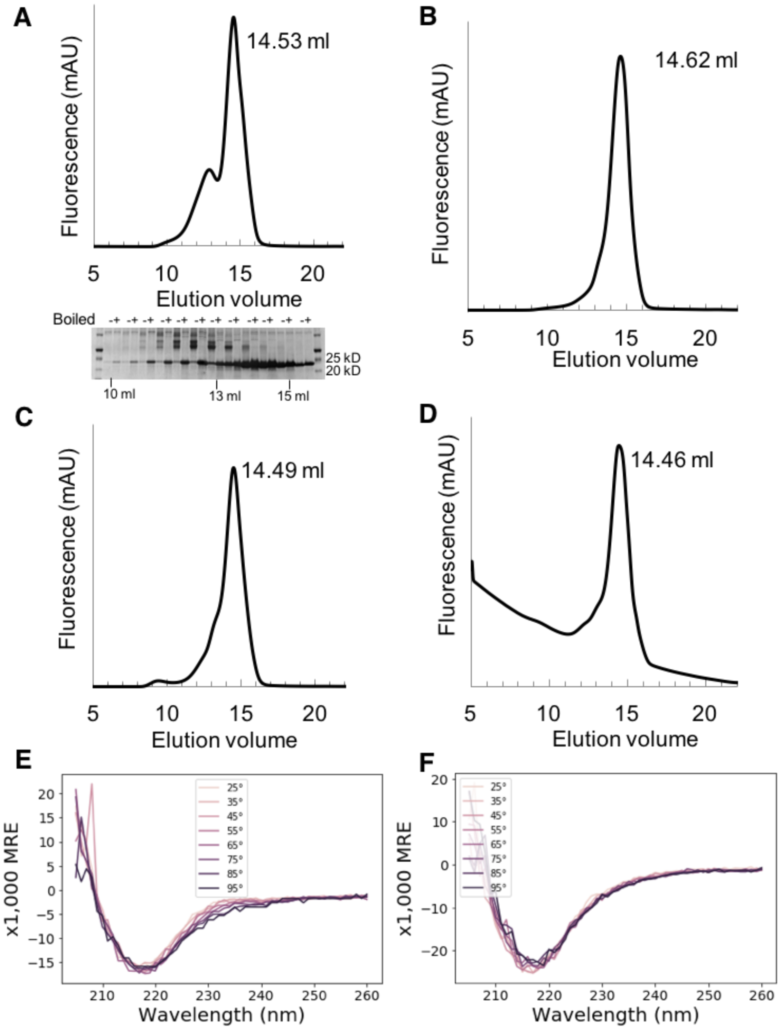


**Fig. S12. Experimental characterization of Omp*Trans* variants of tOmpA.**

(**A**) SEC chromatogram of tOmpA refolded into DDM detergent micelles. The band-shift assay on SDS-PAGE shows the presence of two different heat-modifiable species that match the two major peaks of the chromatogram. The existence of oligomeric OmpA species has been described (*97*). (**B-D**) SEC chromatogram of Omp*Trans*1, Omp*Trans*2 and Omp*Trans*4 refolded into DDM detergent micelles. (**E**) Far-UV CD spectra collected for tOmpA in DDM micelles at temperatures ranging from 25°C to 95°C. (**F**) Far-UV CD spectra collected for Omp*Trans*1 in DDM micelles at temperatures ranging from 25°C to 95°C.

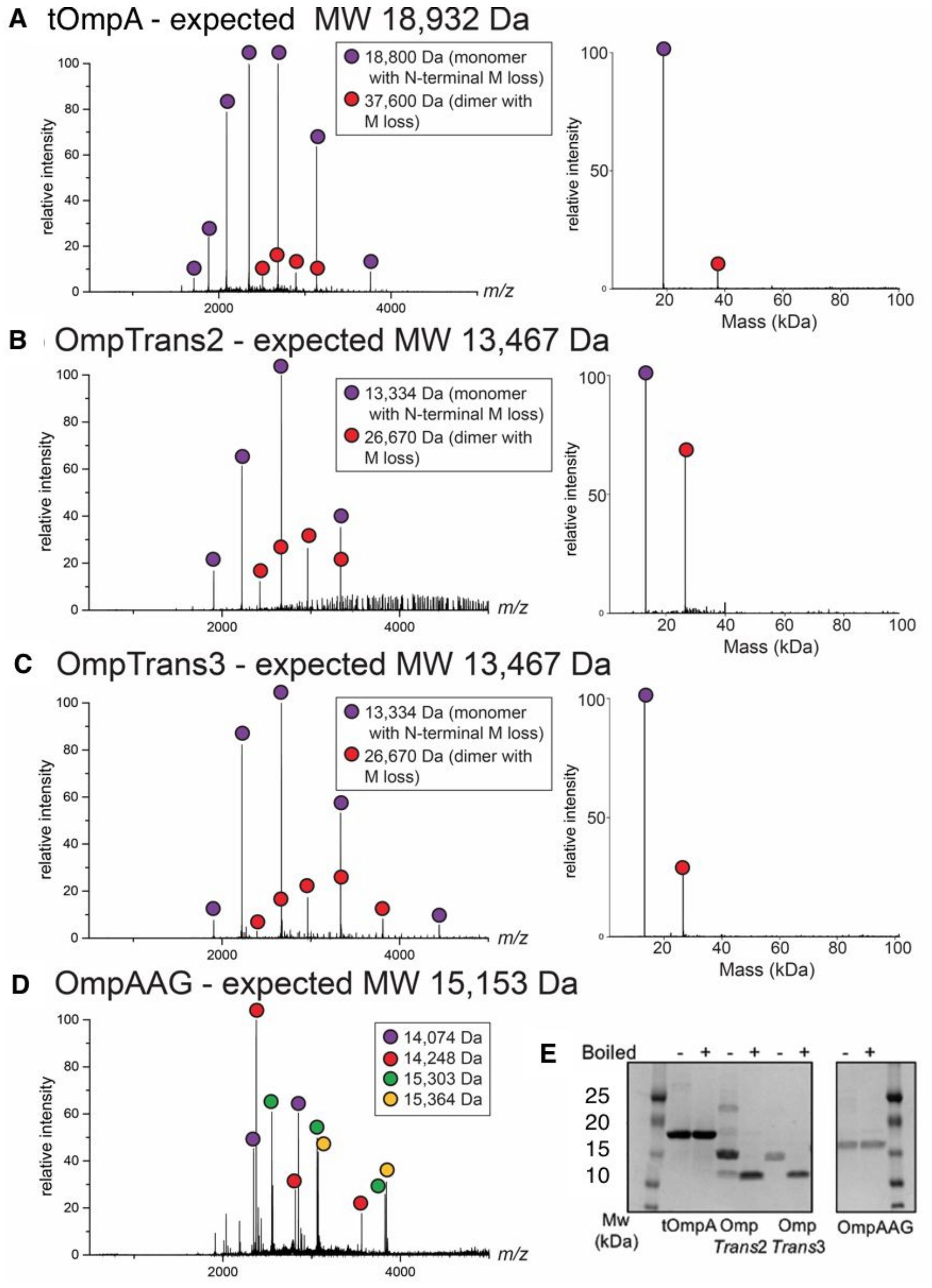


**Fig. S13. Native mass spectrometry (nMS) profiles of tOmpA and *trans* β-turn variants in DDM detergent micelles.**

(**A**) The nMS profile of tOmpA shows the presence of a major monomeric and a minor dimeric species in detergent. (**B**, **C**) Omp*Trans*2 and Omp*Trans*3 show the presence of two species, with a higher dimer to monomer ratio in the Omp*Trans*2 sample. (**D**) The expected mass for OmpAAG was not observed (the sample is probably degrading). (**E**) The SDS-PAGE gel of the samples analyzed by nMS shows the presence of a heat modifiable species for tOmpA, Omp*Trans*2 (with an additional band possibly corresponding to the dimer) and Omp*Trans*3.

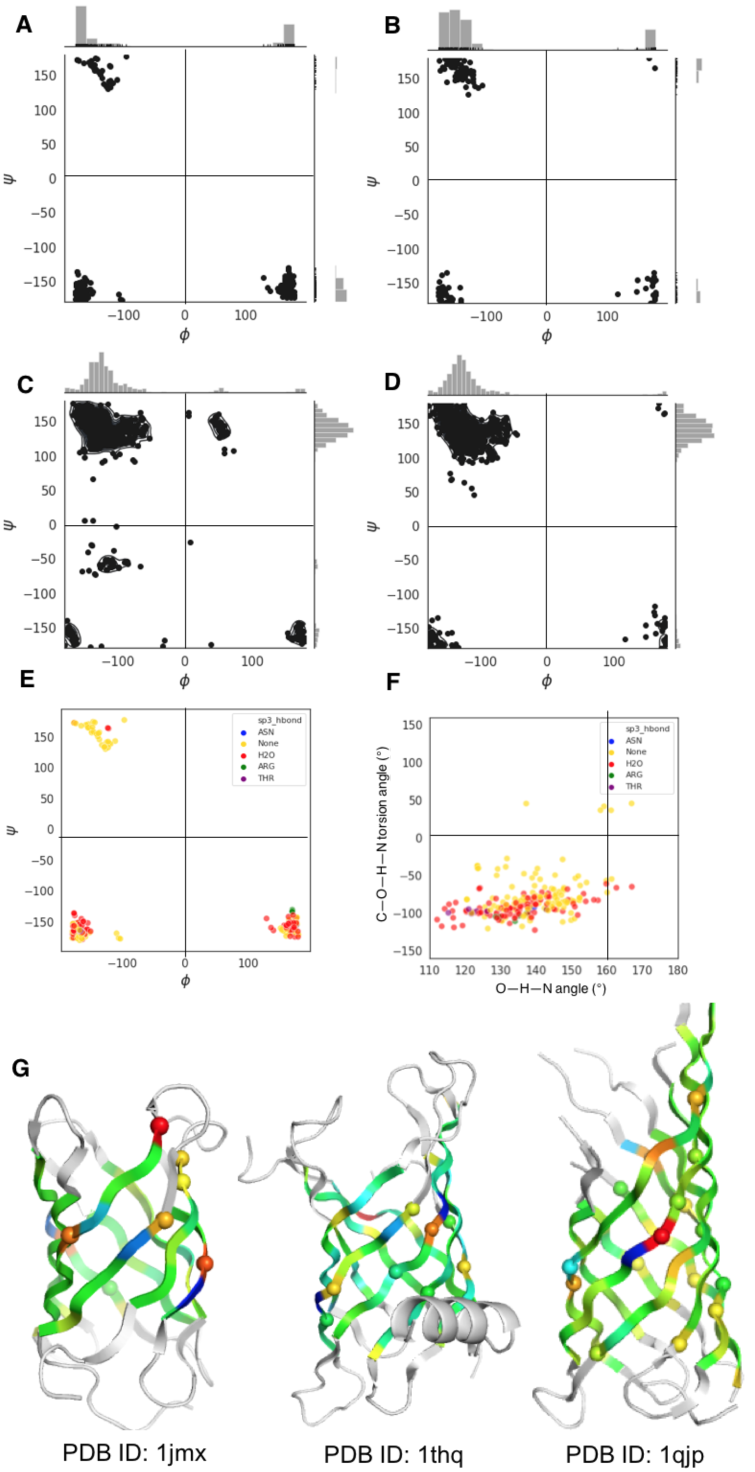


**Figure S14: Backbone and hydrogen bond geometries of β-strand residues in water-soluble and transmembrane β-barrel domains of 8 strands.**

(**A**, **B**) Φ and Ψ backbone torsion angles of glycine kink residues in the core of water-soluble (**A**) and transmembrane (**B**) β-barrel crystal structures. (**C**, **D**) Φ and Ψ backbone torsion angles of all β-strand residues in water-soluble (**C**) and transmembrane (**D**) β-barrel crystal structures. (**E**) $\phi$ and $\psi$ backbone torsion angles of glycine kink residues in the core of crystallized water-soluble β-barrels, colored by the presence or absence of a secondary hydrogen bond donor to the backbone carbonyl. (**F**) Backbone carbonyl hydrogen bond geometries of glycine kink residues in the core of water-soluble crystallized β-barrels. Each data point is colored by the presence or the absence of a secondary hydrogen bond donor to the backbone carbonyl. (**G**) Crystal structures of one water-soluble (1jmx) and two transmembrane (1thq and 1qjp) β-barrel domains, colored by the computed local strand twist per residue (from strong left-hand to strong right-hand twist: red-green-blue).

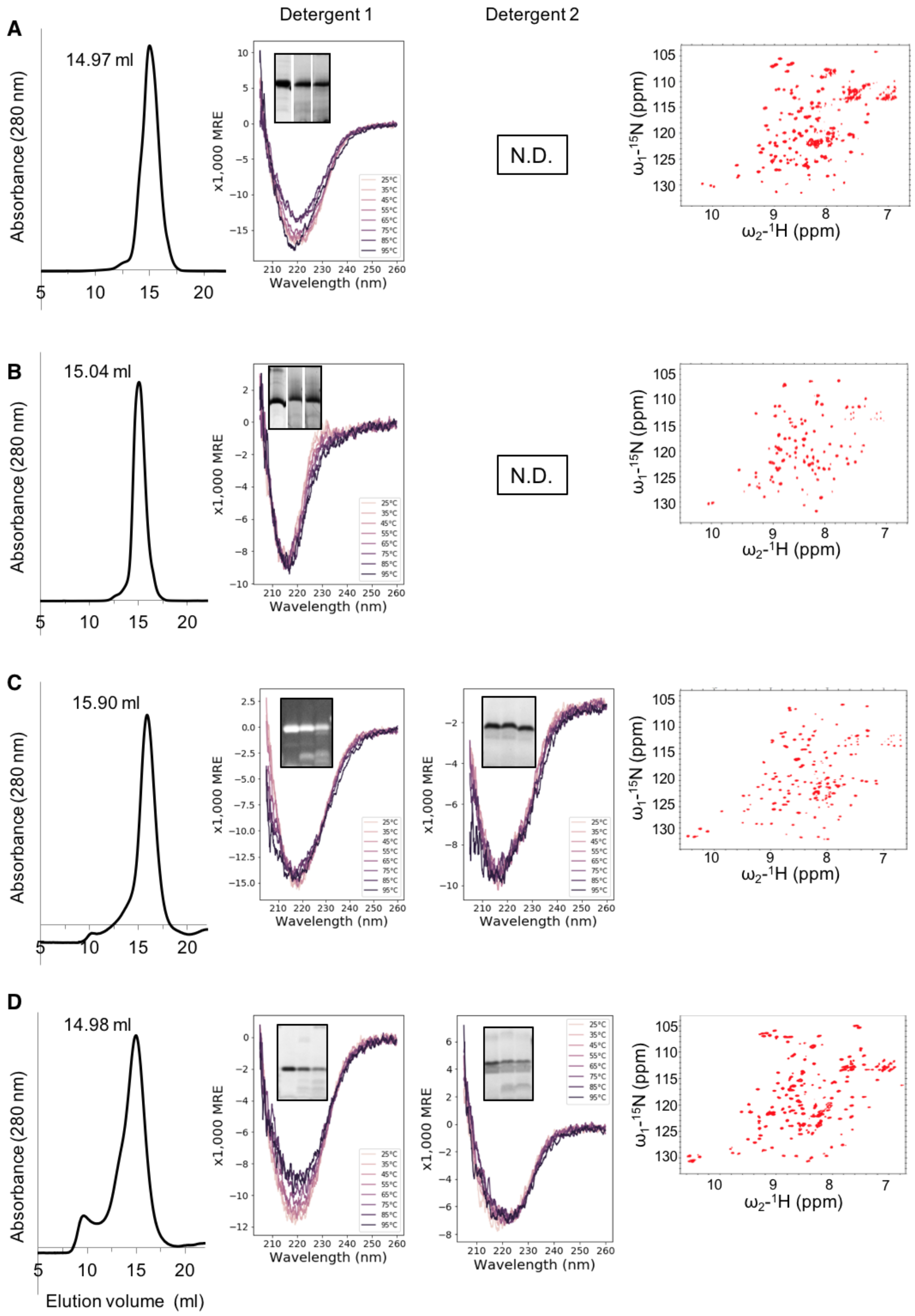


**Fig. S15. Experimental screening data obtained for TMB2 designs that show well-dispersed NMR resonance peaks consistent with a folded TMB (Part 1).**

(**A**) TMB2.3; (**B**) TMB2.17; (**C**) TMB2.35; (**D**) TMB2.58.Left: SEC elution profile of the TMB/DPC complex. Center: Far-UV CD spectra of the TMB/detergent complex at temperatures ranging from 25°C to 95°C in DPC (detergent 1) and OG (detergent 2) micelles. The insert shows a SDS-PAGE gel before (first lane) and after treatment with chymotrypsin (second lane) or trypsin (third lane). Right: NMR resonance peaks collected in DPC detergent micelles.

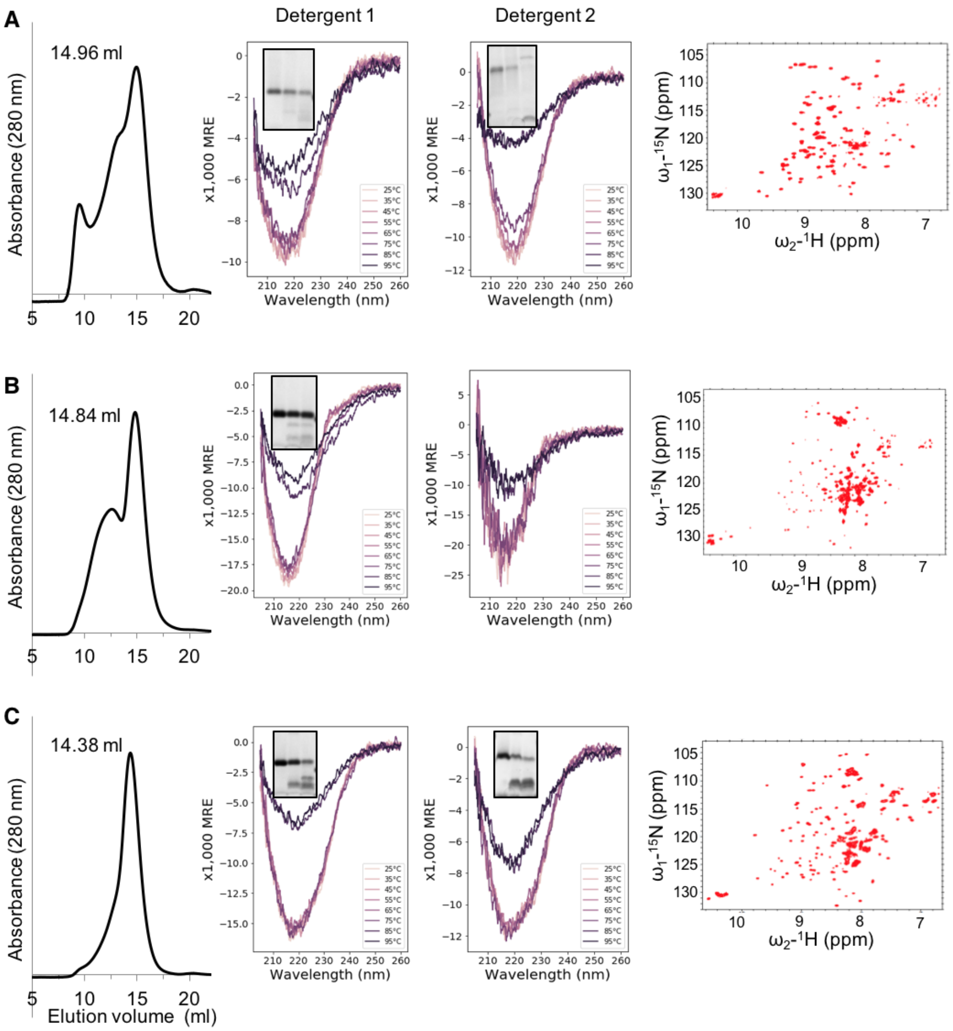


**Fig. S16. Experimental screening data obtained for TMB2 designs that show well-dispersed NMR resonance peaks consistent with a folded TMB (Part 2).**

(**A**) TMB2.69; (**B**) TMB2.70; (**C**) TMB2.73. Left: SEC elution profile of the TMB/DPC complex. Center: Far-UV CD spectra of the TMB/detergent complex at temperatures ranging from 25°C to 95°C in DPC (detergent 1) and OG (detergent 2) micelles. The insert shows a SDS-PAGE gel before (first lane) and after treatment with chymotrypsin (second lane) or trypsin (third lane). Right: NMR resonance peaks collected in DPC detergent micelles.

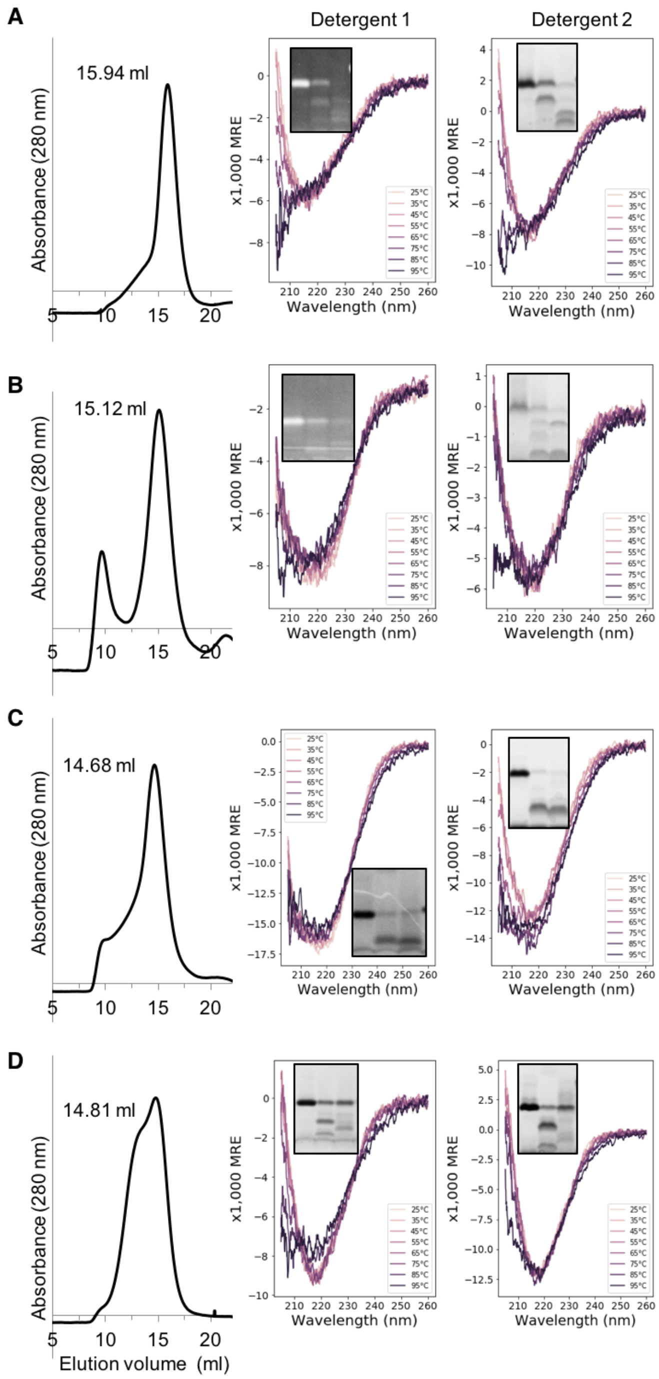


**Fig. S17. Examples of experimental screening data interpreted as negative, obtained for the four TMB2 designs that are unlikely to fold into a TMB structure.**

(**A**) TMB2.32; (**B**) TMB2.42; (**C**) TMB2.48; (**D**) TMB2.76. Left: SEC elution profile of the TMB/DPC complex. Center: Far-UV CD spectra of the TMB/detergent complex at temperatures ranging from 25°C to 95°C in DPC (detergent 1) and OG (detergent 2) micelles. The insert shows a SDS-PAGE gel before (first lane) and after treatment with chymotrypsin (second lane) or trypsin (third lane).

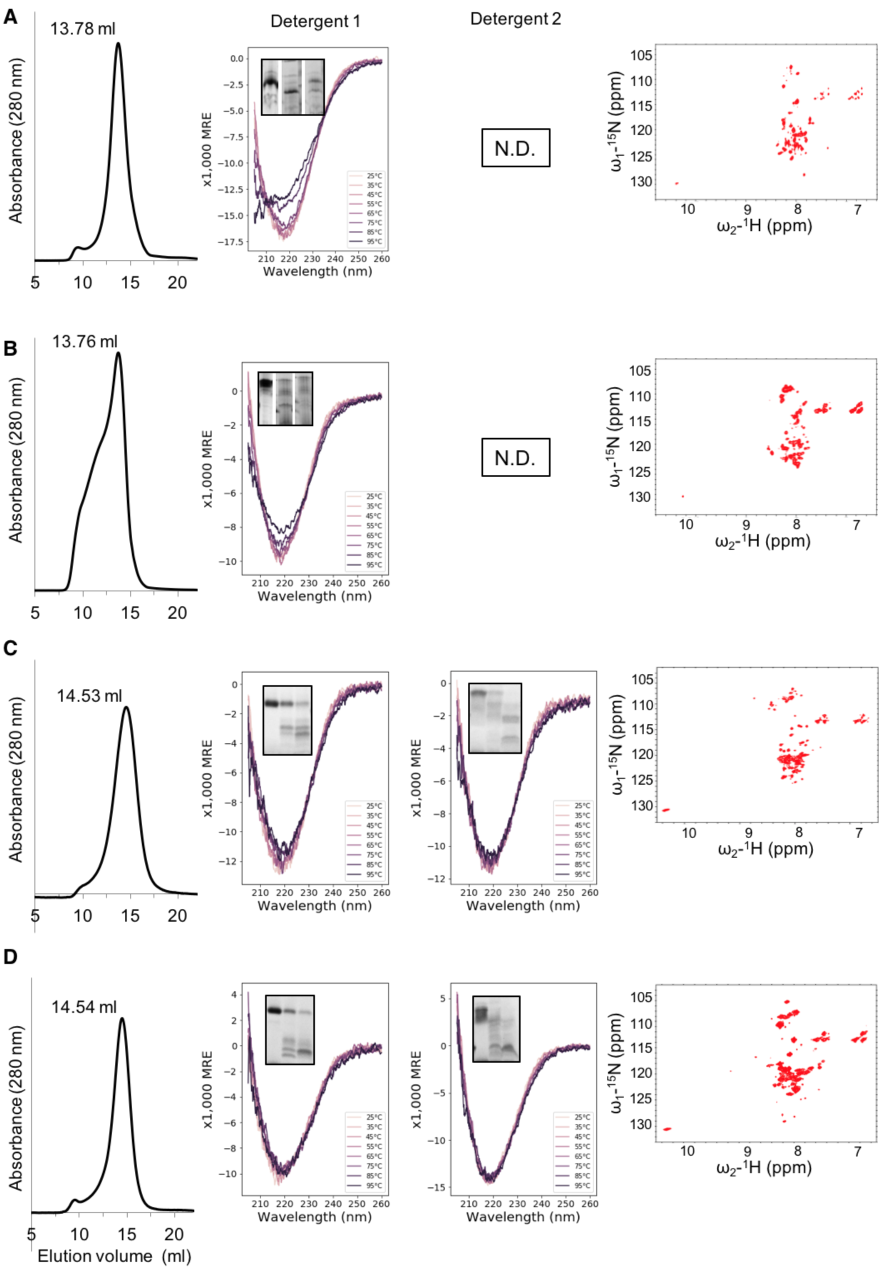


**Fig. S18. Experimental screening data obtained for four TMB2 designs that showed a negative HSQC profile.**

(**A**) TMB2.12; (**B**) TMB2.15; (**C**) TMB2.79; (**D**) TMB2.88. TMB2.12 (**A**) and TMB2.15 (**B**) were re-folded and characterized and analyzed by SEC and CD in DDM detergent. TMB2.79 (**C**) and TMB2.88 (**D**) were re-folded and characterized and analyzed by SEC and CD in DPC detergent. Left: SEC elution profile of the TMB/detergent complex. Center: Far-UV CD spectra of the TMB/detergent complex at temperatures ranging from 25°C to 95°C in detergent 1 (DDM (**A**, **B**) or DPC (**C**, **D**)) and OG (detergent 2) micelles. The insert shows a SDS-PAGE gel before (first lane) and after treatment with chymotrypsin (second lane) or trypsin (third lane). Right: NMR resonance peaks collected in DPC detergent micelles.

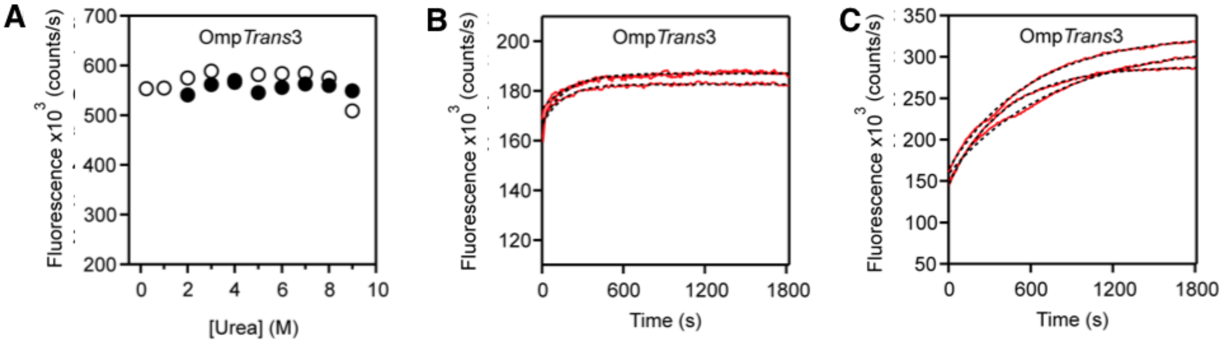


**Fig. S19. Biophysical characterisation of the Omp*Trans*3 variant of tOmpA in synthetic lipid membranes.**

(**A**) Urea dependence of folding and unfolding in DUPC LUVs. The fluorescence intensity at 335 nm was plotted against urea for folding (open circles, dashed line) and unfolding (filled circles, solid line). Omp*Trans*3 is able to fold even in 9 M urea. Kinetics of folding into (**B**) DUPC and (**C**) DMPC LUVs at an LPR of 3200:1 (mol/mol) in 50 mM glycine-NaOH pH 9.5, 2 M urea at 25 °C monitored by tryptophan fluorescence at 335 nm over 30 minutes (red line). Data were fitted with a single exponential function to determine folding rate constants (black dashed line). Three replicates are shown for each.

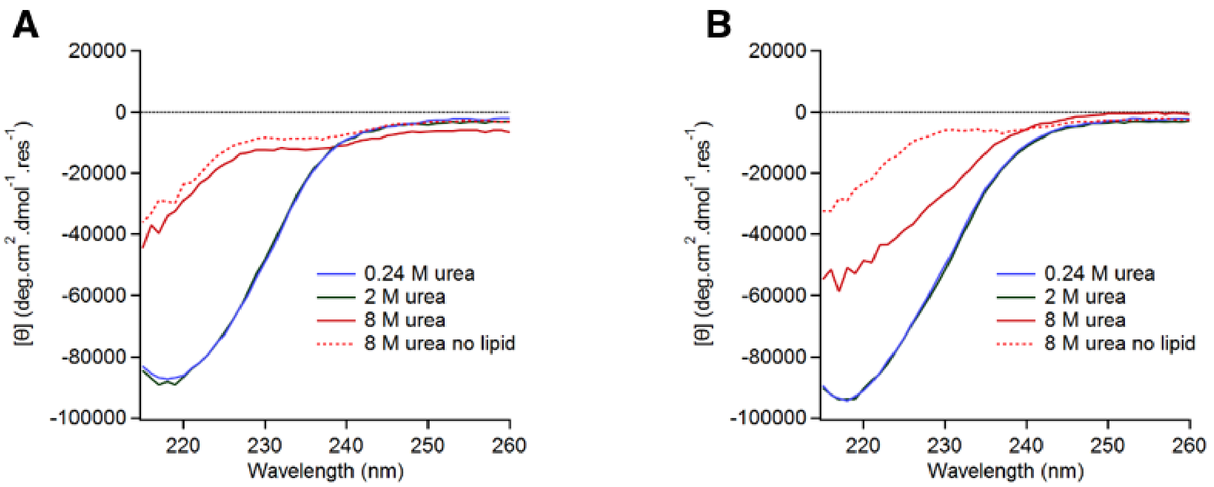


**Fig. S20. Designed OMPs have β-sheet secondary structure.**

Far UV CD spectra of (**A**) TMB2.3 and (**B**) TMB2.17 refolded overnight at 25°C in DUPC LUVs in 50 mM glycine-NaOH pH 9.5 containing 0.24 M urea (blue), 2 M urea (green) or 8 M urea (red). A spectrum was also acquired in 8 M urea without lipid (red dashed).

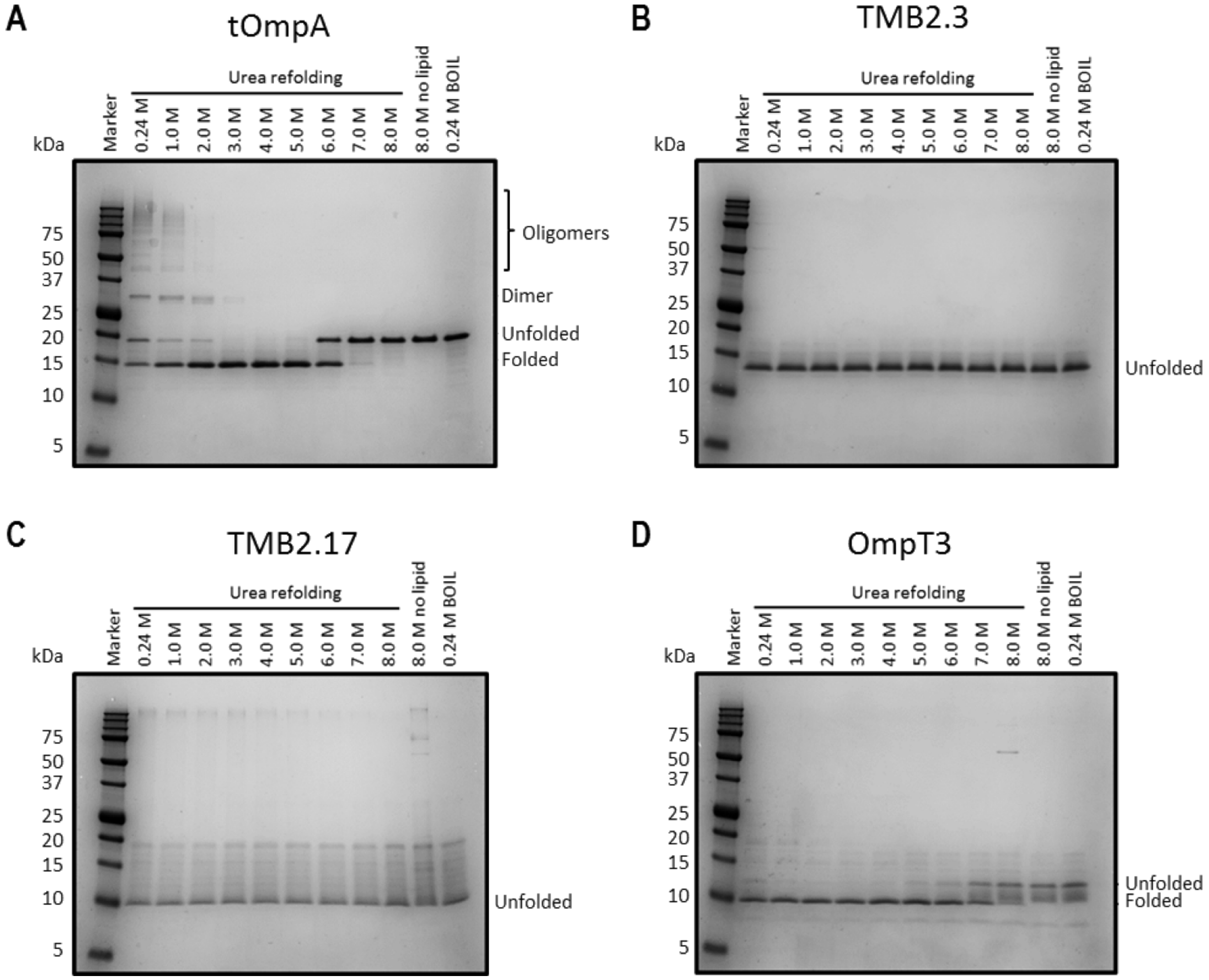


**Fig. S21. SDS-PAGE band-shift folding assays.**

(**A**) tOmpA, (**B**) TMB2.3, (**C**) TMB2.17 and (**D**) Omp*Trans*3 were refolded overnight at 2 °C in DUPC LUVs at a lipid-to-protein ratio (LPR) of 600:1 in 50 mM glycine-NaOH pH 9.5 containing 0.24-8 M urea. Samples were run on 15% (w/v) acrylamide/bis-acrylamide (37.5:1 w/w) Tris-tricine gels to resolve folded and unfolded species. The boiled sample was heated to >95 °C for 10 minutes prior to loading.

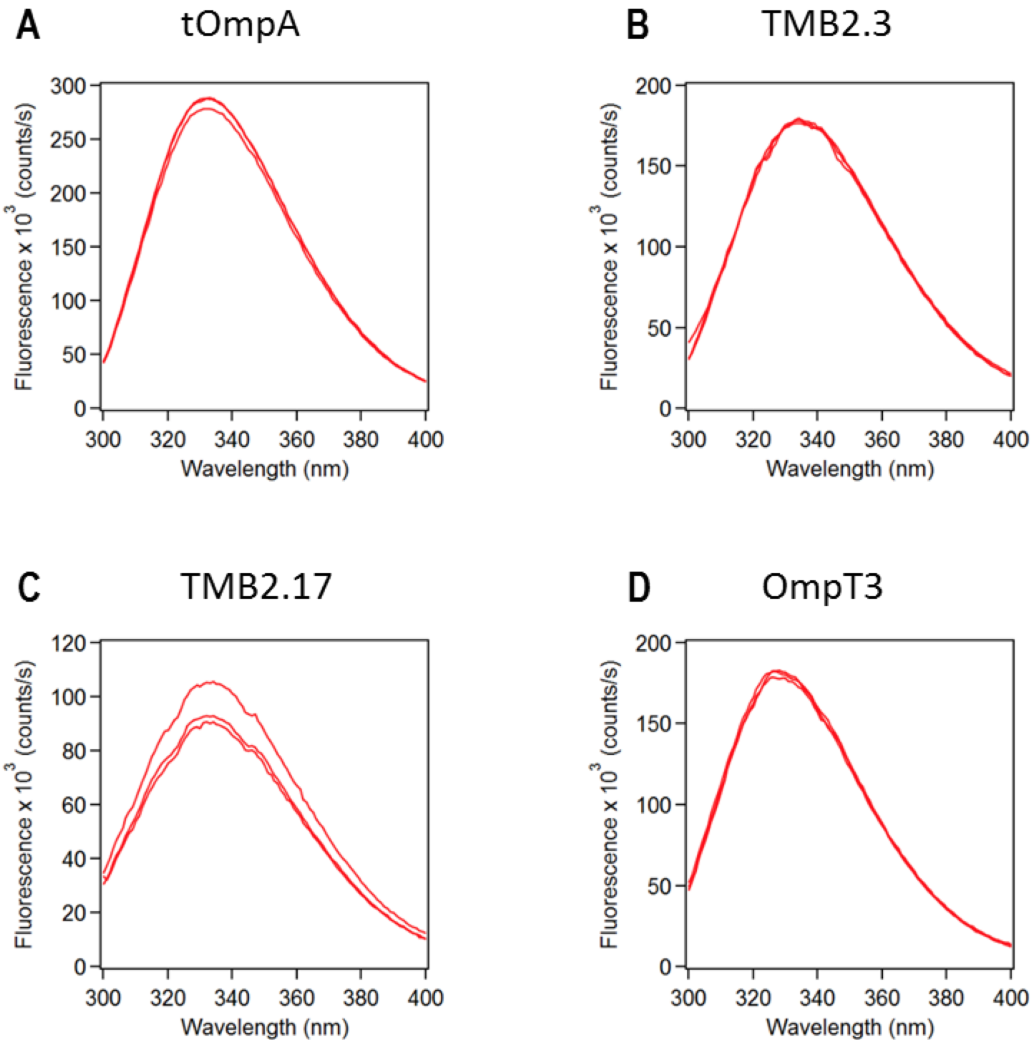


**Fig. S22. Tryptophan fluorescence emission spectra of folded OMPs.**

(**A**) tOmpA, (**B**) TMB2.3, (**C**) TMB2.17 and (**D**) Omp*Trans*3 folded after 30 minutes at 25°C in DUPC LUVs at an LPR of 3200:1 (mol/mol) in 50 mM glycine-NaOH pH 9.5 containing 2 M urea. The spectra show a fluorescence maximum at 335 nm indicative of the folded state. Three replicates are shown for each.

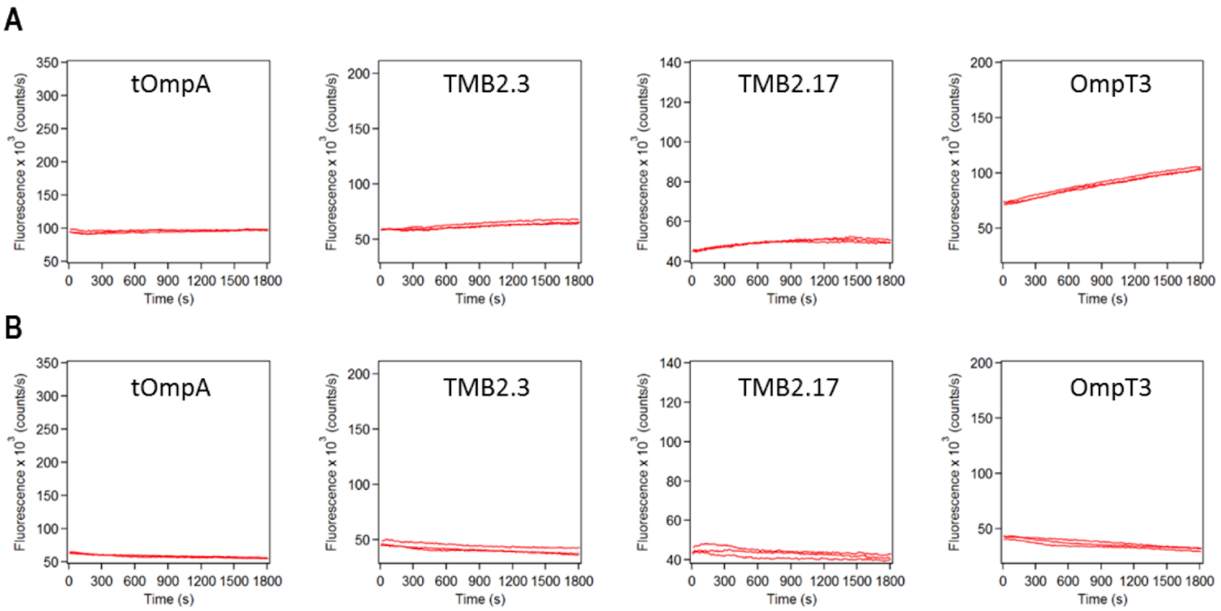


**Fig. S23. Designed TMBs are unable to fold in 9 M urea or without lipids.**

Kinetics of TMB folding were monitored by tryptophan fluorescence emission intensity at 335 nm. (**A**) OMPs were diluted into DUPC LUVs at an LPR of 3200:1 (mol/mol) in 50 mM glycine-NaOH pH 9.5 in 9 M urea at 25°C, and (**B**) TMBs were diluted in 50 mM glycine-NaOH pH 9.5 in 2 M urea at 25 °C in the absence of lipid. TMBs show no folding in 9 M urea over the timescales investigated (30 minutes), with the exception of Omp*Trans*3 which folds with slow kinetics under these conditions. These TMBs do not fold in 2 M urea in the absence of lipids.

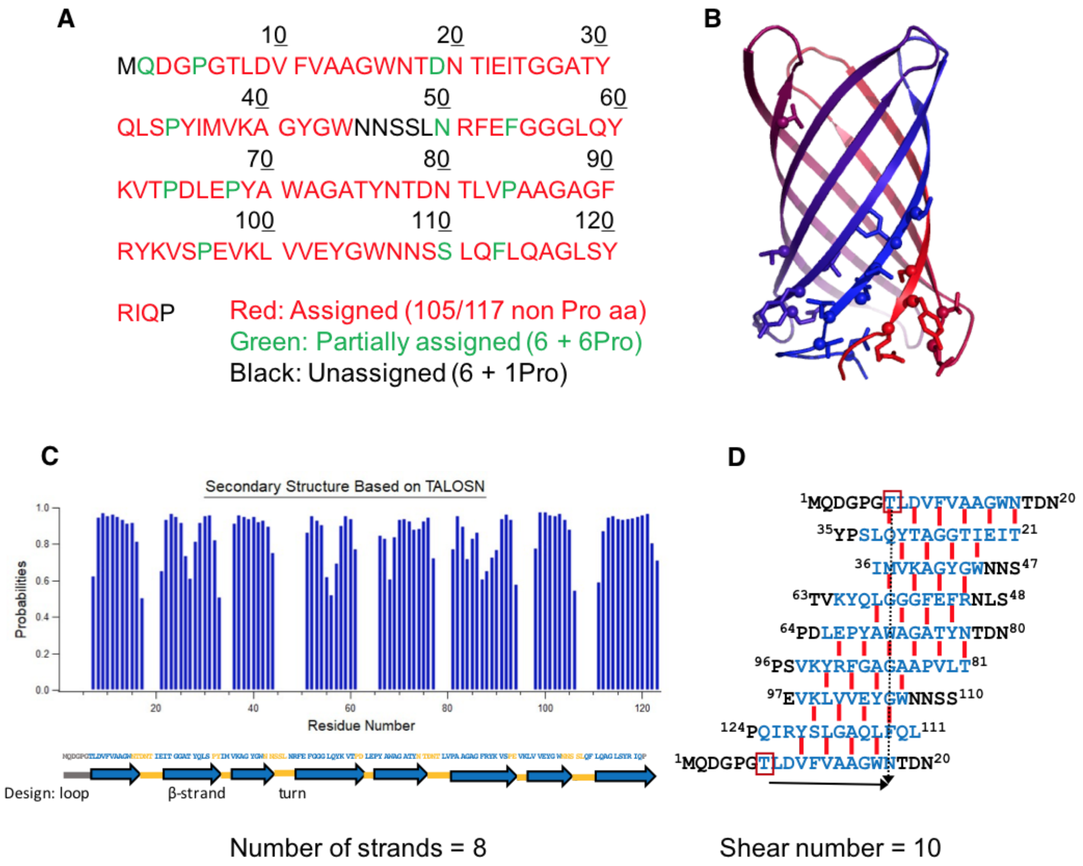


**Fig. S24. NMR spectrometry results validate the number of strands and the shear number of the design TMB2.3**.

(**A**) Coverage of the peak assignments mapped on the sequence of TMB2.3. (**B**) Residues showing multiple resonance peaks in the NMR experiment mapped onto the 3D model of the TMB2.3 design. (**C**) Secondary structures predicted based on secondary chemical shifts using TALOS-N and mapped on the TMB2.3 sequence. The pictogram in the bottom of the figure and the color show the secondary structure properties in the design model (blue shows β-strands and yellow β-turns). (**D**) Secondary structure NMR predictions and NOEs mapped on the sequence of TMB2.3.

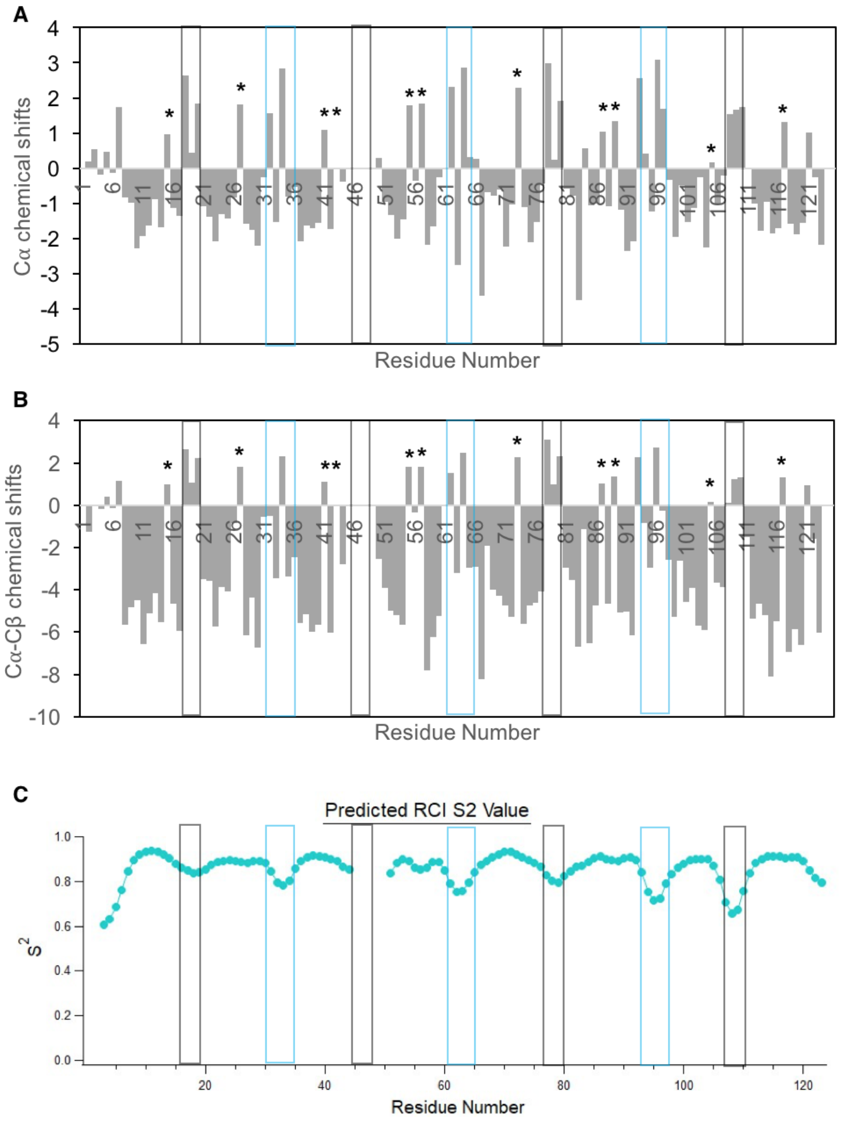


**Fig. S25. Per residue chemical shifts and Random Coil Index (RCI S2) derived from the NMR profile of TMB2.3 in DPC detergent micelles.**

The positions of glycine kink residues are marked with stars. The *cis* β-turns are highlighted by blue boxes (including the associated β-bulge residue) and the *trans* β-turns are highlighted by black boxes. (**A**) Cα chemical shifts of the assigned residues in the β-barrel. (**B**) Cα-Cβ chemical shifts of the assigned residues in the β-barrel. For glycine residues, the Cβ chemical shifts are set to 0. (**C**) Random coil index predicted with the TALOS-N software based on the chemical shifts.

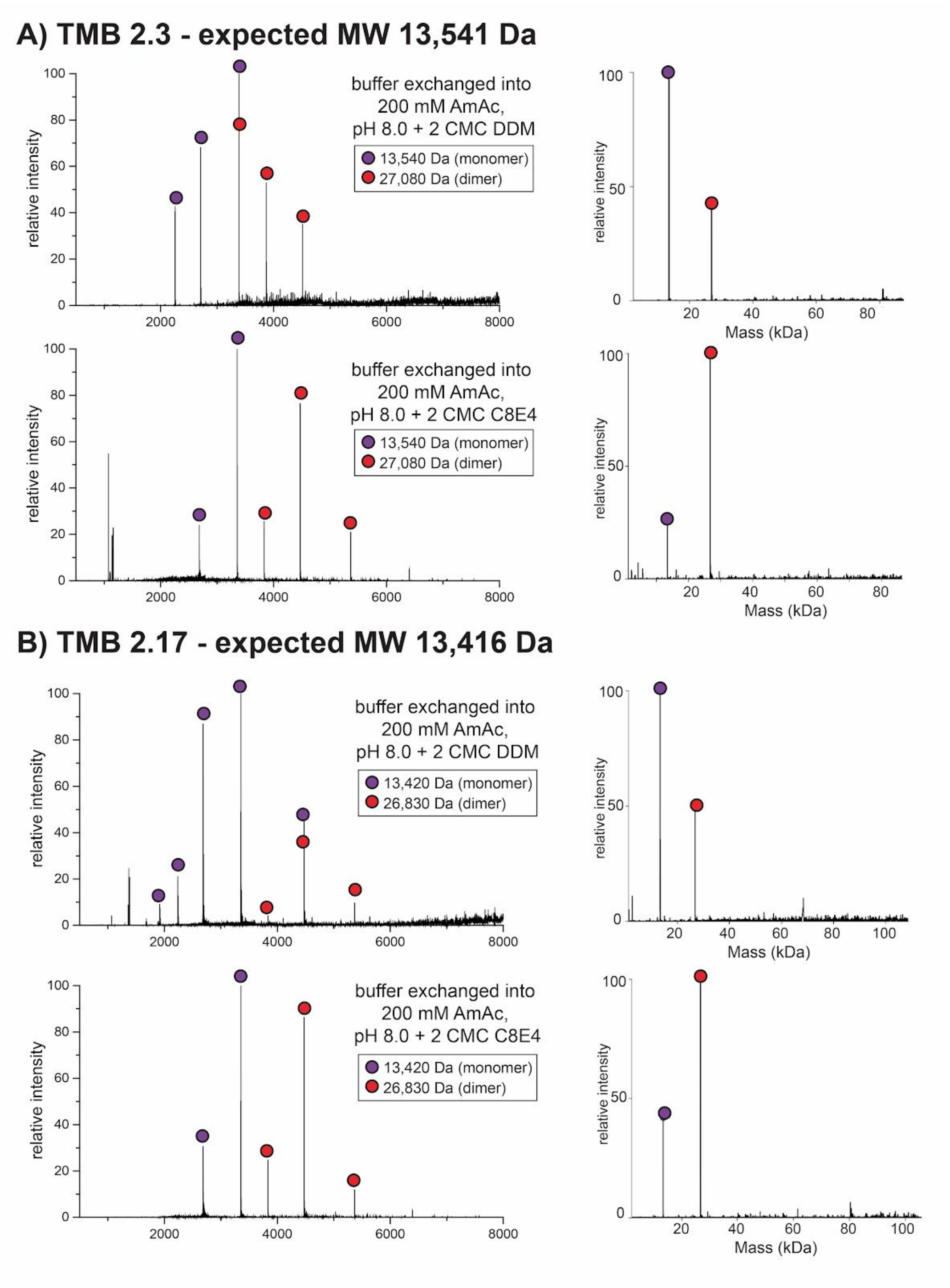


**Fig. S26. nMS profiles of TMB2.3 and TMB2.17 show monomeric and dimeric species.**

TMB2.3 (**A**) and TMB2.17 (**B**) were refolded in a buffer with 2X CMC DPC and the folded TMBs were purified by SEC. The TMBs were then buffer-exchanged into detergents compatible with nMS (DDM in top panel and $C_8E_4$ in bottom panel). The nMS analysis was made on at least two independent protein samples for each design. Monomeric and dimeric species were present each time at different ratios.

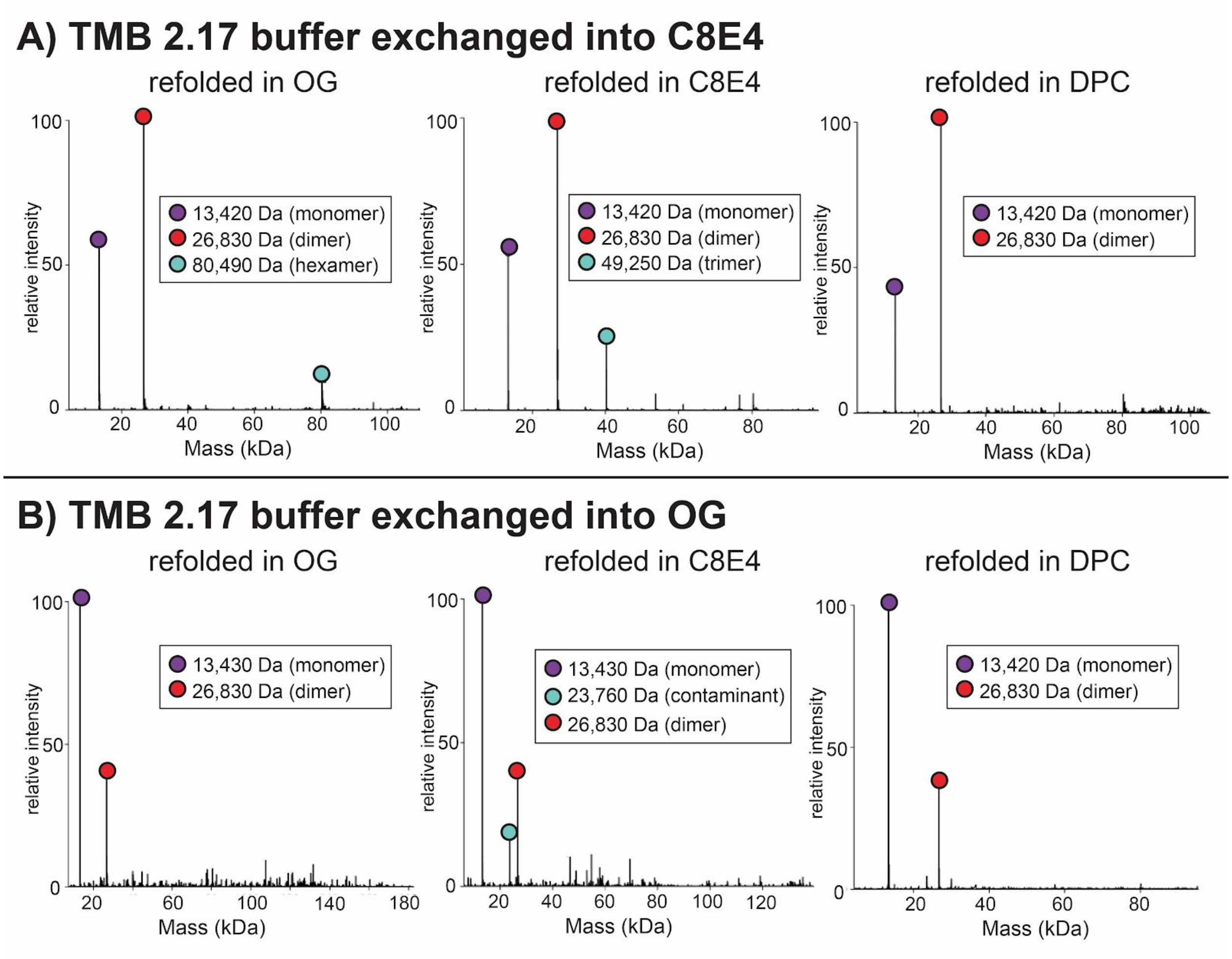


**Fig. S27. The ratio of monomer to dimer of TMB2.17 depends on the sample conditions rather than the refolding conditions.**

TMB2.17 was refolded in parallel into three different buffers containing 2X CMC of OG, $C_8E_4$ or DPC. The refolded proteins were purified by SEC and buffer-exchanged into either $C_8E_4$ (**A**) or OG (**B**) before nMS. The ratio of monomer to dimer detected by nMS depends on the sample conditions, suggesting that the monomer and the higher order species are in equilibrium and that the higher order species do not form larger β-barrels.

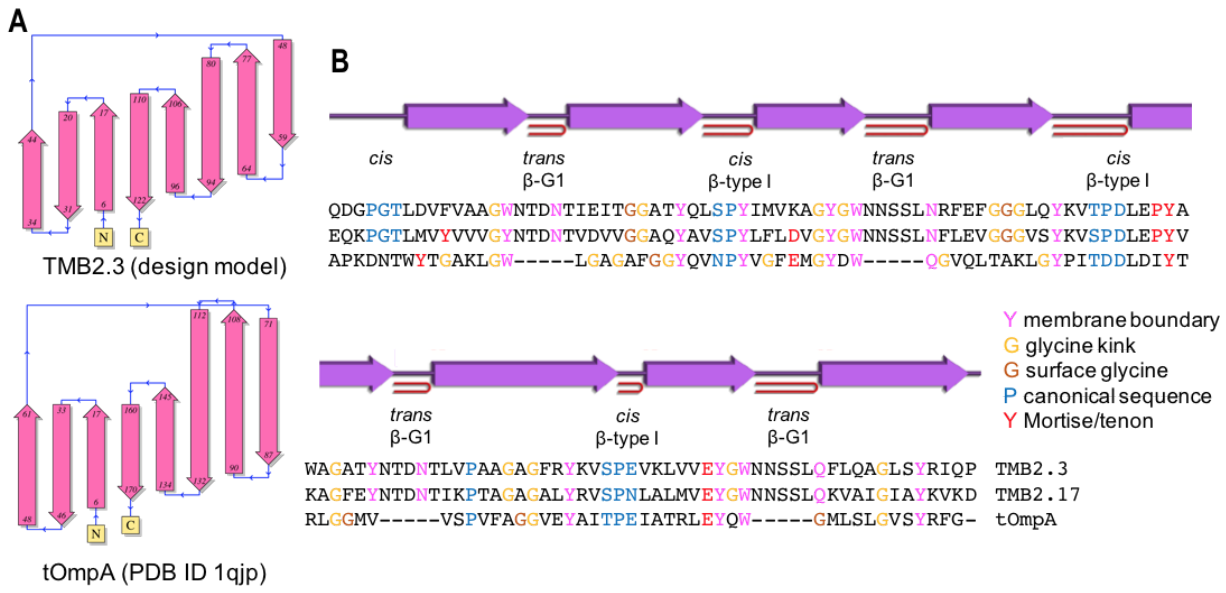


**Figure S28: Comparison of the architecture and sequences of the TMB2.3 and TMB2.17 designs to native tOmpA.**

(**A**) Comparison of the topology diagrams generated with PDBsum (*98*) with the Rosetta model TMB2.3 and the crystal structure of the native tOmpA. (**B**) Alignments of TMB2.3, TMB2.17 and tOmpA sequences mapped to the secondary structure to TMB2.3. The *cis* loops of tOmpA have been truncated to facilitate the graphical representation. Special positions in the sequences are highlighted with colors (legend on the right).

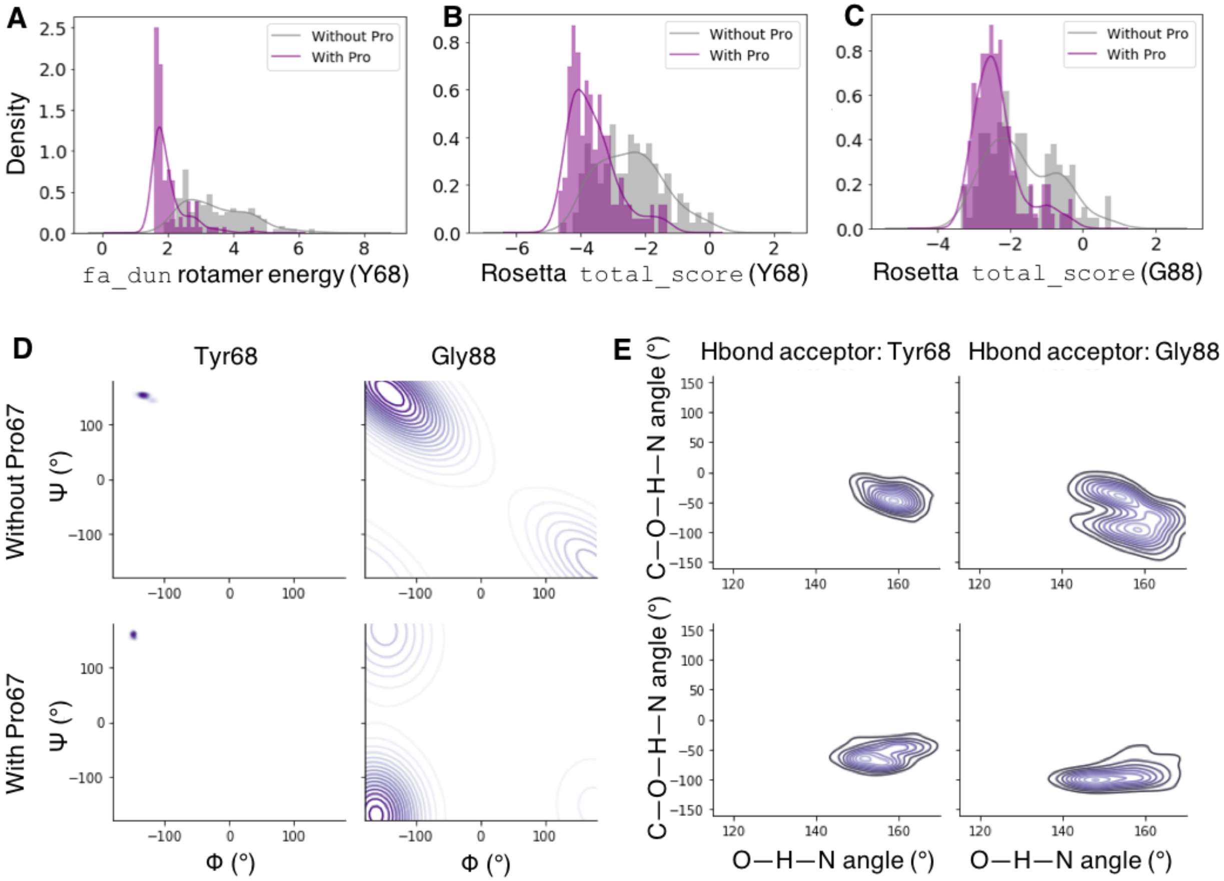


**Fig. S29. TMB backbones relaxed with proline at position 67 preceding the Tyr68 of a mortise/tenon motif stabilizes the aromatic rescue conformation.**

(**A**) The tyrosine rotamer characteristic of the aromatic rescue interaction is more favorable (lower `fa_dun` energy) in the presence of Pro67. (**B**,**C**) Tyr68 (**B**) and G88 (**C**) in the mortise tenon motif have lower energy based on Rosetta's `total_score`. (**D**) The presence of Pro67 enables a more extended conformation for Gly66 glycine kink with a negative Ψ angle. (**E**) The presence of Pro67 enables a more extended conformation for Gly66 glycine kink with more pronounced out-of-plane backbone hydrogen bonds.

| Number of strands (n) | Shear number (S) | R (Å) | $\theta$ (°) | Number of core Cβ-strips |
|---|---|---|---|---|
| 8 | 8 | 7.3 | 36.3 | 4 |
| 8 | 10 | 8.0 | 42.5 | 5 |
| 10 | 12 | 9.7 | 41.3 | 6 |
| 12 | 12 | 10.8 | 36.3 | 6 |
| 12 | 14 | 11.4 | 40.5 | 7 |
| 12 | 16 | 12.2 | 44.4 | 8 |
| 12 | 18 | 12.9 | 47.7 | 9 |
| 12 | 24 | 15.4 | 55.7 | 12 |
| 14 | 14 | 12.5 | 36.3 | 7 |
| 14 | 16 | 13.2 | 40.0 | 8 |
| 16 | 16 | 14.3 | 36.3 | 8 |
| 16 | 18 | 15.0 | 39.5 | 9 |
| 16 | 20 | 15.6 | 42.5 | 10 |
| 16 | 22 | 16.4 | 45.2 | 11 |
| 18 | 18 | 16.1 | 36.3 | 9 |
| 18 | 20 | 16.7 | 39.2 | 10 |
| 18 | 22 | 17.4 | 41.9 | 11 |
| 19 | 20 | 17.3 | 37.7 | 10 |
| 22 | 24 | 20.2 | 38.7 | 12 |
| 24 | 26 | 22.0 | 38.5 | 13 |
| 26 | 30 | 24.5 | 40.2 | 15 |
| 36 | 18 | 27.5 | 20.1 | 9 |
| 60 | 60 | 53.3 | 36.3 | 30 |
| 108 | 54 | 82.4 | 20.1 | 27 |

**Table S1. Number of Cβ-strips, ideal values of β-barrel circumference based on the average radius and strand staggering angle for the identified classes of β-barrels.**

The radius and strand staggering angle were calculated using equation 1 and equation 2 in the main text, which were reported in (*19*). The average distance between two Cα atoms along a βstrand is 3.3 Å and the average distance between two β-strands is 4.5 Å.

| | tOmpA | Omp*Trans*1 | Omp*Trans*2 | Omp*Trans*3 | Omp*Trans*4 | OmpAAG |
|---|---|---|---|---|---|---|
| Yield (mg/L) | 128 | 40 | 44 | 88 | 51 | 61 |

**Table S2. Cytoplasmic protein expression yield obtained for native tOmpA and designed β-turn variants.**

The expressions of the six constructs were carried out in parallel in a single experiment. The given yields were calculated after cleaning the inclusion bodies and dissolving the protein in 8 M urea.

| | |
|---|---|
| **NMR constraints** | |
| Distance constraints | |
| Total unique NOE | 81 |
| Inter-residue | |
| Sequential ($\|i-j\| = 1$) | 28 |
| Medium-range ($\|i-j\| <= 4$) | 12 |
| Long-range ($\|i-j\| >= 5$) | 41 |
| Interstrand hydrogen bonds [a] | 72 |
| Total dihedral angle restraints | 204 |
| TALOS $\phi$ | 102 |
| TALOS $\psi$ | 102 |
| **Structure statistics** | |
| Violations (mean and s.d.) [b] | |
| NOE constraints (Å) | $0.062 \pm 0.001$ |
| H-bond constraints (Å) | $0.009 \pm 0.002$ |
| Dihedral angle constraints (°) | $0.331 \pm 0.017$ |
| Deviations from idealized geometry | |
| Bond lengths (Å) | $0.001 \pm 0.000$ |
| Bond angles (°) | $0.388 \pm 0.002$ |
| Impropers (°) | $0.216 \pm 0.002$ |
| Average pairwise r.m.s. deviation (Å) [c] | |
| Backbone (β-sheet residues [d]) | $0.67 \pm 0.11$ |
| Heavy (β-sheet residues) | $1.83 \pm 0.14$ |
| Backbone (all residues) | $1.21 \pm 0.21$ |
| Heavy (all residues) | $2.14 \pm 0.17$ |

a Each H-bond is restrained with two upper limits of 2.5 and 3.5 Å for HN–O and N–O, respectively.

b No NOE and H-bond violations are greater than 0.5 Å; no dihedral angle violations are more than 5°.

c Calculated for 20 lowest energy conformers. Ramachandran map analysis: 79.7% most favored, 16.6% allowed, 3.4% generously allowed, 0.3% disallowed.

d. β-sheet residues: 7-17, 21-31, 37-44, 51-58, 66-77, 81-93, 99-106, 113-121

**Table S3: NMR and refinement statistics for TMB2.3.**

| | TMB2.17 (PDB: 6X9Z) |
|---|---|
| **Data collection** | |
| Space group | *R3 :H* |
| Cell dimensions | |
| *a*, *b*, *c* (Å) | 51.08 51.08 116.71 |
| *α*, *β*, *γ* (°) | 90, 90, 120 |
| Resolution (Å) | 41.37 - 2.05 (2.12 - 2.05)[a] |
| $R_{merge}$ | 0.270 (1.375) |
| $R_{\text{pim}}$ | 0.103 (0.519) |
| $I/\sigma(I)$ | 6.66 (1.18) |
| $CC_{1/2}$ | 0.995 (0.689) |
| Completeness (%) | 99.54 (98.13) |
| Redundancy | 7.9 (7.7) |
| **Refinement** | |
| Resolution (Å) | 41.37 - 2.05 |
| No. reflections | 7114 (733) |
| $R_{\text{work}}$ / $R_{\text{free}}$ (%) | 0.260/0.273 (0.292/0.311) |
| No. atoms | 948 |
| Protein | 917 |
| Water | 31 |
| Ramachandran<br>Favored/allowed<br>Outlier (%) | <br>97.41/2.59<br>00.00 |
| R.m.s. deviations | |
| Bond lengths (Å) | 0.003 |
| Bond angles (°) | 0.64 |
| $B_{\text{factors}}$ (Å$^2$) | |
| Protein | 36.24 |
| Water | 37.44 |

[a]Values in parentheses are for the highest-resolution shell.

**Table S4: Crystallographic Data Collection and Refinement Statistics for TMB2.17 crystal structure.**

**Data S1. (sequences_designs.xlsx)**
Amino acid and DNA sequences of the designs tested in this study. The BLASTE-values against the non-redundant protein database as of 01/2020 are provided.

**Data S2. (designs_TMB_expression.xlsx)**
Expression and solubility in detergents micelles of the 135 designed sequences characterized in this study. The solubility was tested in 2X CMC of DDM detergent for the designs TMB2.1 to TB2.20, as well as the corresponding constructs with long *trans* loops of tOmpA. The remaining designs from the TMB2 set were tested for solubility in 2X CMC DPC detergent.

**Data S3. (TMB_native_beta_turns.xlsx)**
Short β-turns of 2 to 5 residues occurring at the *cis* and the *trans* sides of natural TMBs. ABEGO types correspond to the Ramachandran plot binning nomenclature used in Rosetta (67). The five 3:5 type I β-turns with a G1 β-bulge grafted into tOmpA are highlighted with black boxes. Omp*Trans*3 contains both β-turns NSS and TDN.

**Data S4. (designs_TMB_folding_screen.xlsx)**
Experimental screening result of designs from the set TMB2 in DDM, DPC and OG detergent micelles. After dissolving the protein pellets in urea and refolding the protein in 2X CMC of detergent, the major species was purified on SEC. The retention volume of the major species is provided if the peak appears later than the void volume. The purified major species was analyzed by far-UV CD and annotated as Alpha or Beta based on its CD spectrum. The temperature was then ramped up in the CD experiment to 95°C - the CD thermal melt column indicates whether the temperature increase resulted in a decrease of the β-sheet content of the protein (beta) or by the emergence of a signal in the region characteristic of a random coil (coil). To determine the resistance of the protein to proteases, trypsin and chymotrypsin treated samples were analyzed on a SDS-PAGE gel alongside an untreated sample to detect the presence of a band of the expected size. N15 labelled proteins were re-folded again in DPC micelles for HSQC. A GOOD HSQC profile denotes well dispersed resonance peaks characteristic of a folded TMB.

**Data S5. (talos_pred.xlsx)**
TALOS-N backbone torsion angle predictions based on chemical shifts. The glycine kink residues (in the core of the barrel) are highlighted in orange. The surface glycine residues are highlighted in red. The 2-residues type I β-turns on the *cis* side of the barrel and the associated β-bulges are shown in blue. The 3-residues 3:5 type I β-turns in *trans*, as well as the G1 β-bulge and the flanking residues are shown in grey.

74. Website, (available at Goddard TD, Kneller DG SPARKY 3. University of California, San Francisco. Available at http://www.cgl.ucsf.edu/home/sparky/).

80. M. D. Winn, C. C. Ballard, K. D. Cowtan, E. J. Dodson, P. Emsley, P. R. Evans, R. M.

Keegan, E. B. Krissinel, A. G. W. Leslie, A. McCoy, S. J. McNicholas, G. N. Murshudov, N. S. Pannu, E. A. Potterton, H. R. Powell, R. J. Read, A. Vagin, K. S. Wilsonc, Overview of the CCP4 suite and current developments. *Acta Crystallogr. D Biol. Crystallogr.* **67**, 235–242 (2011).

92. P. Craveur, A. P. Joseph, J. Rebehmed, A. G. de Brevern, β-Bulges: extensive structural

analyses of β-sheets irregularities. *Protein Sci.* **22**, 1366–1378 (2013).
